# Global failure to track international biodiversity targets on islands

**DOI:** 10.64898/2026.10.12.758463

**Authors:** Fabio Mologni, Franz Essl, Claudine Ah-Peng, Paulo A. V. Borges, Giorgia Camperio, Rüdiger Otto, Rui B. Elias, Brent Emerson, José María Fernández-Palacios, Rosalina Gabriel, Sandra Domingues Gomes, Nathaly R. Guerrero-Ramírez, Holger Kreft, Lea de Nascimento, Jairo Patiño, Elena Rocafull, Samantha Suter, Natalia Sierra Cornejo, Patrick Weigelt, Nicholas Wilding, Delphine Clara Zemp, Bernd Lenzner

## Abstract

Islands harbour exceptional biodiversity but face severe threats, making their conservation crucial to achieving the Kunming–Montreal Global Biodiversity Framework. To evaluate progress toward its 23 targets across islands worldwide, we screened 504 biodiversity and socioeconomic indicators, and retained 47. We modelled historical time series from 1950, then projected to 2030. Progress was uneven: nine targets improved across all available indicators, while nine displayed conflicting trends. Critically, 15 of 23 targets (65%) lacked a suitable headline indicator, and five had none. Furthermore, 29 of 47 indicators (62%) were reported nationally, potentially obscuring island conditions. Most targets also lacked quantitative objectives, limiting evaluation of whether change rates suffice for 2030. Overall, progress toward halting island biodiversity loss remains insufficient and unlikely to meet the Framework targets.

## Main Text

In 2022, the Kunming-Montreal Global Biodiversity Framework (KM-GBF) was adopted by 196 states with the aim to halt and reverse global biodiversity loss by 2050. The KM-GBF is the most ambitious global biodiversity framework so far. It sets 23 action-oriented targets to be achieved by 2030, supported by a monitoring framework that guides member states in their national reporting through a consistent set of quantitative indicators (*1*, *2*). The monitoring framework combines mandatory indicators, namely headline (i.e., high-level metrics that capture overall progress towards a target) and binary indicators (i.e., qualitative metrics tracking the adoption of policy instruments, legislation, and administrative frameworks required to achieve the targets), with optional component and complementary indicators (i.e., indicators capturing specific aspects of a target), as well as national indicators (i.e., indicators tailored to country-specific contexts) (*2*). Despite substantial progress on the monitoring framework, midway through the decade, the world is not on track to achieve the targets (*3*, *4*). Twelve percent of the KM-GBF targets still lack suitable indicators, while many others remain underdeveloped (*2*), a shortfall identified as the principal constraint on tracking and course-correcting global biodiversity conservation efforts (*4*).

Globally, biodiversity is unevenly distributed, with islands harbouring approximately 20% of global biodiversity, three times their proportional share of global land area (*5*). Meanwhile, two-thirds of recorded extinctions and half of endangered species worldwide concern island biota (*5*). In addition, island biota are also particularly vulnerable to global change and anthropogenic pressures (*6*, *7*), making its conservation crucial to achieving the goals and targets of the KM-GBF. Nonetheless, only Target 6, which addresses the introduction and impact of invasive alien species, explicitly mentions islands. Furthermore, islands face specific governance challenges: Small Island Developing States (SIDS) often lack sufficient financial and institutional capacity, Sub-National Island Jurisdictions (SNIJ) are often governed by distant mainland states located in different biogeographic settings, and some islands are politically divided across multiple nations. All these challenges can hamper standardized and consistent reporting under the KM-GBF monitoring framework, and are further exacerbated by limited institutional infrastructure, inadequate legal frameworks, insufficient workforce capacity, and limiting financial resources (*8*). Consequently, despite their critical importance to global biodiversity conservation, islands are at risk of being overlooked in coordinated international monitoring efforts.

To assess progress towards the KM-GBF targets on islands, we screened 504 biodiversity and socio-economic indicators proposed within the monitoring framework and across other global biodiversity monitoring initiatives. We retained 47 indicators that were relevant for islands. These indicators were associated with the KM-GBF targets to evaluate their coverage and identify gaps. Further, we modelled their historical trends from 1950 to the present using Generalized Additive Mixed Models. Their trajectories were then projected until 2030, assuming a continuation of observed trends. We used these projections to evaluate expected progress towards the KM-GBF targets. Finally, we mapped the indicators onto the Driver-Pressure-State-Impact-Response (DPSIR) (*9*) and the Theory of Change (ToC) frameworks (*1*). DPSIR organizes biodiversity change indicators along a causal chain spanning underlying drivers and direct pressures, the state of biodiversity, impacts on people, and responses tracking policy and management actions. ToC maps biodiversity indicators to each stage of an intervention, from initial actions to their impacts, providing a complementary framework to guide coherent selection and evaluation of biodiversity indicators. Our assessment reveals uneven progress towards KM-GBF targets on islands, with persistent negative trends in key state and pressure indicators and substantial gaps in indicator coverage.

### Insufficient island indicators to track progress towards the KM-GBF

We found that nine of the 23 targets (39%) showed positive progress across all available indicators. No target showed consistent negative trends, but nine targets (39%) reported conflicting progress, with different indicators showing both positive and negative trends (Fig. 1). Coverage across DPSIR and ToC categories was highly uneven (Fig. S1), with half of the indicators focusing on policy and management actions (DPSIR Responses 47%; ToC Outputs 50%). Response indicators tracked most positively, while trends across the other categories, such as pressure on biodiversity (e.g., Fertiliser Use) and system state (e.g., Proportion of Fish Stocks within Biologically Sustainable Levels) were mixed. This misalignment between accelerating positive policy action and persistent pressures has been shown before and suggests either significant lags between actions and their effect or that responses may be insufficient, poorly targeted, or inappropriate (*9*).

**Fig. 1.**
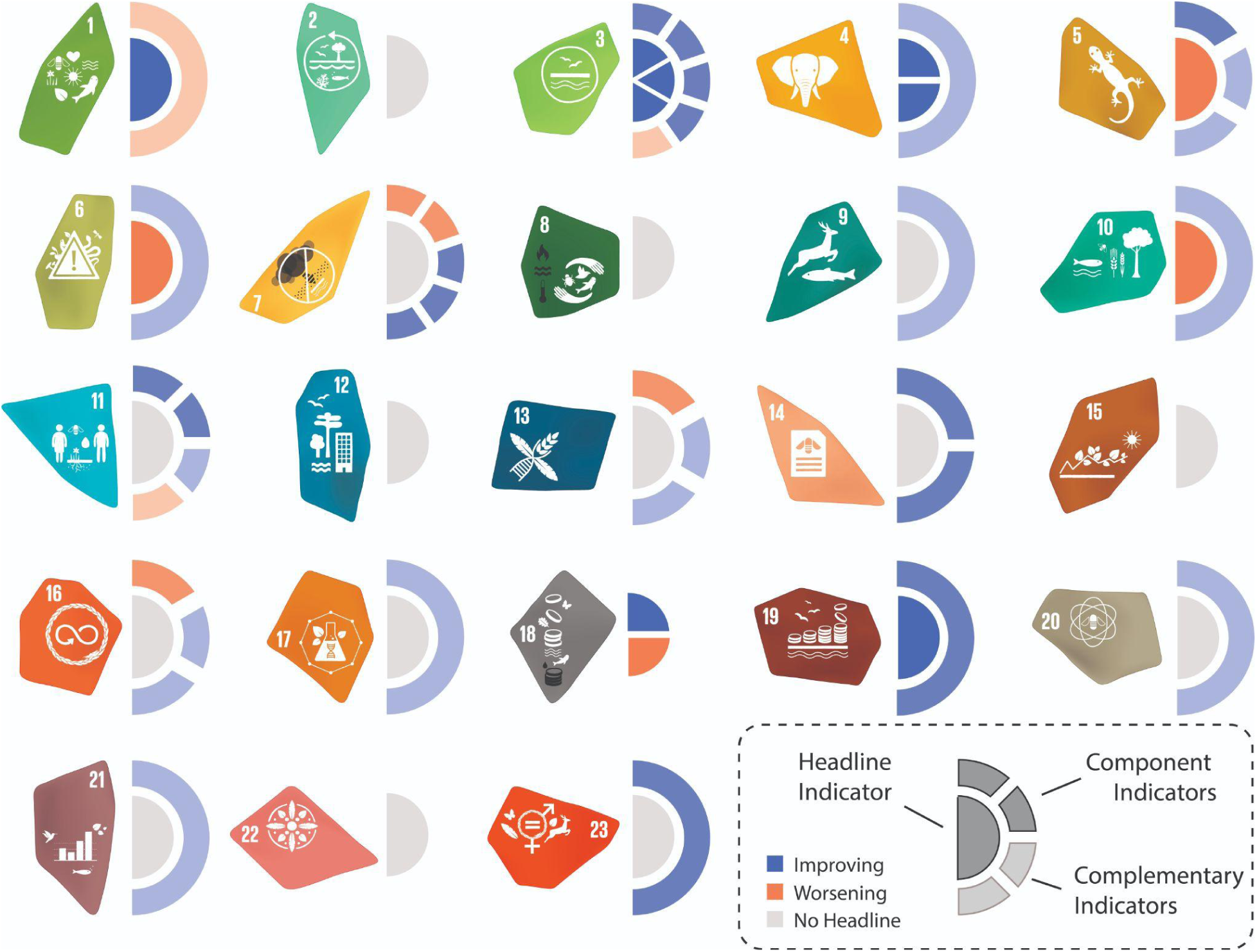
- Progress on the 23 KM-GBF targets on islands (see Table S4 for full target names). Targets are indicated by their official icons, and progress is illustrated by the half-circle diagrams. The inner half-circle represents the headline indicator(s) of the target, while the outer ring indicates the component (dark colour) and complementary (light colour) indicators. The number of slices in the inner and outer half-circles corresponds to the number of available indicators. For each indicator, we modelled its future trend until 2030 based on historically available data (see Methods and Supplement). Target progress is indicated by the colour of the slices: blue if the indicator moves towards the target (improving) and orange if it moves away from the target (worsening).

Our results further show that existing indicators provide limited capacity to track progress towards the KM-GBF targets on islands. Only eight targets (35%) were covered by a headline indicator, while ten (44%) by component or complementary indicators alone (Figs 1 and S1). Consequently, the overall coverage of targets on islands is low, with 15 targets (65%) lacking a suitable headline indicator and 5 targets (22%) not covered by any indicator at all. While earlier assessments of the overall framework found that only 30% of targets were not covered by a headline indicator (*2*), our results show this monitoring gap is substantially worse on islands.

For biodiversity threat related targets (targets 1 to 8) we found on average 3 indicators per target (Fig. 1). Five targets featured one or more headline indicators (targets 1 and 3-6). By contrast, targets 2 and 8 lacked island indicators entirely which is particularly concerning because habitat loss and degradation (target 2) and climate change impacts (target 8) pose severe threats to island biodiversity (*7*). Where data exist, progress is consistently positive only for Target 4. Notably, this target includes the only island-specific indicator identified in our searches: the Number of Island Invasive Alien Species Eradications. Despite positive progress, with successful eradications increasing and projected to continue rising, the indicator is no longer being maintained. Most targets displayed conflicting progress (1, 3, and 5–7). Indicators under Target 6 on Invasive Alien Species (IAS) showed stabilizing extinction risks driven by IAS (Supplementary Factsheets), while the rate of new IAS establishment continues to increase (Fig. 2H). A negative trend for Target 6 is particularly worrying, given that biological invasions contributed to 90% of island extinctions (*10*). This concern is compounded by limited data on IAS impacts and the use of inconsistent reporting protocols and terminology, which hinders cross-country comparison, despite the reduction of such impacts being an explicit objective of Target 6. This is in line with prior assessments, which suggest current indicators on IAS impacts do not monitor progress adequately (*11*). Target 5 on wild species showed similarly conflicting indicators. Compliance with the Convention on International Trade in Endangered Species of Wild Fauna and Flora has increased, wild species trade and threats to utilized species have declined, but the proportion of biologically sustainable fish stocks continues to fall (Fig. 2G). Target 7 on pollution follows a similar pattern: Chlorofluorocarbon emissions have fallen, cropland nutrient budgets are improving, and pollution levels are decreasing, while pesticide and fertilizer use continues to increase (Supplementary Factsheets).

**Fig. 2.**
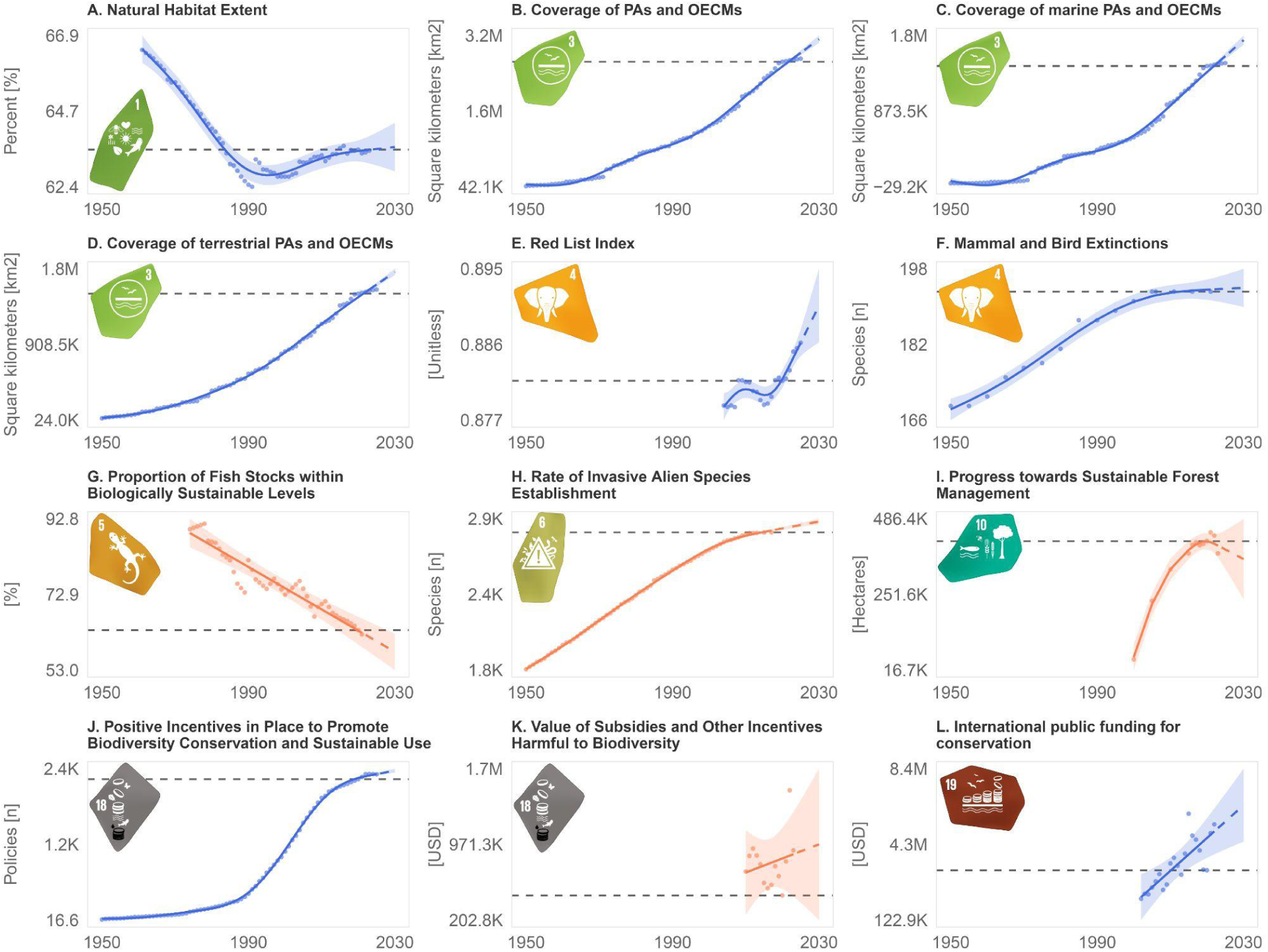
- Modelled temporal trends for the available headline indicators using generalized additive mixed models (GAMMs) grouped by target (see Table S4 for full target names). Time series run from 1950 and are projected to 2030 with some indicators only available for shorter time periods. The dark grey horizontal dashed line marks the 2020 reference year against which future KM-GBF progress is measured. Solid colored lines show historical trends, and dashed colored lines show projections. Blue indicates positive progress towards a KM-GBF target, while orange indicates negative progress. Note that positive progress may correspond to either an increasing (e.g., conservation funding) or a decreasing (e.g., habitat loss) statistical trend, depending on the direction of the specific target. For brevity, in panels B, C, and D, PAs refers to Protected Areas and OECMs refers to Other Effective Area-Based Conservation Measures, while panel L is labeled ‘International public funding for conservation’, which corresponds to the official indicator: ‘International public funding, including official development assistance (ODA) for conservation and sustainable use of biodiversity and ecosystems’.

Tracking progress toward sustainably using nature and its services (targets 9 to 13) on islands remains challenging. We found two indicators per target on average; however, only one headline across all five targets (Fig. 1 and S1). This headline indicator measures progress towards sustainable forest management (Target 10) and is declining (Fig. 2I). Negative progress compounds with the indicator’s narrow scope: it covers forests but does not capture improvements in other sectors like agriculture, aquaculture or fisheries. Conversely, fisheries biomass increased and forestry yields declined, both indicating positive progress for Targets 9 and 10. Similarly, indicators for human well-being (Target 11) show improved access to clean water and better urban air quality, yet the number of people killed, missing, or directly affected by disasters is increasing, underscoring the vulnerability of island populations to climate change. Finally, Target 13 suggests a gap between formal uptake and implementation of access and benefit-sharing (ABS) mechanisms from the use of genetic resources, with increasing Nagoya Protocol ratification, ABS reporting, and published certificates of compliance.

Tracking progress in the thematic section tools, solutions, and implementation (targets 14 to 23) is equally challenging, with about one indicator per target (Figs 1 and S1). Headline indicators remain sparse and are only available for two targets (18 and 19). Most indicators show improvements (Supplementary Factsheets). For example, more countries are submitting National Biodiversity Strategies and Action Plans (Target 14), international funding has risen (Target 19), knowledge transfer via biodiversity publications has increased (Target 21), and women’s representation in parliaments is growing (Target 23). The primary areas of concern are the growing overall ecological footprint (Target 16) and a parallel increase of both positive and harmful incentives (i.e., policy instruments, financial mechanisms) to biodiversity (Target 18). Notably, the monetary value of harmful incentives is likely underestimated, as we could account for only the fossil-fuel sector due to lack of data for other sectors (i.e., fisheries and agriculture). Finally, for two targets (15 and 22) no indicators for tracking are available. Our findings are consistent with previous overall assessments, showing a disproportionate focus on direct biodiversity measures while targets on equitable benefit-sharing, financing, and implementation tools are poorly covered (*2*). This makes these two thematic sections a critical priority for future indicator development.

To understand whether we are on track to meet the KM-GBF targets by 2030, directionality alone is insufficient; we must also assess the rate of progress. However, most targets (n=15, 65%) lack quantitative objectives, preventing such an evaluation. Where available, many benchmarks cannot be assessed at the island scale. For instance, Target 3 has both clear quantitative benchmarks and headline indicators with available data for islands. The quantitative objective of Target 3 is the flagship 30×30 target: at least 30% of terrestrial and marine areas should be protected and effectively managed by 2030. However, scaling this benchmark to the island scale might not be appropriate. Given their limited size but exceptional biodiversity, maintaining viable populations of island endemics might require protecting more than 30% of the landmass, potentially warranting higher thresholds for islands. For example, approximately 40% of the land area of the Canary Islands is already under some form of legal protection, exceeding the 30% benchmark. For the Galapagos, the proportion of land protected reaches 97% of the emerged area and as much as 133.000 km^2^ of the surrounding ocean. For SNIJs, it is also unclear whether the target should be assessed at the island or archipelago scale or at the country level, which would include the mainland. A similar argument applies to Target 19’s quantitative objective: to mobilize at least US$200 billion per year for biodiversity by 2030. Downscaling this global estimate to islands is challenging, especially for SNIJs. Hence, the lack of island-specific quantitative objectives makes it difficult to evaluate whether documented progress is occurring at a rate sufficient for reaching the KM-GBF targets by 2030. In the absence of guidance at the global scale, nation states need to be explicit on their island-specific targets within the National Biodiversity Strategy and Action Plans developed under the KM-GBF.

Over half of all available indicators (29 of 47, 62%) and headline indicators specifically (6 of 12, 50%) are reported at national scale, limiting their applicability to SNIJs. Furthermore, because of their comparatively small area and isolation, some anthropogenic pressures are more pronounced on islands. Invasive alien species (IAS) illustrate both issues. Country-level tallies overestimate IAS richness on SNIJs. For instance, the number of established IAS recorded in the Canary Islands is half of that reported for Spain. Yet IAS are one of the primary threats to island biodiversity (*7*), making their impacts disproportionately consequential on islands. Country-level estimates of natural habitat extent may likewise mask island-scale habitat loss, a critical limitation given that land-use change is a key driver of biodiversity decline on islands (*7*). These examples highlight the need for island-specific indicators and explicit disaggregation of existing indicators into their mainland and island components.

### Prioritize islands in the KM-GBF to protect global biodiversity

This first global assessment of the status and trends of biodiversity and socio-economic indicators for islands under the KM-GBF shows that biodiversity loss continues despite increasing policy and management responses. Critical pressure and state metrics show conflicting or negative trends (e.g., rising IAS establishment; declining proportions of sustainable fish stocks). At the same time, the capacity to evaluate progress remains severely constrained; 65% of targets lack a headline indicator and 22% fall short of any indicator. Furthermore, over half of the indicators lack island-level information. About two-thirds of the targets lack quantitative baselines, and existing benchmarks are difficult to apply to islands, impeding an assessment of whether progress is occurring at a sufficient rate to meet the 2030 targets.

We therefore recommend strengthening the KM-GBF monitoring framework to islands by requiring island-level reporting, especially for countries with SNIJs. We further propose adopting headline indicators that can be spatially disaggregated and reported for islands and defining clear quantitative benchmarks for targets directly related to the main threats to island biodiversity: habitat loss and degradation (Targets 1, 2, and 10), biological invasions (Target 6), and climate change (Target 8). Island-level reporting should also cover conservation efforts and biodiversity outcomes under Targets 3 and 4. Yet spatial disaggregation alone may not capture the unique characteristics of islands. We therefore propose developing island-specific indicators based on existing frameworks (*12*) and resuming data collection for key metrics, such as the number of IAS eradications on islands. More broadly, greater attention should be given to targets under the *Meeting people’s needs through sustainable use & benefits sharing* (Targets 9 to 13) and the *Tools and solutions for implementation and mainstreaming* (Targets 14 to 23) thematic sections.

To deliver this, we suggest establishing an island-focused technical expert group, modeled on the Ad Hoc Technical Expert Group on Indicators (AHTEG, *2*). Partnering with coordinated global biodiversity monitoring initiatives could support the development and implementation of relevant indicators and the supporting infrastructure for the monitoring framework. We suggest the development of an island subgroup within the global monitoring system to ensure that island-specific requirements are addressed (e.g., the Global Biodiversity Observation System, or GBioS; *12)*. Further, the establishment of coordinated monitoring networks for islands (e.g., under the umbrella of the Group on Earth Observations Biodiversity Observation Network, GEO BON), alongside the adoption of standardized measurement frameworks (e.g., Essential Biodiversity Variables, *14)* and sustained investments in capacity, training, and long-term funding specific to islands is needed.

Island-tailored consortia are already demonstrating this model. An example is the Global Island Monitoring Scheme (GIMS) (*15*), which sets up permanent 50 × 50 m plots along island elevational gradients and applies harmonised protocols across several taxa. The GIMS provides a ready-made blueprint for long-term, multi-taxon tracking of native-forest integrity across island archipelagos. A second standardized, multi-taxon model is New Zealand’s national biodiversity monitoring grid. Operating across an 8 × 8 km grid, it quantifies biodiversity data at over 1,800 permanent sample points. This grid serves as a successful long-term monitoring at a national island scale, mapped to the Essential Biodiversity Variable (EBV) standard measurement framework (*16*). Prioritizing islands within the KM-GBF is essential given their disproportionate contribution to global biological and cultural diversity (*5*). Developing suitable, disaggregated island-specific indicators to track progress is therefore a critical further step towards halting and reversing island biodiversity loss by 2030.

## Supporting information

Supplementary Factsheets

## Data availability statement

The data and code supporting this preprint are currently being prepared for deposit in a public repository and will be made available upon peer-reviewed publication. In the interim, materials are available from the corresponding author upon reasonable request.

## Acknowledgments

We thank Gilles Colling, developer of the Vectra R package, for support in extracting some of the spatial indicators. Gemini 3.1 was used for code debugging and figure formatting.

## Authors contributions

- FM: Conceptualization, Data curation, Formal analysis, Investigation, Methodology, Project administration, Resources, Software, Validation, Visualization, Writing – original draft, Writing – review & editing
- FE: Conceptualization, Funding acquisition, Methodology, Supervision, Writing – review & editing
- BL: Conceptualization, Funding acquisition, Investigation, Methodology, Project administration, Supervision, Visualization, Writing – review & editing
- CA, PAVB, GC, ROD, RBE, BE, JMFP, RG, SDG, NGR, HK, LN, JP, ER, SS, NSC, PW, NW, DCZ: Writing – review & editing

## Supplementary Materials

Materials and Methods

Figs. S1 to S6

Tables S1 to S8

References (17–30)

Supplementary Factsheets

## Supplementary material

### Materials and Methods

#### Indicators screening

We compiled a list of indicators officially adopted by the KM-GBF monitoring framework (*17*). This includes required headline (i.e., high-level metrics that capture the overall progress towards the target) and binary indicators (i.e., qualitative metrics tracking the adoption of policy instruments, legislation, and administrative frameworks required to achieve the targets), as well as optional component and complementary indicators (i.e., indicators on specific aspects of a target) (*2, 17*). The list was supplemented with indicators previously used in other global international monitoring efforts such as GEO BON and assessments of previous global policy initiatives (9). GEO BON indicators were extracted from both the GEO BON portal (*18*) and website (*19*). If no indicator from the three main sources met our inclusion criteria for a target, we opportunistically searched the KM-GBF component and complementary indicators, as well as the SDG (*20*), World Bank (*21*), OECD (*22*), and FAO (*23*) databases, for suitable indicators. In total, our list contains 504 indicators.

We screened each indicator against seven criteria, adapted from Tittensor et al. (*9*) and the Convention on Biological Diversity (*24*):

1. **Relevance** - Directly addresses at least one KM-GBF target/goal.
2. **Credibility** - Scientific or institutional.
3. **Availability** - Data and metadata are publicly available, with a minimum of five time points available.
4. **Temporal span** - Start point before 2020 and endpoint after 2020.
5. **Temporal resolution** - A gap of less than five years between updates.
6. **Spatial extent** - Data are or can be aggregated at the global scale. For country-level indicators, data are available for at least 60% of non-landlocked.
7. **Island data** - Data for islands is available.

These selection criteria are taxon-independent, encompassing both terrestrial and marine ecosystems. For more details, see Table S1. As an additional filter, complementary indicators were included exclusively if the data were already pre-processed, while headline and component indicators were computed in all cases. We also excluded national level indicators due to the global scope of our project. After screening, we retained 47 indicators that met our criteria. The full list of included and excluded indicators with reasons for exclusion can be found in Table S2 and S7, respectively.

#### Indicator alignment with targets

The alignment between indicators and targets corresponds directly to the KM-GBF categories. Headline and Binary indicators capture the overall scope, Component indicators address specific elements, and Complementary indicators provide thematic support (*2*). If an included indicator was not classified under the KM-GBF system, we conducted a qualitative assessment, assigned it to a target, and identified its alignment with that target. A justification was provided for each case (Table S3).

#### Indicator classification

The included indicators were mapped to the DPSIR (Driver–Pressure–State–Impact–Response) and ToC (Theory of Change) frameworks (*1*, *9*). The DPSIR framework serves as a functional classification linking societal activities to environmental outcomes, while the ToC framework represents the logical pathway to achieve a given target by connecting actions to their intended impacts. The DPSIR classifies indicators as drivers (i.e., activities that impact the environment), pressure (i.e., consequences and stresses that drivers put on the state of the environment), state (i.e., condition of the environment), impact (i.e., the effects resulting from changes in environmental quality), and response (i.e., responses by government and society). The ToC framework further categorises responses into five stages: inputs (i.e., resources invested into a system), process (i.e., actions or mechanisms implemented to drive change), outputs (i.e., quantification of process results), outcome (i.e., changes resulting from the intervention), and impact (i.e., changes in the condition or state of the system).

#### Spatial coverage

Indicators were organised by level of disaggregation. Most indicators were either spatially explicit or could be disaggregated to the country level. In the latter case, we categorized countries from the UN list as landlocked (lacking a seacoast), coastal (territory has a seacoast), and island states (territory consists only of islands). Landlocked countries were removed from analyses.

#### Statistical analyses

The included indicators were then modelled and projected to 2030 by using historic time series spanning from 1950 to the present (Fig. S3 and Supplementary Factsheets). We used the projected trends to evaluate progress towards the KM-GBF targets (Supplementary Factsheets). Despite calls for more numerically quantifiable targets during the formulation of the KM-GBF, many targets still lack such definitions. Hence, we project future trends based on historical data to evaluate their directionality toward the respective target. Lastly, we identified gaps in the availability of KM-GBF indicators for islands.

Before modelling, we assessed whether the data were dominated by noise by calculating the Signal-to-Noise Ratio (SNR) of each series, with SNR < 1 indicating a noisy series. In case of high noise, we planned to model data using a Dynamic Linear Model (State-space model) (*9*). However, all series displayed high signal-to-noise ratios (SNR >= 1), hence, a low level of noise.

We modelled time series for indicators individually using Generalized Additive Mixed Models (GAMMs). The indicator value was modelled as a smooth function of the year, and model parameters were estimated using the Restricted Maximum Likelihood (REML) method. The time series span from the earliest available data point (capped at 1950) to the latest available data point. Time series were only modelled if at least five data points were available. Then, future trends were extrapolated up to the year 2030 using the fitted model. To account for temporal autocorrelation, we used a two-step conditional approach. First, all series were fitted with an AR(1) (Autoregressive process of order 1) correlation structure. This approach treats the current residual error (e_t_) as being dependent only on the total residual generated in the previous time step (e_t_-1). If the residual error is still high (rho >= 0.9), we refit the model using an ARMA (0,1) structure (Moving Average process of order 1). This approach explicitly accounts for the remaining error, providing greater numerical stability.

To check the robustness of the extrapolations, we implemented a model averaging approach. We employed 14 parametric models, both linear (n = 1) and non-linear (n = 13) (see Table S5 for the full list). We modelled each time series individually. Models with >2 parameters were run only on indicators with at least 10 data points for model robustness. We selected the best models based on the small-sample corrected Akaike Information Criterion (AICc) and retained only those with a ΔAICc < 2 in the averaging process. GAMM and model averaging were compared visually (Supplementary Factsheets). To assess the sensitivity of our models to spatial coverage for country-based indicators, we modelled separately coastal and island countries (Fig. S4). To contextualize spatial coverage, we mapped the number of indicators for both country-level (Fig. S6) and spatially explicit (Fig. S7) indicators. A detailed indicator by country matrix can be found in Table S8. All analyses were conducted in R (version 4.6.0) (*25*). We used the package *mgcv* for GAMMs, *dlm* for state-space modelling, *MuMIn* for model averaging and selection, and *nlme* for non-linear models (*26–29*).

#### Data limitations and reporting constraints

Similar to prior work, we have identified recurring problems in indicator reporting hindering measurement of progress toward the KM-GBF targets and goals. These include reporting lags, inconsistent or incomplete metadata, geographic and temporal gaps, lack of FAIR compliance (non-responsive holders, gatekeeping, lack of interoperability, *2*, *6*, *10*). The geographic isolation and fragmented configuration of island systems exacerbate these issues, which compound constraints in finance and personnel, limited data infrastructure, and sparse and discontinuous monitoring networks. These issues are particularly acute in Small Island Developing States (SIDS, *6*), although the KM-GBF explicitly commits to supporting these countries with headline-indicator reporting (*1*). These limitations are documented for each indicator in the Supplementary Factsheets.

---

**Fig. S1.**
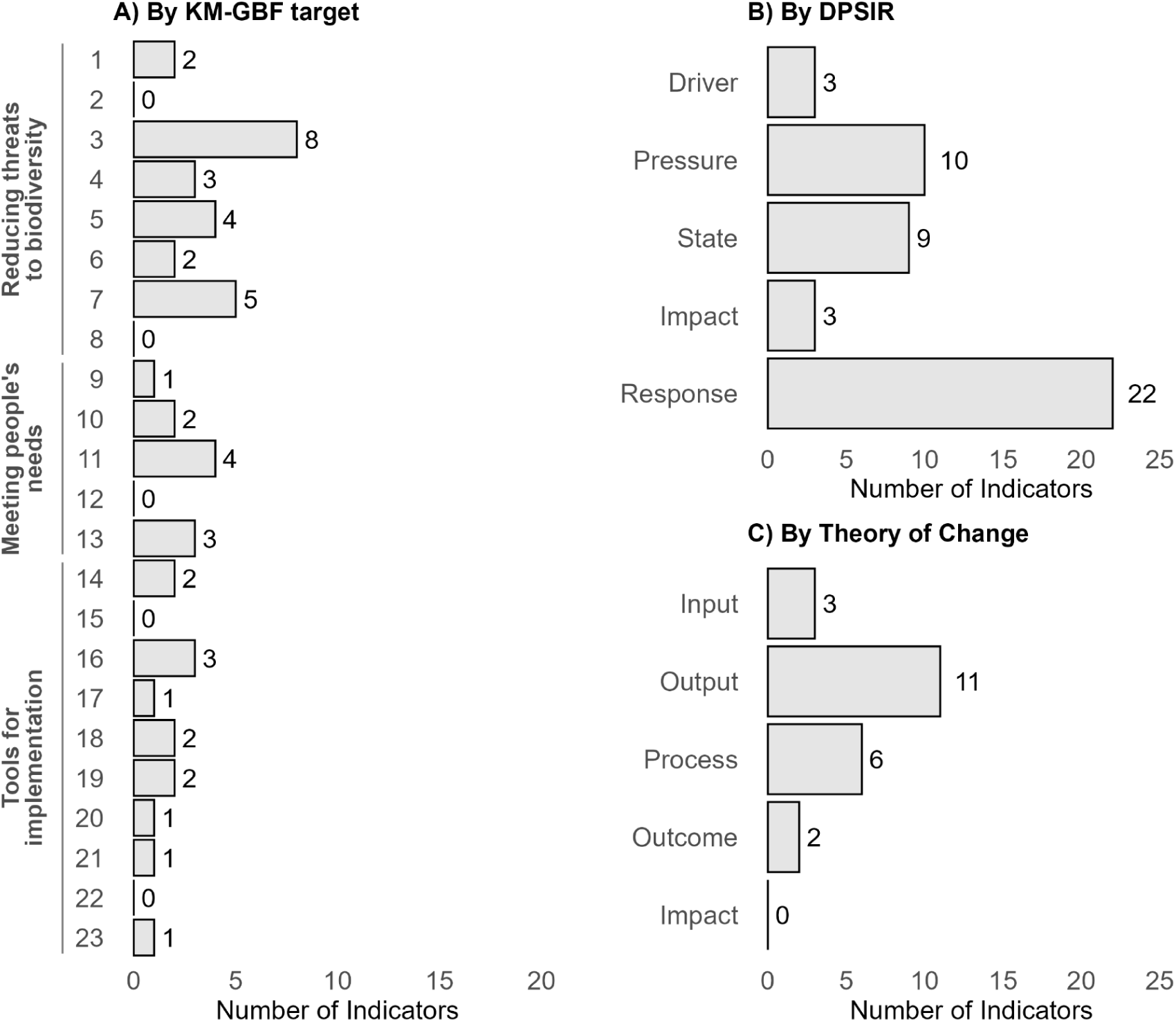
Number of indicators per target and categorised by DPSIR (Driver-Pressure-State-Impact-Response) and ToC (Theory of Change) frameworks.

**Fig. S2.**
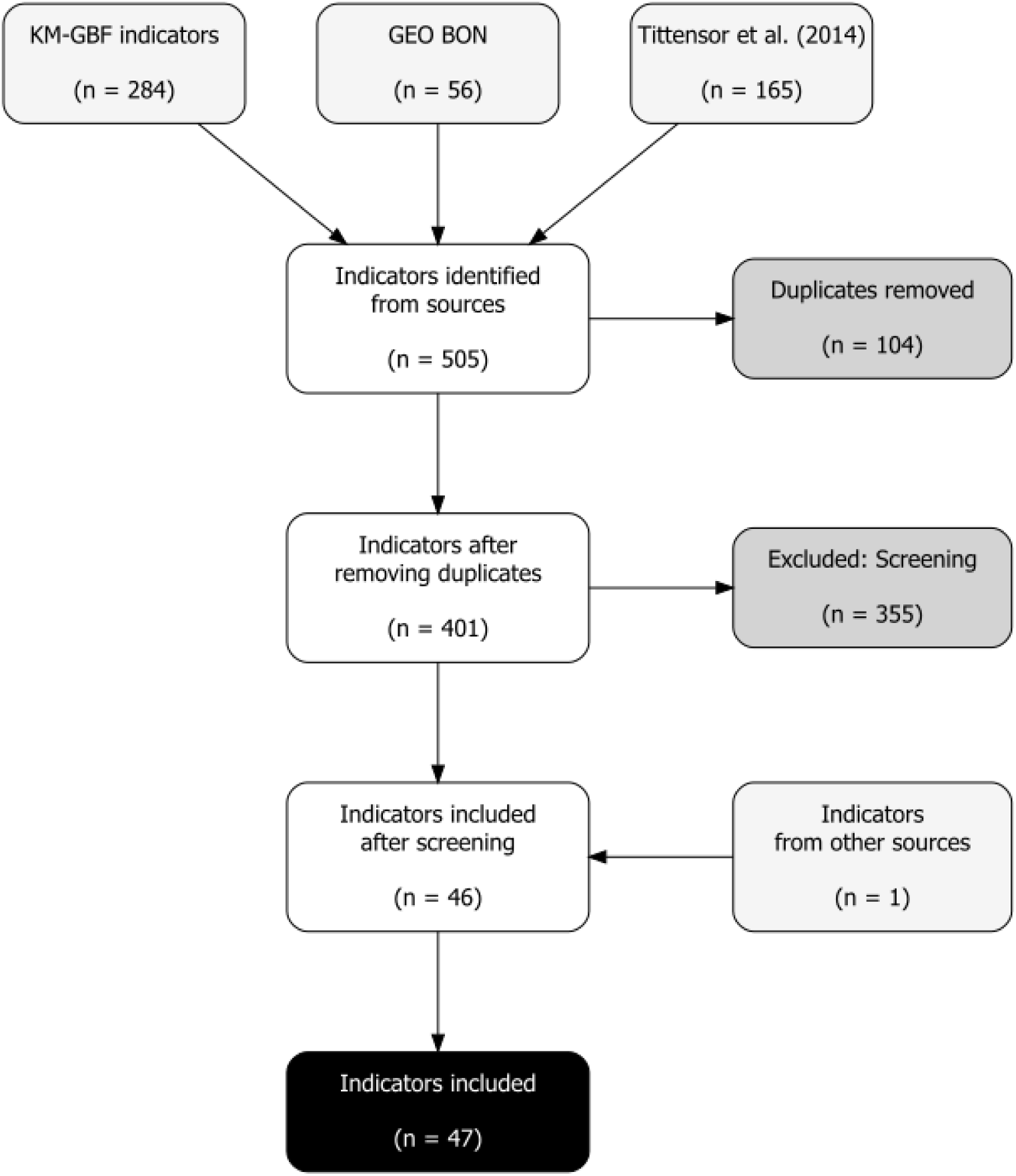
Flowchart illustrating the screening process for identifying and selecting indicators for the KM-GBF targets for islands. Box background colours indicate: indicators identified by source (light grey), duplicates removed or excluded after screening (dark grey), indicators retained at each intermediate step (white), and indicators included in the analysis (black).

**Fig. S3.**
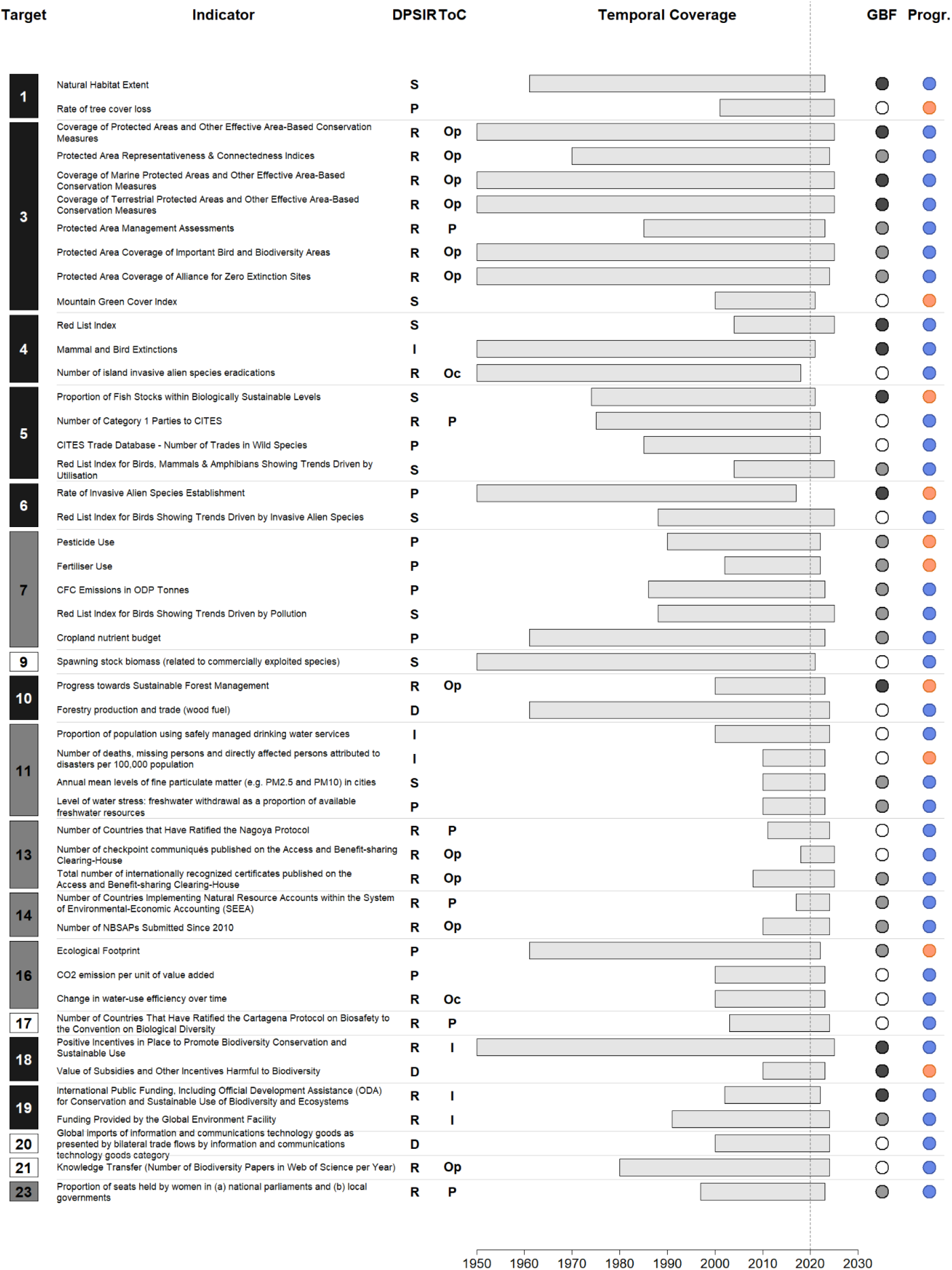
Overview of KM-GBF targets and associated indicators. The “Target” column lists all targets, with background colour indicating coverage: high (black), medium (grey), low (white). The “Indicator” column lists all indicators, followed by their classification according to the DPSIR (Driver–Pressure–State–Impact–Response) and ToC (Theory of Change) frameworks. Temporal coverage spans 1950–2025. KM-GBF (GBF) categories and progress (Progr.) by indicator are also shown. KM-GBF categories use the same colour scheme as coverage: headline and binary are in black, while component and complementary are grey and white, respectively. Progress is shown in blue (positive) and orange (negative). See methods for more information regarding the coverage categories, KM-GBF categories, and progress. Abbreviations are as follows: For the DPSIR column: D = Driver, P = Pressure, S = State, I = Impact, R = Response; for the ToC column: I = Input, Op = Output, Oc = Outcome, P = Process. No impact indicators were present under the ToC framework.

**Fig. S4.**
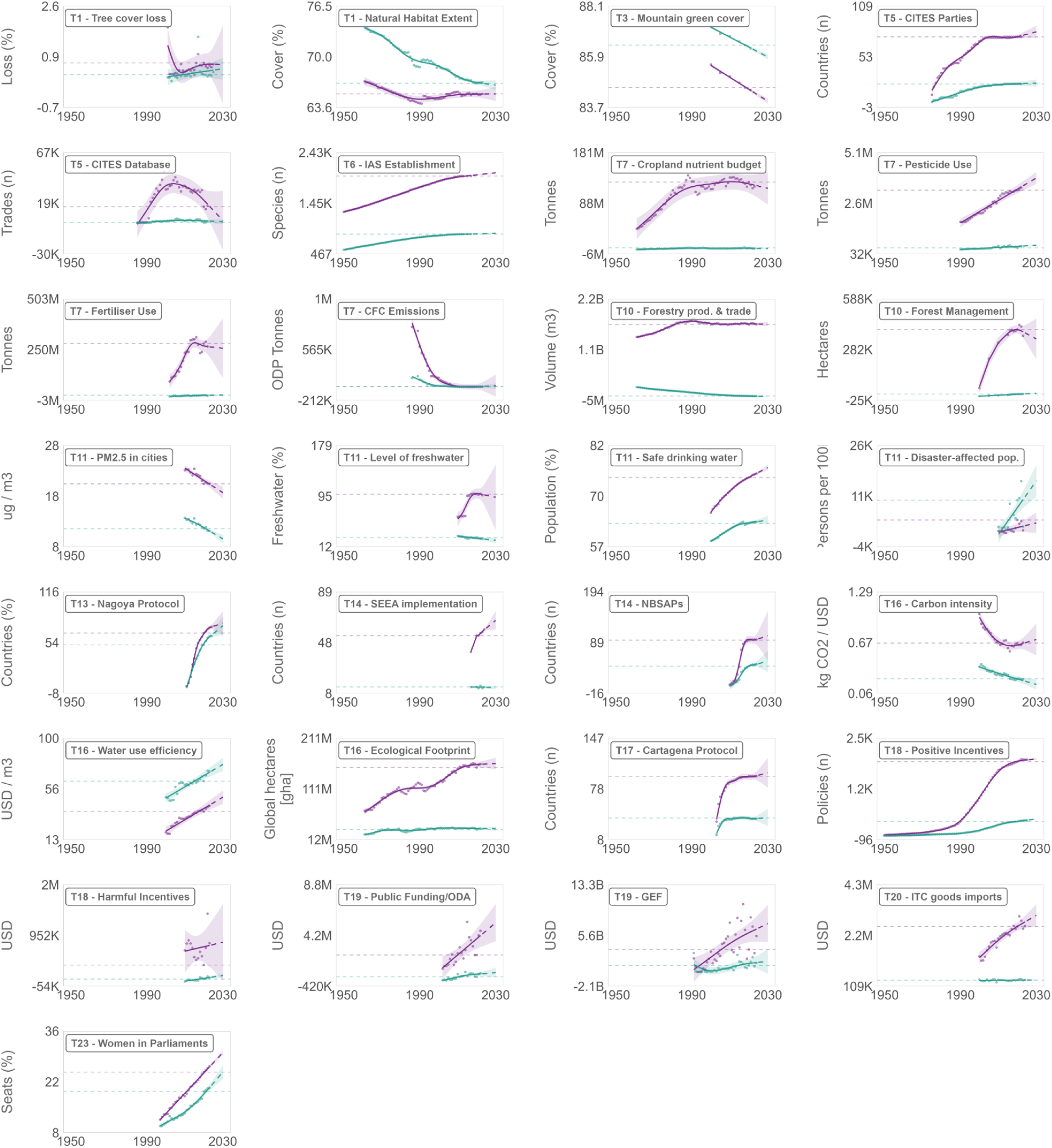
GAMM results for country-level indicators, showing trends separately for coastal (purple) and island countries (teal). Time series run from the most recent year per indicator and are projected to 2030. The horizontal dashed line marks the 2020 KM-GBF baseline for both coastal and island countries. Solid lines show historical trends, and dashed lines show projections. Top-left labels indicate the alphanumeric of the indicator and the respective KM-GBF target. Indicator names were shortened for visualization purposes as such: T1 – Tree cover loss (Rate of tree cover loss), T3 - Mountain green cover (Mountain Green Cover Index), T5 - CITES Parties (Number of Category 1 Parties to CITES), T5 - CITES Database (CITES Trade Database - Number of Trades in Wild Species), T6 - IAS Establishment (Rate of Invasive Alien Species Establishment), T7 - CFC Emissions (CFC Emissions in ODP Tonnes), T7 - Cropland nutrient budget (Cropland nutrient budget), T10 - Forest Management (Progress towards Sustainable Forest Management), T10 - Forestry prod. & trade (Forestry production and trade (wood fuel)), T11 - Safe drinking water (Proportion of population using safely managed drinking water services), T11 - Disaster-affected pop. (Number of deaths, missing persons and directly affected persons attributed to disasters per 100,000 population), T11 - PM2.5 in cities (Annual mean levels of fine particulate matter (e.g. PM2.5 and PM10) in cities), T11 - Level of freshwater (Level of water stress: freshwater withdrawal as a proportion of available freshwater resources), T13 - Nagoya Protocol (Number of Countries that Have Ratified the Nagoya Protocol), T14 - SEEA implementation (Number of Countries Implementing Natural Resource Accounts within the System of Environmental-Economic Accounting (SEEA)), T14 – NBSAPs (Number of NBSAPs Submitted Since 2010), T15 - Companies Reports (Number of Companies Publishing Sustainability Reports with Disclosure by Dimension, by Level of Requirement), T16 - Carbon intensity (CO2 emission per unit of value added), T16 - Water use efficiency (Change in water-use efficiency over time), T17 - Cartagena Protocol (Number of Countries That Have Ratified the Cartagena Protocol on Biosafety to the Convention on Biological Diversity), T18 - Harmful Incentives (Value of Subsidies and Other Incentives Harmful to Biodiversity), T18 - Positive Incentives (Positive Incentives in Place to Promote Biodiversity Conservation and Sustainable Use), T19 - Public Funding/ODA (International Public Funding, Including Official Development Assistance (ODA) for Conservation and Sustainable Use of Biodiversity and Ecosystems), T19 – GEF (Funding Provided by the Global Environment Facility), T20 - ITC goods imports (Global imports of information and communications technology goods as presented by bilateral trade flows by information and communications technology goods category), T23 - Women in Parliaments (Proportion of Seats Held by Women in National Parliaments).

**Fig. S5.**
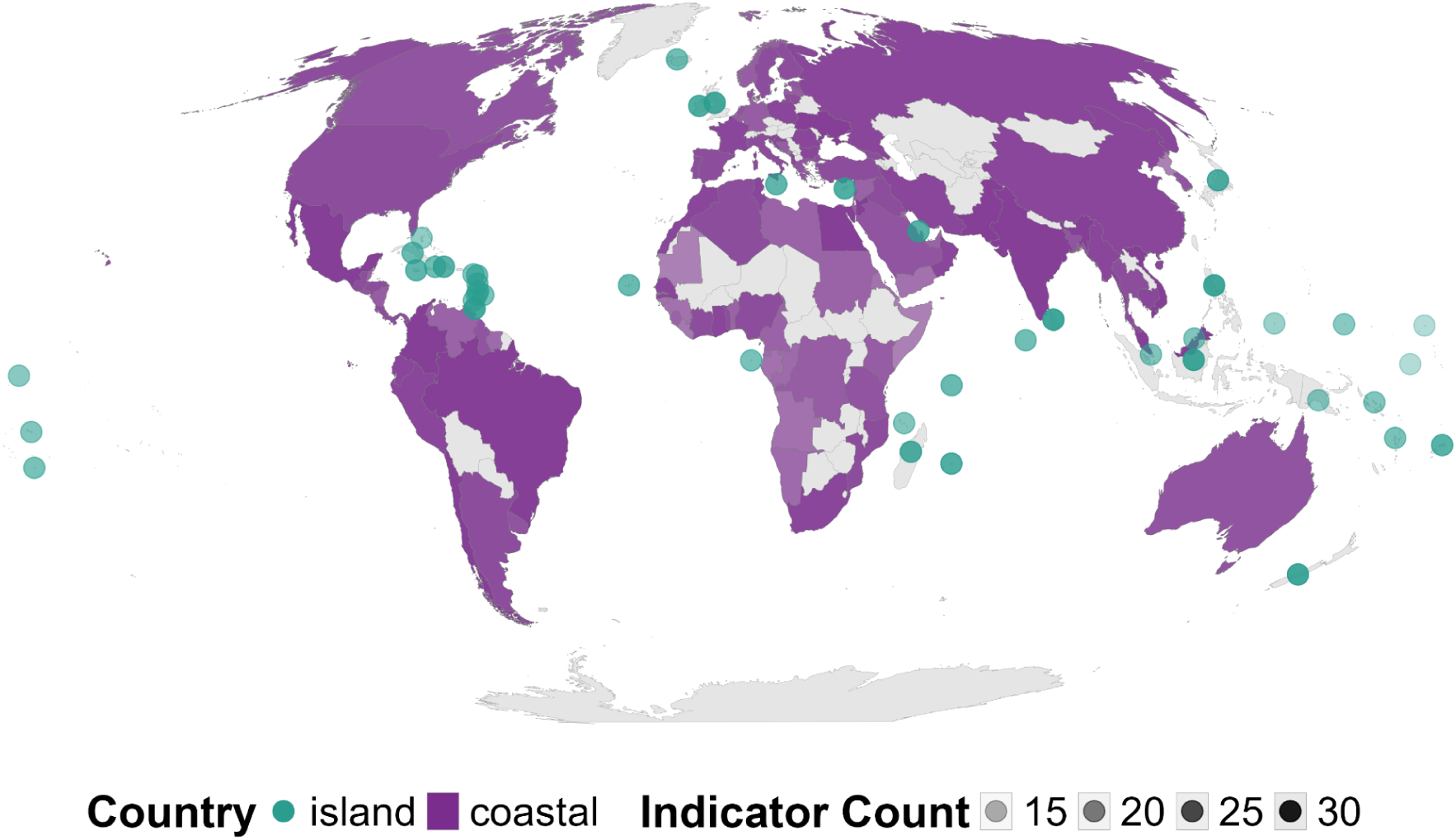
Spatial coverage of coastal (purple) and island (teal circles) countries. Island countries are depicted as circular points to ensure visibility at the global scale. Colour shading displays the number of indicators available per country, with darker colours representing a higher number of indicators.

**Fig. S6.**
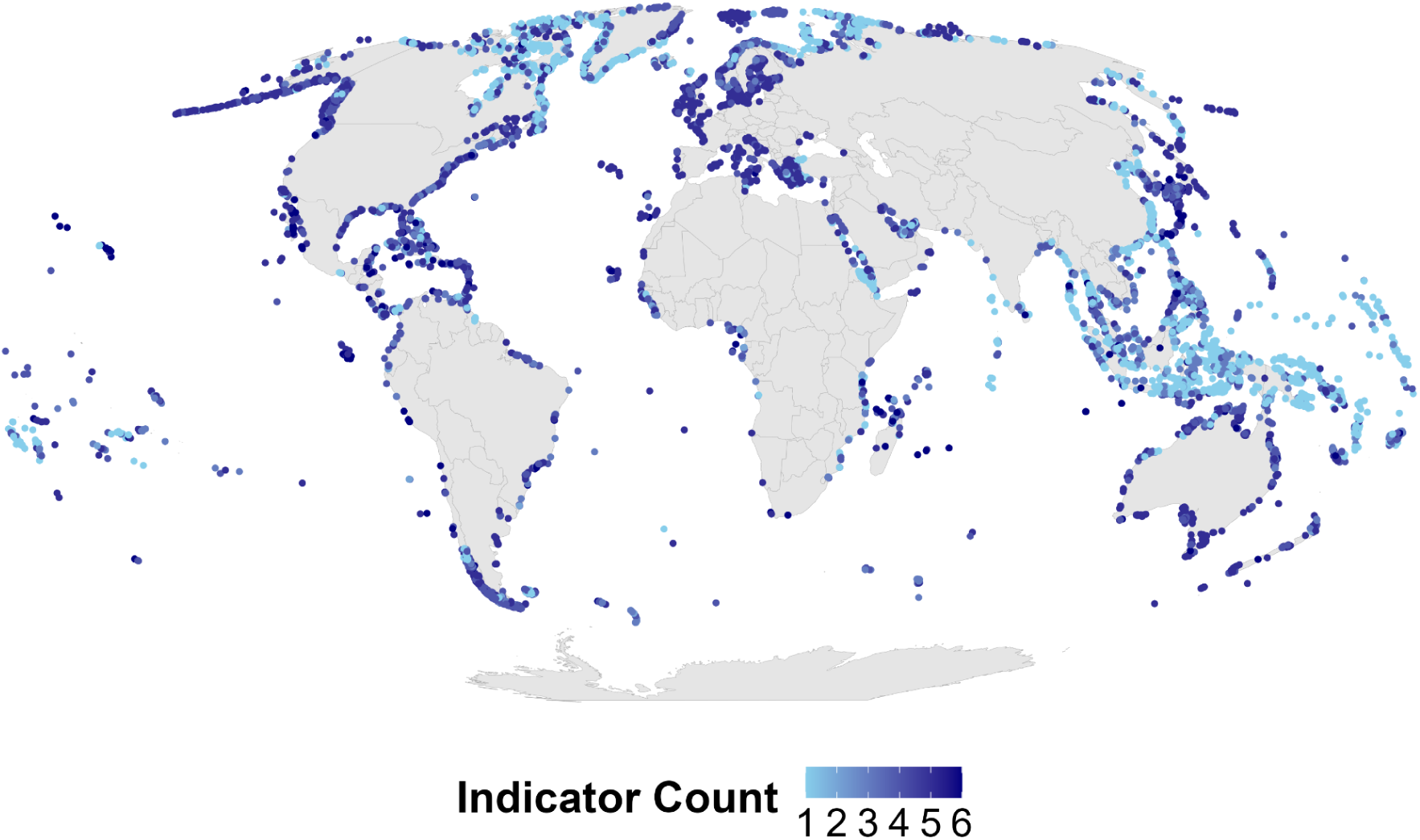
Spatial coverage for spatially explicit indicators, displaying the number of indicators available per island, with darker colors indicating a higher number of indicators. Red List Index (RLI) indicators (n = 4) are not represented, as RLIs are calculated globally and all islands overlap with at least one species range.

**Table S1.**
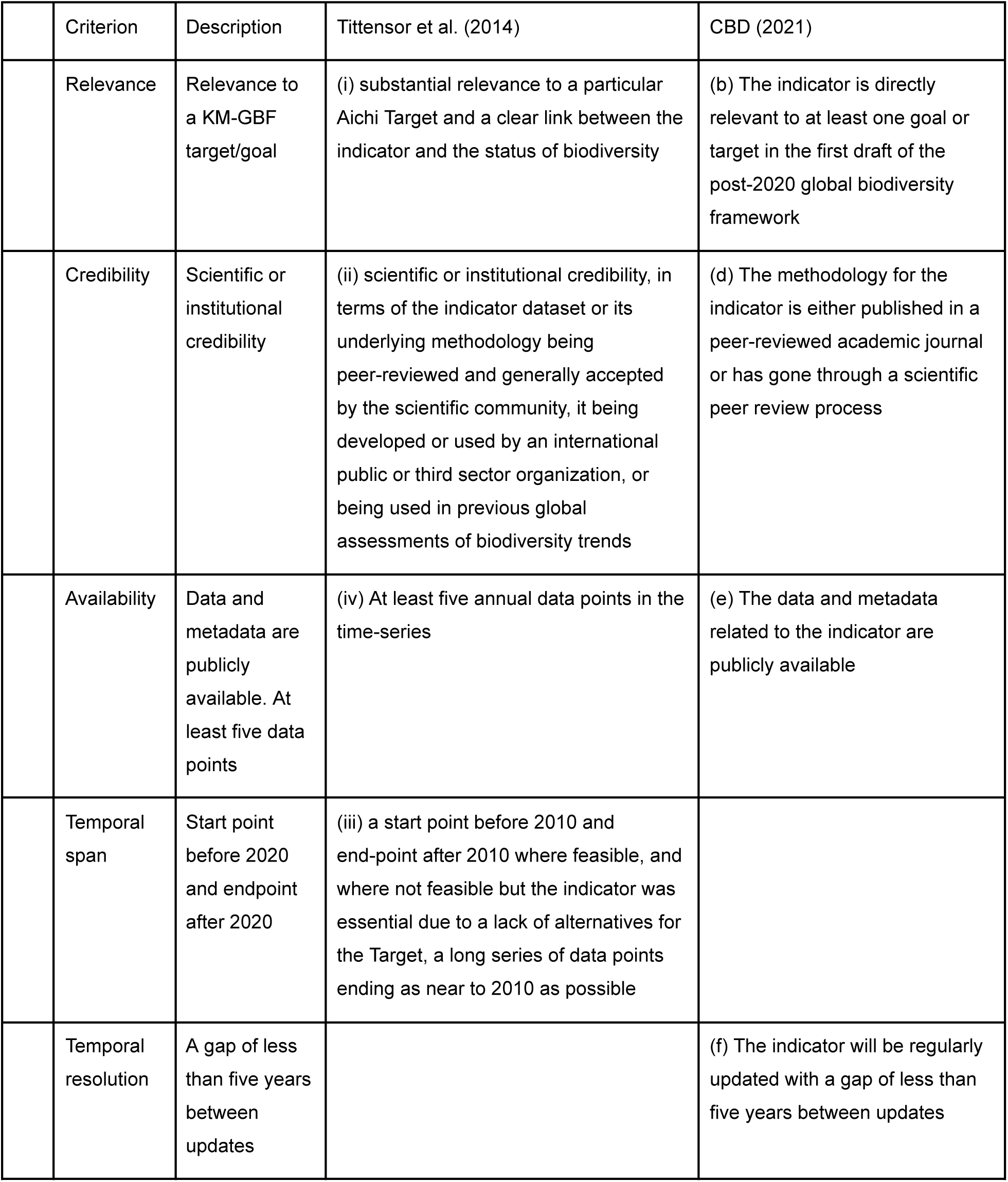
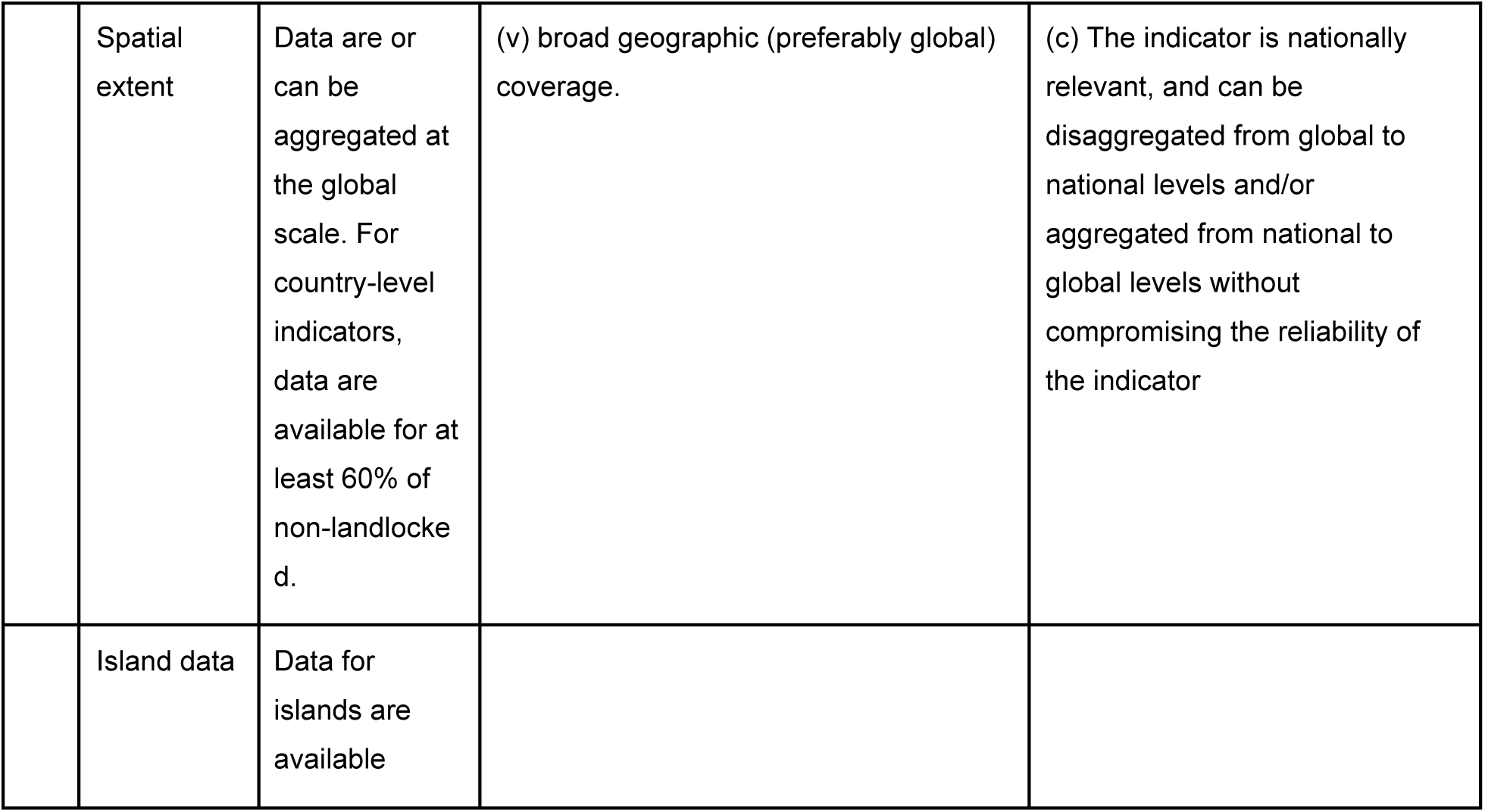
Selection criteria for biodiversity indicators, with description and match with Tittensor et al. (2014) and CBD (2021)’s criteria. Discrepancies from Tittensor et al. (2014) and CBD (2021) are indicated with grey shading.

|  | Criterion | Description | Tittensor et al. (2014) | CBD (2021) |
|---|---|---|---|---|
|  | Relevance | Relevance to a KM-GBF target/goal | (i) substantial relevance to a particular Aichi Target and a clear link between the indicator and the status of biodiversity | (b) The indicator is directly relevant to at least one goal or target in the first draft of the post-2020 global biodiversity framework |
|  | Credibility | Scientific or institutional credibility | (ii) scientific or institutional credibility, in terms of the indicator dataset or its underlying methodology being peer-reviewed and generally accepted by the scientific community, it being developed or used by an international public or third sector organization, or being used in previous global assessments of biodiversity trends | (d) The methodology for the indicator is either published in a peer-reviewed academic journal or has gone through a scientific peer review process |
|  | Availability | Data and metadata are publicly available. At least five data points | (iv) At least five annual data points in the time-series | (e) The data and metadata related to the indicator are publicly available |
|  | Temporal span | Start point before 2020 and endpoint after 2020 | (iii) a start point before 2010 and end-point after 2010 where feasible, and where not feasible but the indicator was essential due to a lack of alternatives for the Target, a long series of data points ending as near to 2010 as possible |  |
|  | Temporal resolution | A gap of less than five years between updates |  | (f) The indicator will be regularly updated with a gap of less than five years between updates |
|  | Spatial extent | Data are or can be aggregated at the global scale. For country-level indicators, data are available for at least 60% of non-landlocked. | (v) broad geographic (preferably global) coverage. | (c) The indicator is nationally relevant, and can be disaggregated from global to national levels and/or aggregated from national to global levels without compromising the reliability of the indicator |
|  | Island data | Data for islands are available |  |  |

**Table S2.** List of included indicators and associated targets. For full descriptions of the indicators please refer to the respective factsheets (Supplementary Factsheets). Abbreviations: ABSCH, Access and Benefit-sharing Clearing-House; CBD, Convention on Biological Diversity; CITES, Convention on International Trade in Endangered Species of Wild Fauna and Flora; CSIRO, Commonwealth Scientific and Industrial Research Organisation; DIISE, Database of Island Invasive Species Eradications; FAO, Food and Agriculture Organization of the United Nations; GEF, Global Environment Facility; IUCN, International Union for Conservation of Nature; OECD, Organisation for Economic Co-operation and Development; SInAS, Standardised Information on Alien Species Database; UN, United Nations; UN SDGs, United Nations Sustainable Development Goals; WOS, Web of Science.

| **Indicator Name** | **Target** | **Source** |
|---|---|---|
| Natural Habitat Extent | 1 | FAO |
| Rate of tree cover loss | 1 | Global Forest Watch |
| Coverage of Marine Protected Areas and Other Effective Area-Based Conservation Measures | 3 | Protected Planet |
| Coverage of Protected Areas and Other Effective Area-Based Conservation Measures | 3 | Protected Planet |
| Coverage of Terrestrial Protected Areas and Other Effective Area-Based Conservation Measures | 3 | Protected Planet |
| Mountain Green Cover Index | 3 | UN SDGs |
| Protected Area Coverage of Alliance for Zero Extinction Sites | 3 | Bird Life International |
| Protected Area Coverage of Important Bird and Biodiversity Areas | 3 | Bird Life International |
| Protected Area Management Assessments | 3 | Protected Planet |
| Protected Area Representativeness & Connectedness Indices | 3 | CSIRO |
| Mammal and Bird Extinctions | 4 | IUCN Red List |
| Number of island invasive alien species eradications | 4 | DIISE |
| Red List Index | 4 | IUCN |
| CITES Trade Database - Number of Trades in Wild Species | 5 | CITES |
| Number of Category 1 Parties to CITES | 5 | CITES |
| Proportion of Fish Stocks within Biologically Sustainable Levels | 5 | UN SDGs |
| Red List Index for Birds, Mammals & Amphibians Showing Trends Driven by Utilisation | 5 | IUCN Red List |
| Rate of Invasive Alien Species Establishment | 6 | SInAS |
| Red List Index for Birds Showing Trends Driven by Invasive Alien Species | 6 | IUCN Red List |
| CFC Emissions in ODP Tonnes | 7 | UN |
| Cropland nutrient budget | 7 | FAO |
| Fertiliser Use | 7 | FAO |
| Pesticide Use | 7 | FAO |
| Red List Index for Birds Showing Trends Driven by Pollution | 7 | IUCN Red List |
| Spawning stock biomass (related to commercially exploited species) | 9 | RAM Legacy Dataset |
| Forestry production and trade (wood fuel) | 10 | FAO |
| Progress towards Sustainable Forest Management | 10 | UN SDGs |
| Annual mean levels of fine particulate matter (e.g. PM2.5 and PM10) in cities | 11 | UN SDGs |
| Level of water stress: freshwater withdrawal as a proportion of available freshwater resources | 11 | UN SDGs |
| Number of deaths, missing persons and directly affected persons attributed to disasters per 100,000 population | 11 | UN SDGs |
| Proportion of population using safely managed drinking water services | 11 | UN SDGs |
| Number of Countries that Have Ratified the Nagoya Protocol | 13 | CBD |
| Number of checkpoint communiqués published on the Access and Benefit-sharing Clearing-House | 13 | CBD ABSCH |
| Total number of internationally recognized certificates published on the Access and Benefit-sharing Clearing-House | 13 | CBD ABSCH |
| Number of Countries Implementing Natural Resource Accounts within the System of Environmental-Economic Accounting (SEEA) | 14 | UN |
| Number of NBSAPs Submitted Since 2010 | 14 | CBD |
| CO<sub>2</sub> emission per unit of value added | 16 | UN SDGs |
| Change in water-use efficiency over time | 16 | UN SDGs |
| Ecological Footprint | 16 | Global Footprint Network |
| Number of Countries That Have Ratified the Cartagena Protocol on Biosafety to the Convention on Biological Diversity | 17 | CBD |
| Positive Incentives in Place to Promote Biodiversity Conservation and Sustainable Use | 18 | OECD |
| Value of Subsidies and Other Incentives Harmful to Biodiversity | 18 | UN SDGs |
| Funding Provided by the Global Environment Facility | 19 | GEF |
| International Public Funding, Including Official Development Assistance (ODA) for Conservation and Sustainable Use of Biodiversity and Ecosystems | 19 | OECD |
| Global imports of information and communications technology goods as presented by bilateral trade flows by information and communications technology goods category | 20 | UN Trade Data Hub |
| Knowledge Transfer (Number of Biodiversity Papers in Web of Science per Year) | 21 | WOS |
| Proportion of seats held by women in national parliaments | 23 | World Bank |

**Table S3.** Target assignment and categorisation according to the KM-GBF of unclassified indicators. These indicators were qualitatively assessed and assigned to the corresponding categories, with rationales provided to support the assessments.

| Indicator | Target | KM-GBF category | Rationale |
|---|---|---|---|
| Natural habitat extent | 1 | Headline | Analogous to headline, calculated using a different data and method |
| Coverage of Terrestrial Protected Areas and Other Effective Area-Based Conservation Measures | 3 | Headline | Subset of Target 3 headline |
| Coverage of Marine Protected Areas and Other Effective Area-Based Conservation Measures | 3 | Headline | Subset of Target 3 headline |
| Protected area coverage of Alliance for Zero Extinction sites | 3 | Component | Analogous to “Protected area coverage of Important Bird and Biodiversity Areas” |
| Mammal and bird extinctions | 4 | Headline | Subset of Target 4 headline |
| Number of Category 1 parties to CITES | 5 | Complementary | Only the number of parties, not whether trade is sustainable |
| CITES Trade Database - Number of trades in wild species | 5 | Complementary | A measure of total trade, not whether trade is sustainable |
| Red List Index for birds, mammals & amphibians showing trends driven by utilisation | 5 | Component | Subset of Red List Index for used species (Target 5 component) |
| Pesticide use | 7 | Component | Analogous to “Fertilizer use” (Target 7 component) |
| CFC emissions (Consumption of chlorofluorocarbons in ODP tonnes) | 7 | Component | Analogous to “Fertilizer use” (Target 7 component) |
| Red List Index for birds showing trends driven by pollution | 7 | Component | Does not measure pollution levels, but its impacts |
| Number of countries ratified the Nagoya Protocol | 13 | Complementary | Measures only countries’ agreements to share benefits, not the actual sharing of benefits |
| Number of NBSAPs submitted since 2010 | 14 | Component | This is only one policy; Target 14 implies multiple policies |
| Number of countries that have ratified the Cartagena Protocol on Biosafety to the Convention on Biological Diversity | 17 | Complementary | Analogous to “Percentage of Parties to the Cartagena Protocol on Biosafety implementing the relevant provisions of the Protocol” (Target 17 complementary) |
| Knowledge transfer (number of biodiversity papers in Web of Science per year) | 21 | Complementary | Only the amount of scientific literature, not whether the material is available, not information regarding data and other types of knowledge |

**Table S4.** Goals and targets set by the Global Biodiversity Framework.

| **Goal/Target** | **Definition** |
|---|---|
| Goal A | Protect and Restore - The integrity, connectivity and resilience of all ecosystems are maintained, enhanced, or restored, substantially increasing the area of natural ecosystems by 2050; human induced extinction of known threatened species is halted, and, by 2050, the extinction rate and risk of all species are reduced tenfold and the abundance of native wild species is increased to healthy and resilient levels; the genetic diversity within populations of wild and domesticated species, is maintained, safeguarding their adaptive potential. |
| Goal B | Prosper with Nature - Biodiversity is sustainably used and managed and nature’s contributions to people, including ecosystem functions and services, are valued, maintained and enhanced, with those currently in decline being restored, supporting the achievement of sustainable development for the benefit of present and future generations by 2050. |
| Goal C | Share Benefits Fairly - The monetary and non-monetary benefits from the utilization of genetic resources and digital sequence information on genetic resources, and of traditional knowledge associated with genetic resources, as applicable, are shared fairly and equitably, including, as appropriate with indigenous peoples and local communities, and substantially increased by 2050, while ensuring traditional knowledge associated with genetic resources is appropriately protected, thereby contributing to the conservation and sustainable use of biodiversity, in accordance with internationally agreed access and benefit-sharing instruments. |
| Goal D | Invest and Collaborate - Adequate means of implementation, including financial resources, capacity-building, technical and scientific cooperation, and access to and transfer of technology to fully implement the Kunming-Montreal Global Biodiversity Framework are secured and equitably accessible to all Parties, especially developing country Parties, in particular the least developed countries and small island developing States, as well as countries with economies in transition, progressively closing the biodiversity finance gap of \$700 billion per year, and aligning financial flows with the Kunming-Montreal Global Biodiversity Framework and the 2050 Vision for biodiversity. |
| *Reducing threats to biodiversity* |  |
| Target 1 | Plan and Manage all Areas To Reduce Biodiversity Loss |
| Target 2 | Restore 30% of all Degraded Ecosystems |
| Target 3 | Conserve 30% of Land, Waters and Seas |
| Target 4 | Halt Species Extinction, Protect Genetic Diversity, and Manage Human-Wildlife Conflicts |
| Target 5 | Ensure Sustainable, Safe and Legal Harvesting and Trade of Wild Species |
| Target 6 | Reduce the Introduction of Invasive Alien Species by 50% and Minimize Their Impact |
| Target 7 | Reduce Pollution to Levels That Are Not Harmful to Biodiversity |
| Target 8 | Minimize the Impacts of Climate Change on Biodiversity and Build Resilience |
| *Meeting people’s needs through sustainable use and benefit-sharing* |  |
| Target 9 | Manage Wild Species Sustainably To Benefit People |
| Target 10 | Enhance Biodiversity and Sustainability in Agriculture, Aquaculture, Fisheries, and Forestry |
| Target 11 | Restore, Maintain and Enhance Nature’s Contributions to People |
| Target 12 | Enhance Green Spaces and Urban Planning for Human Well-Being and Biodiversity |
| Target 13 | Increase the Sharing of Benefits From Genetic Resources, Digital Sequence Information and Traditional Knowledge |
| *Tools and solutions for implementation and mainstreaming* |  |
| Target 14 | Integrate Biodiversity in Decision-Making at Every Level |
| Target 15 | Businesses Assess, Disclose and Reduce Biodiversity-Related Risks and Negative Impacts |
| Target 16 | Enable Sustainable Consumption Choices To Reduce Waste and Overconsumption |
| Target 17 | Strengthen Biosafety and Distribute the Benefits of Biotechnology |
| Target 18 | Reduce Harmful Incentives by at Least \$500 Billion per Year, and Scale Up Positive Incentives for Biodiversity |
| Target 19 | Mobilize \$200 Billion per Year for Biodiversity From all Sources, Including \$30 Billion Through International Finance |
| Target 20 | Strengthen Capacity-Building, Technology Transfer, and Scientific and Technical Cooperation for Biodiversity |
| Target 21 | Ensure That Knowledge Is Available and Accessible To Guide Biodiversity Action |
| Target 22 | Ensure Participation in Decision-Making and Access to Justice and Information Related to Biodiversity for all |
| Target 23 | Ensure Gender Equality and a Gender-Responsive Approach for Biodiversity Action |

**Table S5.** Models used in the multi-model approach for fitting to low noise data. The list of models is modified from Tittensor et al. (2014), and some have been similarly modified from their traditional form. Reported is the proportion of times a model failed across the indicators (i.e., failure rate).

| # | Model Name | Functional form | Parameters | Failure rate (%) |
|---|---|---|---|---|
| 1 | Linear Model | a + b x | 2 | 0 |
| 2 | Michaelis-Menten | a x / (b + x) | 2 | 26 |
| 3 | Holling Type III | a x^2 / (b + x^2) | 2 | 13 |
| 4 | Holling Type IV | a x^2 / (b + c x + x^2) | 3 | 26 |
| 5 | Rational | (a + b x) / (1 + c x) | 3 | 23 |
| 6 | Exponential | a e^(b x) | 2 | 9 |
| 7 | Monomolecular | a (1 - e^(-b x)) | 2 | 49 |
| 8 | Ricker | a x e^(-b x) | 2 | 6 |
| 9 | Gompertz | a e^(-b e^(-c x)) | 3 | 45 |
| 10 | Weibull | a (1 - e^(-(b x)^c)) | 3 | 55 |
| 11 | Power | a (x + 1)^b | 2 | 6 |
| 12 | Shepherd | a x / (b + x^c) | 3 | 30 |
| 13 | Sigmoid/Logistic | a / (1 + e^(-b (x - c))) | 3 | 43 |
| 14 | Asymptotic | a - b e^(-c x) | 3 | 51 |

**Table S6.** Summary statistics for the Generalized Additive Mixed Models (GAMMs) used to assess temporal trends for each indicator. Reported are the smooth term statistics: Effective Degrees of Freedom (EDF), F-value, and p-value.

| **Indicator** | **EDF** | **F_value** | **p_value** |
|---|---|---|---|
| Annual mean levels of fine particulate matter (e.g. PM2.5 and PM10) in cities | 1.32 | 46.45 | <0.001 |
| Level of water stress: freshwater withdrawal as a proportion of available freshwater resources | 2.81 | 14.85 | <0.001 |
| Proportion of population using safely managed drinking water services | 3.98 | 1301.54 | <0.001 |
| Number of deaths, missing persons and directly affected persons attributed to disasters per 100,000 population | 1 | 7.51 | 0.018 |
| Total number of internationally recognized certificates published on the Access and Benefit-sharing Clearing-House | 4.84 | 1181.53 | <0.001 |
| CO2 emission per unit of value added | 3 | 22.77 | <0.001 |
| Change in water-use efficiency over time | 1 | 44.24 | <0.001 |
| Global imports of information and communications technology goods as presented by bilateral trade flows by information and communications technology goods category | 1.36 | 40.61 | <0.001 |
| Number of island invasive alien species eradications | 4.95 | 5066.18 | <0.001 |
| Cropland nutrient budget | 3.21 | 17.35 | <0.001 |
| Spawning stock biomass (related to commercially exploited species) | 1 | 4.52 | 0.037 |
| Rate of tree cover loss | 1 | 38.92 | <0.001 |
| Mountain Green Cover Index | 1 | 359.23 | <0.001 |
| Forestry production and trade (wood fuel) | 4.84 | 66.63 | <0.001 |
| Number of checkpoint communiquÃ©s published on the Access and Benefit-sharing Clearing-House | 3.34 | 69.49 | <0.001 |
| Red List Index | 3.55 | 17.39 | <0.001 |
| International Public Funding, Including Official Development Assistance (ODA) for Conservation and Sustainable Use of Biodiversity and Ecosystems | 1 | 11.52 | 0.003 |
| Coverage of Protected Areas and Other Effective Area-Based Conservation Measures | 4.91 | 3836.64 | <0.001 |
| Proportion of Fish Stocks within Biologically Sustainable Levels | 1 | 68.47 | <0.001 |
| Rate of Invasive Alien Species Establishment | 4.93 | 24514.86 | <0.001 |
| Progress towards Sustainable Forest Management | 3.35 | 72.27 | <0.001 |
| Positive Incentives in Place to Promote Biodiversity Conservation and Sustainable Use | 4.99 | 23540.17 | <0.001 |
| Value of Subsidies and Other Incentives Harmful to Biodiversity | 1 | 0.27 | 0.612 |
| Protected Area Representativeness & Connectedness Indices | 3.34 | 1119.01 | <0.001 |
| Ecological Footprint | 1 | 208.04 | <0.001 |
| Number of Category 1 Parties to CITES | 4.6 | 555.96 | <0.001 |
| Natural Habitat Extent | 4.24 | 64.26 | <0.001 |
| Pesticide Use | 1 | 97.28 | <0.001 |
| Coverage of Marine Protected Areas and Other Effective Area-Based Conservation Measures | 4.91 | 2169.54 | <0.001 |
| Coverage of Terrestrial Protected Areas and Other Effective Area-Based Conservation Measures | 4.79 | 5460.72 | <0.001 |
| Protected Area Management Assessments | 4.58 | 58.16 | <0.001 |
| Protected Area Coverage of Important Bird and Biodiversity Areas | 4.94 | 3978.96 | <0.001 |
| Protected Area Coverage of Alliance for Zero Extinction Sites | 3.87 | 250.1 | <0.001 |
| Mammal and Bird Extinctions | 3.42 | 120.59 | <0.001 |
| Knowledge Transfer (Number of Biodiversity Papers in Web of Science per Year) | 3.87 | 111.79 | <0.001 |
| Funding Provided by the Global Environment Facility | 1 | 16.08 | <0.001 |
| Number of Countries Implementing Natural Resource Accounts within the System of Environmental-Economic Accounting (SEEA) | 3.24 | 484.94 | 0.002 |
| Fertiliser Use | 3.48 | 32.81 | <0.001 |
| CFC Emissions in ODP Tonnes | 4.46 | 178.11 | <0.001 |
| CITES Trade Database - Number of Trades in Wild Species | 3.01 | 7.71 | <0.001 |
| Number of Countries that Have Ratified the Nagoya Protocol | 4.59 | 1245.77 | <0.001 |
| Number of NBSAPs Submitted Since 2010 | 4.21 | 97.98 | <0.001 |
| Red List Index for Birds, Mammals & Amphibians Showing Trends Driven by Utilisation | 3.55 | 41.12 | <0.001 |
| Red List Index for Birds Showing Trends Driven by Pollution | 3.73 | 181.78 | <0.001 |
| Red List Index for Birds Showing Trends Driven by Invasive Alien Species | 4.1 | 83.91 | <0.001 |
| Number of Countries That Have Ratified the Cartagena Protocol on Biosafety to the Convention on Biological Diversity | 4.53 | 55.84 | <0.001 |
| Proportion of seats held by women in (a) national parliaments and (b) local governments | 2.93 | 1389.04 | <0.001 |

