## Supplementary Factsheets for "Global failure to track international biodiversity targets on islands"

### Indicator Factsheets

October 25, 2026

### Table of Contents

**Target 1. Plan and Manage all Areas To Reduce Biodiversity Loss**

- Natural Habitat Extent
- Rate of tree cover loss

**Target 2. Restore 30% of all Degraded Ecosystems**

**Target 3. Conserve 30% of Land, Waters and Seas**

- Coverage of Protected Areas and Other Effective Area-Based Conservation Measures
- Coverage of Marine Protected Areas and Other Effective Area-Based Conservation Measures
- Coverage of Terrestrial Protected Areas and Other Effective Area-Based Conservation Measures
- Protected Area Representativeness & Connectedness Indices
- Protected Area Management Assessments
- Protected Area Coverage of Important Bird and Biodiversity Areas
- Protected Area Coverage of Alliance for Zero Extinction Sites
- Mountain Green Cover Index

**Target 4. Halt Species Extinction, Protect Genetic Diversity, and Manage Human-Wildlife Conflicts**

- Red List Index
- Mammal and Bird Extinctions
- Number of island invasive alien species eradications

**Target 5. Ensure Sustainable, Safe and Legal Harvesting and Trade of Wild Species**

- Proportion of Fish Stocks within Biologically Sustainable Levels
- Red List Index for Birds, Mammals & Amphibians Showing Trends Driven by Utilisation
- Number of Category 1 Parties to CITES
- CITES Trade Database - Number of Trades in Wild Species

**Target 6. Reduce the Introduction of Invasive Alien Species by 50% and Minimize Their Impact**

- Rate of Invasive Alien Species Establishment
- Red List Index for Birds Showing Trends Driven by Invasive Alien Species

**Target 7. Reduce Pollution to Levels That Are Not Harmful to Biodiversity**

- Pesticide Use
- Fertiliser Use
- CFC Emissions in ODP Tonnes
- Red List Index for Birds Showing Trends Driven by Pollution
- Cropland nutrient budget

**Target 8. Minimize the Impacts of Climate Change on Biodiversity and Build Resilience**

**Target 9. Manage Wild Species Sustainably To Benefit People**

- Spawning stock biomass (related to commercially exploited species)

**Target 10. Enhance Biodiversity and Sustainability in Agriculture, Aquaculture, Fisheries, and Forestry**

- Progress towards Sustainable Forest Management
- Forestry production and trade (wood fuel)

**Target 11. Restore, Maintain and Enhance Nature's Contributions to People**

- Annual mean levels of fine particulate matter (e.g. PM2.5 and PM10) in cities
- Level of water stress: freshwater withdrawal as a proportion of available freshwater resources
- Proportion of population using safely managed drinking water services
- Number of deaths, missing persons and directly affected persons attributed to disasters per 100,000 population

**Target 12. Enhance Green Spaces and Urban Planning for Human Well-Being and Biodiversity**

**Target 13. Increase the Sharing of Benefits From Genetic Resources, Digital Sequence Information and Traditional Knowledge**

- Total number of internationally recognized certificates published on the Access and Benefit-sharing Clearing-House
- Number of Countries that Have Ratified the Nagoya Protocol
- Number of checkpoint communiqués published on the Access and Benefit-sharing Clearing-House

**Target 14. Integrate Biodiversity in Decision-Making at Every Level**

- Number of Countries Implementing Natural Resource Accounts within the System of Environmental-Economic Accounting (SEEA)
- Number of NBSAPs Submitted Since 2010

**Target 15. Businesses Assess, Disclose and Reduce Biodiversity-Related Risks and Negative Impacts**

**Target 16. Enable Sustainable Consumption Choices To Reduce Waste and Overconsumption**

- Ecological Footprint
- CO2 emission per unit of value added
- Change in water-use efficiency over time

**Target 17. Strengthen Biosafety and Distribute the Benefits of Biotechnology**

- Number of Countries That Have Ratified the Cartagena Protocol on Biosafety to the Convention on Biological Diversity

**Target 18. Reduce Harmful Incentives by at Least $500 Billion per Year, and Scale Up Positive Incentives for Biodiversity**

- Positive Incentives in Place to Promote Biodiversity Conservation and Sustainable Use
- Value of Subsidies and Other Incentives Harmful to Biodiversity

**Target 19. Mobilize $200 Billion per Year for Biodiversity From all Sources, Including $30 Billion Through International Finance**

- International Public Funding, Including Official Development Assistance (ODA) for Conservation and Sustainable Use of Biodiversity and Ecosystems
- Funding Provided by the Global Environment Facility

**Target 20. Strengthen Capacity-Building, Technology Transfer, and Scientific and Technical Cooperation for Biodiversity**

- Global imports of information and communications technology goods as presented by bilateral trade flows by information and communications technology goods category

**Target 21. Ensure That Knowledge Is Available and Accessible To Guide Biodiversity Action**

- Knowledge Transfer (Number of Biodiversity Papers in Web of Science per Year)

**Target 22. Ensure Participation in Decision-Making and Access to Justice and Information Related to Biodiversity for all**

**Target 23. Ensure Gender Equality and a Gender-Responsive Approach for Biodiversity Action**

- Proportion of seats held by women in (a) national parliaments and (b) local governments

#### Target 1

##### Plan and Manage all Areas To Reduce Biodiversity Loss

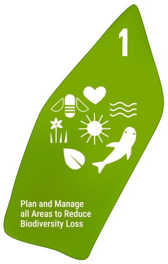


Ensure that all areas are under participatory, integrated, and biodiversity inclusive spatial planning and/or effective management processes addressing land and sea use change, to bring the loss of areas of high biodiversity importance, including ecosystems of high ecological integrity, close to zero by 2030, while respecting the rights of indigenous peoples and local communities.

#### Headline indicator: Natural Habitat Extent

##### Description

Proportion of land which has not been converted to agricultural use by a country.

##### Temporal Trends

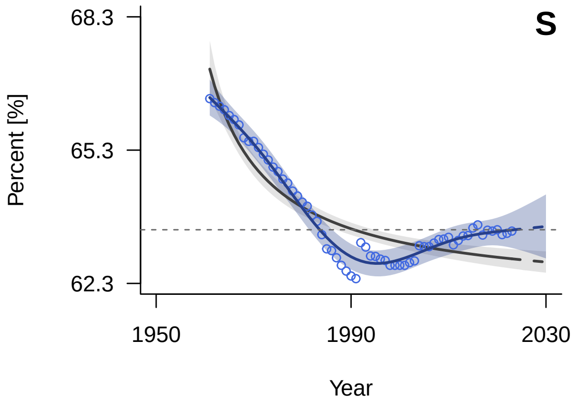


*General Additive Mixed Model (GAMM) fitted trend (solid line) with 95% confidence interval (shaded area). The dashed continuation of the trend line represents the extrapolated future trajectory until 2030. Blue indicates positive tracking towards the target; orange indicates negative tracking. The horizontal grey dashed line marks the 2020 benchmark value. The grey line (solid and dashed) and shading show the model averaging trend and confidence interval.*

##### Spatial Coverage

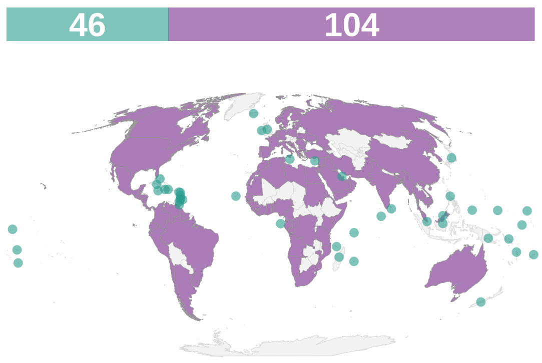


*Spatial coverage of data availability across 150 coastal and island nations. Purple: coastal countries; teal: island countries; grey: countries without data or landlocked. The bars on top of the map indicate the count of countries in each category (purple: coastal countries; teal: island countries; grey: no data).*

##### Key Information

| | |
|---|---|
| **Units** | Percent [%] |
| **Time Period** | 1961-2022 |
| **Disaggregation** | Country |
| **Source** | FAO, https://www.fao.org/faostat/en/#data/RL/metadata |
| **Accession Date** | Apr-26 |
| **DPSIR Framework** | State |
| **Theory of Change (ToC)** | N/A |

##### Methodology

Data were extracted from the FAOSTAT Land Use Indicators (Inputs_LandUse_E) dataset. We used the `NOFLAG` version, keeping all records regardless of data quality flags (which included `A` - Official figure, `E` - Estimated value, `I` - Value imputed by a receiving agency, and `X` - Figure from external organization). We filtered the raw data to retrieve the area in hectares (Element 5110) for two specific categories: total land area (Item 6601) and agricultural land (Item 6610). Data were aggregated by year, and the non-agricultural proportion was calculated.

#### Notes & Limitations

- GBF official headline indicator (Extent of natural ecosystems) is computed differently, but data are not available. Thus, we used the FAOSTAT Land Use dataset as in Tittensor et al. (2014).
- The Natural habitat extent includes also urban areas, because they cannot be separated from other categories in the FAOSTAT database. Additionally, this indicator cannot account for land that has been abandoned after agricultural use.
- There is a sudden artificial increase in the indicator value around 1992. Because the USSR dissolved and our methodology excludes landlocked countries, we drop highly agricultural countries (e.g., Kazakhstan, Uzbekistan) while retaining nations with expansive forests (e.g., Russia's Siberian forest). Nonetheless, the overall observed trend remains similar to the findings in Tittensor et al. (2014).

#### Complementary indicator: Rate of tree cover loss

##### Description

This indicator tracks the annual spatial extent and speed of tree canopy canopy removal due to human activities or natural disturbances.

##### Temporal Trends

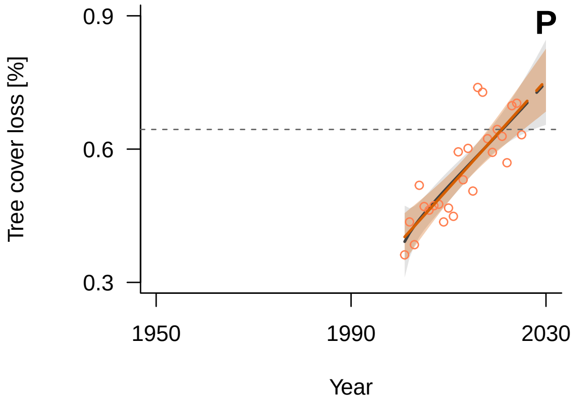


*General Additive Mixed Model (GAMM) fitted trend (solid line) with 95% confidence interval (shaded area). The dashed continuation of the trend line represents the extrapolated future trajectory until 2030. Blue indicates positive tracking towards the target; orange indicates negative tracking. The horizontal grey dashed line marks the 2020 benchmark value. The grey line (solid and dashed) and shading show the model averaging trend and confidence interval.*

##### Spatial Coverage

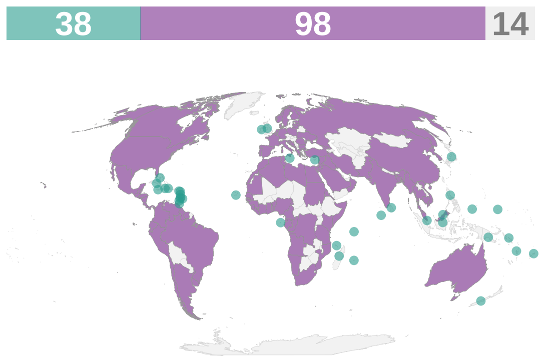


*Spatial coverage of data availability across 150 coastal and island nations. Purple: coastal countries; teal: island countries; grey: countries without data or landlocked. The bars on top of the map indicate the count of countries in each category (purple: coastal countries; teal: island countries; grey: no data).*

##### Key Information

| | |
|---|---|
| **Units** | Tree cover loss [%] |
| **Time Period** | 2001-2025 |
| **Disaggregation** | Country |
| **Source** | Global Forest Watch, https://www.globalforestwatch.org/ |
| **Accession Date** | Jul-26 |
| **DPSIR Framework** | Pressure |
| **Theory of Change (ToC)** | N/A |

##### Methodology

The dataset relies on subnational estimates of yearly tree cover extent. These estimates were aggregated to the country level, and the annual percentage rate of tree cover loss was derived by comparing each year's total extent against the baseline extent from the year 2000.

##### Notes & Limitations

N/A

### Target 2

#### Restore 30% of all Degraded Ecosystems

Ensure that by 2030 at least 30 per cent of areas of degraded terrestrial, inland water, and coastal and marine ecosystems are under effective restoration, in order to enhance biodiversity and ecosystem functions and services, ecological integrity and connectivity.

*No indicators available for this target.*

### Target 3

#### Conserve 30% of Land, Waters and Seas

Ensure and enable that by 2030 at least 30 per cent of terrestrial, inland water, and of coastal and marine areas, especially areas of particular importance for biodiversity and ecosystem functions and services, are effectively conserved and managed through ecologically representative, well-connected and equitably governed systems of protected areas and other effective area-based conservation measures, recognizing indigenous and traditional territories where applicable, and integrated into wider landscapes, seascapes and the ocean, while ensuring that any sustainable use, where appropriate in such areas, is fully consistent with conservation outcomes, recognizing and respecting the rights of indigenous peoples and local communities, including over their traditional territories.

### Headline indicator: Coverage of Protected Areas and Other Effective Area-Based Conservation Measures

#### Description

The total area of terrestrial, inland water, coastal, and marine ecosystems combined that are formally recognized, dedicated, and managed for long-term biodiversity conservation.

#### Temporal Trends

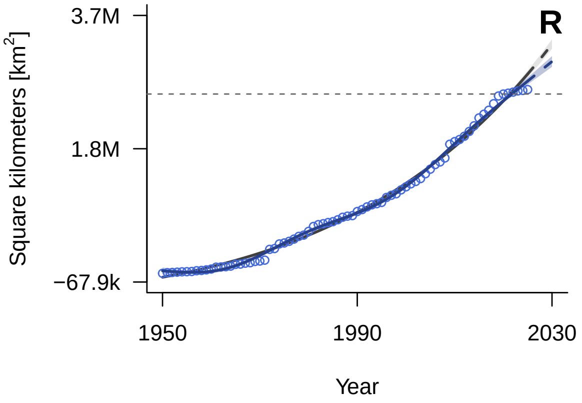


*General Additive Mixed Model (GAMM) fitted trend (solid line) with 95% confidence interval (shaded area). The dashed continuation of the trend line represents the extrapolated future trajectory until 2030. Blue indicates positive tracking towards the target; orange indicates negative tracking. The horizontal grey dashed line marks the 2020 benchmark value. The grey line (solid and dashed) and shading show the model averaging trend and confidence interval.*

#### Spatial Coverage

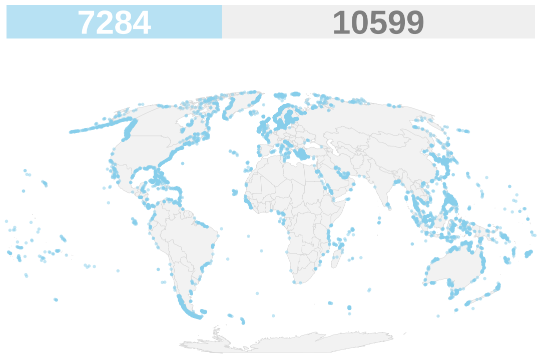


*Spatial coverage of data availability across islands globally. Islands with available data are represented in light blue. The bars on top of the map indicate how many of all islands larger than 1 km² globally have data (light blue) or not (grey).*

#### Key Information

| | |
|---|---|
| **Units** | Square kilometers [km²] |
| **Time Period** | 1950-2024 |
| **Disaggregation** | Spatially explicit |
| **Source** | Protected Planet, https://www.protectedplanet.net/en |
| **Accession Date** | Mar-26 |
| **DPSIR Framework** | Response |
| **Theory of Change (ToC)** | Output |

#### Methodology

We extracted global protected area data from the World Database on Protected Areas (WDPA) and the World Database on Other Effective Area-based Conservation Measures (WD-OECM) and processed data using QGIS 3.34.15. Both polygon and point records were reprojected to the Eckert IV equal-area projection (EPSG: 54012). For polygons, we removed areas with a reported size of zero. For points, we estimated their physical area by generating circular buffers, defining the radius such that the resulting circular area equals the site's reported area. The polygon and buffer layers were then merged to create a global base layer of protected areas and OECMs.

To calculate the total coverage of PAs and OECMs on islands, we first merged the terrestrial island polygons with their surrounding marine shelf (defined as the continental shelf within a 12-nautical-mile buffer and a bathymetry of 0 to 200 meters below sea level, GEBCO Bathymetric Compilation Group 2023; following Mouillot et al. 2020). We then intersected this island-and-shelf layer with the global base layer, using the `vectra` R package to streamline the spatial union process. We removed all overlapping PAs and retained the overlapping fraction that had the earliest known designation year. Finally, we calculated the area in square kilometers for each polygon and computed the cumulative sum of PAs and OECMs across all islands and the respective shelf over time.

##### Notes & Limitations

- The 12-nautical-mile buffer used to delineate the surrounding marine shelf represents a political/economic boundary rather than a biological one.
- National reporting delays or omissions may result in incomplete spatial coverage.
- The indicator measures formal legal designation, which does not guarantee active enforcement or conservation success ("paper parks").
- Utilizing establishment years introduces temporal error for sites with missing dates or those experiencing subsequent boundary and status revisions.

### Headline indicator: Coverage of Marine Protected Areas and Other Effective Area-Based Conservation Measures

#### Description

The total area of marine and coastal ecosystems formally recognized, dedicated, and managed for long-term biodiversity conservation.

##### Temporal Trends

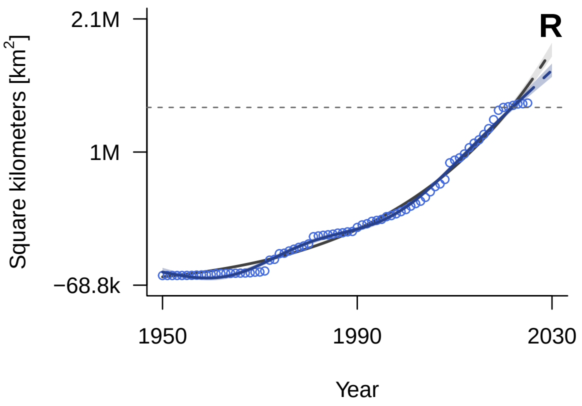


*General Additive Mixed Model (GAMM) fitted trend (solid line) with 95% confidence interval (shaded area). The dashed continuation of the trend line represents the extrapolated future trajectory until 2030. Blue indicates positive tracking towards the target; orange indicates negative tracking. The horizontal grey dashed line marks the 2020 benchmark value. The grey line (solid and dashed) and shading show the model averaging trend and confidence interval.*

##### Spatial Coverage

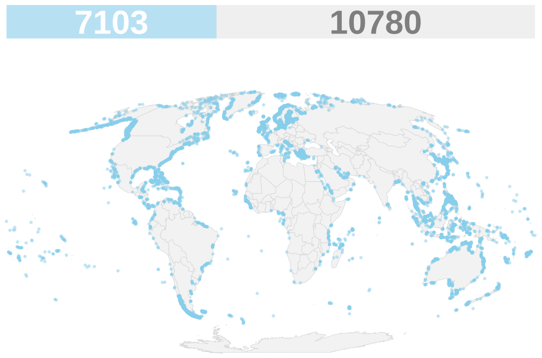


*Spatial coverage of data availability across islands globally. Islands with available data are represented in light blue. The bars on top of the map indicate how many of all islands larger than 1 km$^2$ globally have data (light blue) or not (grey).*

#### Key Information

| | |
|---|---|
| **Units** | Square kilometers [km$^2$] |
| **Time Period** | 1950-2023 |
| **Disaggregation** | Spatially explicit |
| **Source** | Protected Planet, https://www.protectedplanet.net/en |
| **Accession Date** | Mar-26 |
| **DPSIR Framework** | Response |
| **Theory of Change (ToC)** | Output |

#### Methodology

We extracted global protected area data from the World Database on Protected Areas (WDPA) and the World Database on Other Effective Area-based Conservation Measures (WD-OECM) and processed data using QGIS 3.34.15. Both polygon and point records were reprojected to the Eckert IV equal-area projection (EPSG: 54012). For polygons, we removed areas with a reported size of zero. For points, we estimated their physical area by generating circular buffers, defining the radius such that the resulting circular area equals the site's reported area. The polygon and buffer layers were then merged to create a global base layer of protected areas and OECMs.

To calculate the coverage of marine PAs and OECMs for each island, we first delineated the relevant marine shelf territory (defined as the continental shelf within a 12-nautical-mile buffer and a bathymetry of 0 to 200 meters below sea level, GEBCO Bathymetric Compilation Group 2023; following Weigelt et al. 2013, Mouillot et al. 2020). We then intersected this island marine shelf layer with the global base layer, using the `vectra` R package to streamline the spatial union process. We removed all overlapping PAs and retained the overlapping fraction that had the earliest known designation year. Finally, we calculated the area in square kilometers for each polygon and computed the cumulative sum of PAs and OECMs across all island shelves over time.

##### Notes & Limitations

- The 12-nautical-mile buffer used to delineate the surrounding marine shelf represents a political/economic boundary rather than a biological one.
- National reporting delays or omissions may result in incomplete spatial coverage.
- The indicator measures formal legal designation, which does not guarantee active enforcement or conservation success ("paper parks").
- Utilizing establishment years introduces temporal error for sites with missing dates or those experiencing subsequent boundary and status revisions.

#### Headline indicator: Coverage of Terrestrial Protected Areas and Other Effective Area-Based Conservation Measures

##### Description

The total area of land and inland water ecosystems formally recognized, dedicated, and managed for long-term biodiversity conservation.

##### Temporal Trends

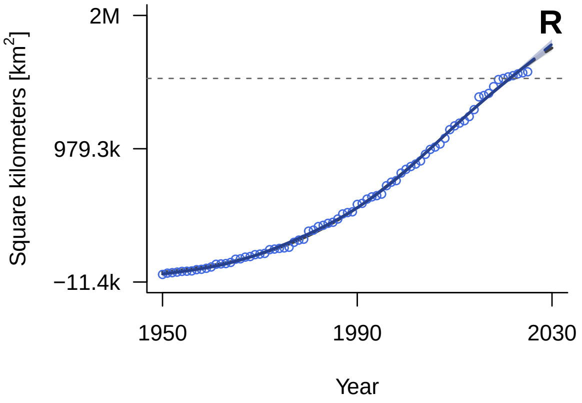


*General Additive Mixed Model (GAMM) fitted trend (solid line) with 95% confidence interval (shaded area). The dashed continuation of the trend line represents the extrapolated future trajectory until 2030. Blue indicates positive tracking towards the target; orange indicates negative tracking. The horizontal grey dashed line marks the 2020 benchmark value. The grey line (solid and dashed) and shading show the model averaging trend and confidence interval.*

##### Spatial Coverage

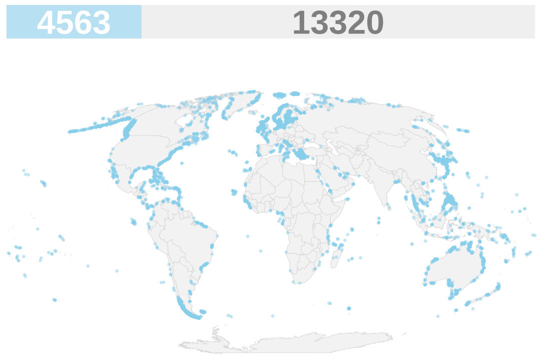


*Spatial coverage of data availability across islands globally. Islands with available data are represented in light blue. The bars on top of the map indicate how many of all islands larger than 1 km$^2$ globally have data (light blue) or not (grey).*

##### Key Information

| | |
|---|---|
| **Units** | Square kilometers [km$^2$] |
| **Time Period** | 1950-2023 |
| **Disaggregation** | Spatially explicit |
| **Source** | Protected Planet, https://www.protectedplanet.net/en |
| **Accession Date** | Mar-26 |
| **DPSIR Framework** | Response |
| **Theory of Change (ToC)** | Output |

##### Methodology

We extracted global protected area data from the World Database on Protected Areas (WDPA) and the World Database on Other Effective Area-based Conservation Measures (WD-OECM) and processed data using QGIS 3.34.15. Both polygon and point records were reprojected to the Eckert IV equal-area projection (EPSG: 54012). For polygons, we removed areas with a reported size of zero. For points, we estimated their physical area by generating circular buffers, defining the radius such that the resulting circular area equals the site's reported area. The polygon and buffer layers were then merged to create a global base layer of protected areas and OECMs.

To calculate the coverage of terrestrial PAs and OECMs on islands, we first intersected the terrestrial island polygons (Weigelt et al. 2013) with the global base layer. We removed all overlapping PAs and retained the overlapping fraction that had the earliest known designation year. We subsequently processed the exported data in R 4.4.2, where we systematically excluded records classified strictly as “Marine” and retained only “Terrestrial” and “Coastal” PAs. Finally, we calculated the cumulative sum of terrestrial PAs and OECMs across all islands over time.

**Notes & Limitations**

- National reporting delays or omissions may result in incomplete spatial coverage.
- The indicator measures formal legal designation, which does not guarantee active enforcement or conservation success (“paper parks”).
- Utilizing establishment years introduces temporal error for sites with missing dates or those experiencing subsequent boundary and status revisions.

#### Component indicator: Protected Area Representativeness & Connectedness Indices

##### Description

The Protected Area Representativeness and Connectedness (PARC) indices measure the extent to which terrestrial protected areas, and other effective area-based conservation measures (OECMs), are ecologically representative, and well-connected (both to one another, and to other areas of intact natural ecosystems in the surrounding landscape).

##### Temporal Trends

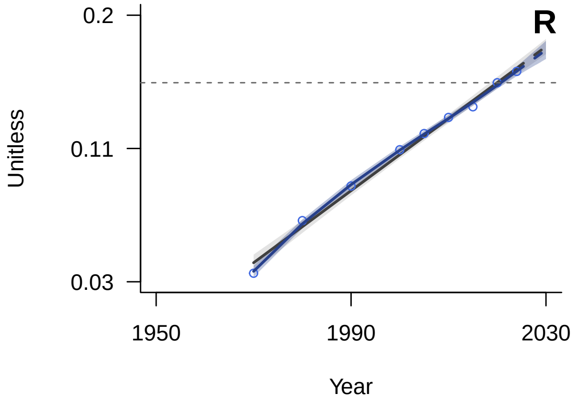


*General Additive Mixed Model (GAMM) fitted trend (solid line) with 95% confidence interval (shaded area). The dashed continuation of the trend line represents the extrapolated future trajectory until 2030. Blue indicates positive tracking towards the target; orange indicates negative tracking. The horizontal grey dashed line marks the 2020 benchmark value. The grey line (solid and dashed) and shading show the model averaging trend and confidence interval.*

##### Spatial Coverage

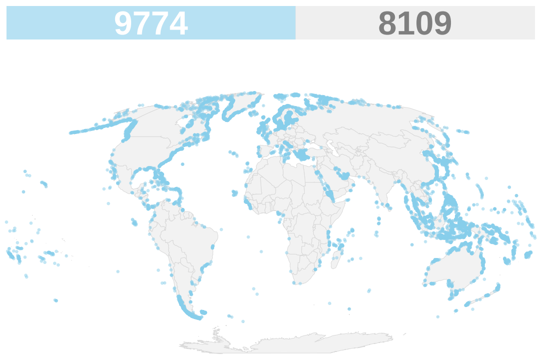


*Spatial coverage of data availability across islands globally. Islands with available data are represented in light blue. The bars on top of the map indicate how many of all islands larger than 1 km$^2$ globally have data (light blue) or not (grey).*

##### Key Information

| | |
|---|---|
| **Units** | Unitless |
| **Time Period** | 1970-2024 |
| **Disaggregation** | Spatially explicit |
| **Source** | CSIRO, https://data.csiro.au/collection/csiro:53973 |
| **Accession Date** | Feb-25 |
| **DPSIR Framework** | Response |
| **Theory of Change (ToC)** | Output |

##### Methodology

We extracted protected area representativeness and connectedness data from the PARC (Protected Area Representativeness Index) 30-arcsecond global time series database. We used the Zonal Statistics tool in QGIS 3.34.15, along with the global island vector layer (Weigelt et al. 2013), to calculate the mean PARC value for each island from the yearly global rasters. These results were then exported as CSV files. Aggregate annual values were calculated by taking the unweighted mean of the PARC values across all retained islands per year.

##### Notes & Limitations

- The global aggregate trend is calculated as the unweighted mean across islands, meaning smaller islands have the same influence on the aggregate trend as large islands regardless of land area.

#### Component indicator: Protected Area Management Assessments

##### Description

The number of documented evaluations of the management effectiveness and conservation outcomes of designated protected and conserved areas.

##### Temporal Trends

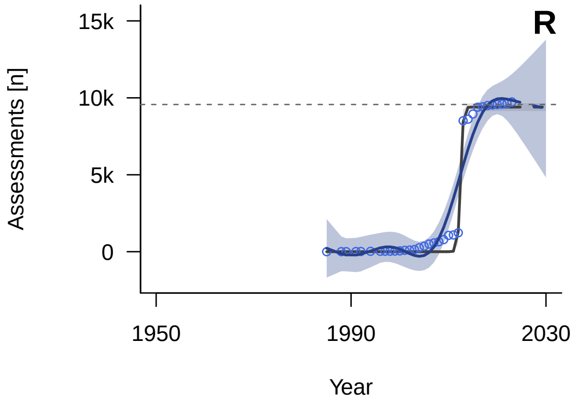


*General Additive Mixed Model (GAMM) fitted trend (solid line) with 95% confidence interval (shaded area). The dashed continuation of the trend line represents the extrapolated future trajectory until 2030. Blue indicates positive tracking towards the target; orange indicates negative tracking. The horizontal grey dashed line marks the 2020 benchmark value. The grey line (solid and dashed) and shading show the model averaging trend and confidence interval.*

##### Spatial Coverage

*No spatial coverage data:*
*Calculated as a global count of assessments.*

##### Key Information

| | |
|---|---|
| **Units** | Assessments [n] |
| **Time Period** | 1985-2023 |
| **Disaggregation** | Assessments |
| **Source** | Protected Planet, https://www.protectedplanet.net/en/thematic-areas/protected-areas-management-effectiveness-pame?tab=Results |
| **Accession Date** | Jan-25 |
| **DPSIR Framework** | Response |
| **Theory of Change (ToC)** | Process |

##### Methodology

We extracted global protected area data from the World Database on Protected Areas (WDPA) and the World Database on Other Effective Area-based Conservation Measures (WD-OECM) and processed data using QGIS 3.34.15. Both polygon and point records were reprojected to the Eckert IV equal-area projection (EPSG: 54012). For polygons, we removed areas with a reported size of zero. For points, we estimated their physical area by generating circular buffers, defining the radius such that the resulting circular area equals the site's reported area. The polygon and buffer layers were then merged to create a global base layer of protected areas and OECMs.

We extracted all areas from the global base layer falling either on islands larger than 1 km2 or within 12 nautical miles from the islands and within 200 m of depth (Weigelt et al., 2013; Mouillot et al. 2020). We then joined the output with the Global Database on Protected Area Management Effectiveness (GD-PAME). The records were then deduplicated to avoid double-counting assessments for protected areas spanning multiple islands. Records missing an assessment date (year = 0) were excluded. Finally, we aggregated the data by counting the yearly assessments across all islands and computing a cumulative sum to produce the running total of management effectiveness assessments over time.

##### Notes & Limitations

- Completing a management effectiveness assessment does not ensure that a protected area is being effectively managed or delivering meaningful conservation outcomes.
- Most assessment frameworks evaluate management processes, inputs, and governance rather than direct biodiversity outcomes, limiting their ability to capture whether protection translates into measurable ecological benefits.

#### Component indicator: Protected Area Coverage of Important Bird and Biodiversity Areas

##### Description

This indicator quantifies the percentage of IBAs (Important Bird and Biodiversity Areas) that is covered by officially designated protected areas.

##### Temporal Trends

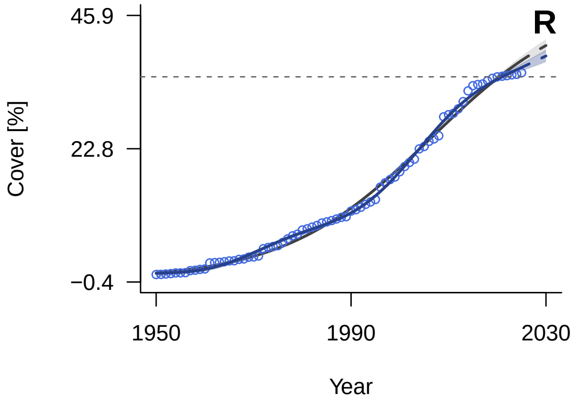


*General Additive Mixed Model (GAMM) fitted trend (solid line) with 95% confidence interval (shaded area). The dashed continuation of the trend line represents the extrapolated future trajectory until 2030. Blue indicates positive tracking towards the target; orange indicates negative tracking. The horizontal grey dashed line marks the 2020 benchmark value. The grey line (solid and dashed) and shading show the model averaging trend and confidence interval.*

##### Spatial Coverage

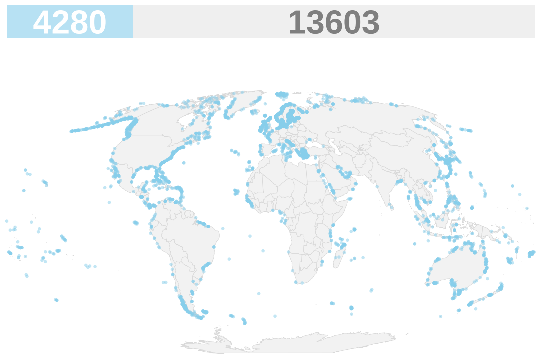


*Spatial coverage of data availability across islands globally. Islands with available data are represented in light blue. The bars on top of the map indicate how many of all islands larger than 1 km$^2$ globally have data (light blue) or not (grey).*

##### Key Information

| | |
|---|---|
| **Units** | Cover [%] |
| **Time Period** | 2005-2024 |
| **Disaggregation** | Spatially explicit |
| **Source** | Bird Life International, https://birdlife.org/ |
| **Accession Date** | Jan-25 |
| **DPSIR Framework** | Response |
| **Theory of Change (ToC)** | Output |

##### Methodology

We extracted global protected area data from the World Database on Protected Areas (WDPA) and the World Database on Other Effective Area-based Conservation Measures (WD-OECM) and processed data using QGIS 3.34.15. Both polygon and point records were reprojected to the Eckert IV equal-area projection (EPSG: 54012). For polygons, we removed areas with a reported size of zero. For points, we estimated their physical area by generating circular buffers, defining the radius such that the resulting circular area equals the site's reported area. The polygon and buffer layers were then merged to create a global base layer of protected areas and OECMs.

To calculate the protected area coverage of Important Bird and Biodiversity Areas (IBAs) on islands, we first merged the terrestrial island polygons from Weigelt et al. (2013) with their surrounding marine shelf (defined as the continental shelf within a 12-nautical-mile buffer and a bathymetry of 0 to 200 meters below sea level, see Mouillot et al. 2020). We then removed all overlapping areas in the global IBA spatial dataset to prevent double-counting and clipped it to the island-and-shelf layer to get the IBA area on islands and their respective shelves. Next, we intersected the island IBAs layer with the global base layer, in which overlapping protected areas had been resolved to the earliest known designation year. Finally, we calculated the cumulative sum of yearly protected IBA area and divided it by the total IBA area on islands to produce the proportion of IBAs covered by protected areas.

##### Notes & Limitations

- The 12-nautical-mile buffer used to delineate the surrounding marine shelf represents a political/economic boundary rather than a biological one.
- National reporting delays or omissions may result in incomplete spatial coverage.
- The physical area of IBAs provided exclusively as point coordinates is estimated via geometric buffering.
- The indicator measures formal legal designation, which does not guarantee active enforcement or conservation success (“paper parks”).
- Utilizing establishment years introduces temporal error for sites with missing dates or those experiencing subsequent boundary and status revisions.

#### Component indicator: Protected Area Coverage of Alliance for Zero Extinction Sites

##### Description

This indicator quantifies the percentage of area of sites holding the last remaining global populations of Endangered or Critically Endangered species that is covered by officially designated protected areas.

##### Temporal Trends

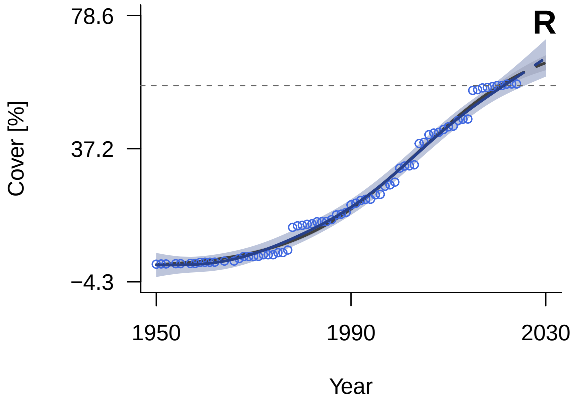


*General Additive Mixed Model (GAMM) fitted trend (solid line) with 95% confidence interval (shaded area). The dashed continuation of the trend line represents the extrapolated future trajectory until 2030. Blue indicates positive tracking towards the target; orange indicates negative tracking. The horizontal grey dashed line marks the 2020 benchmark value. The grey line (solid and dashed) and shading show the model averaging trend and confidence interval.*

##### Spatial Coverage

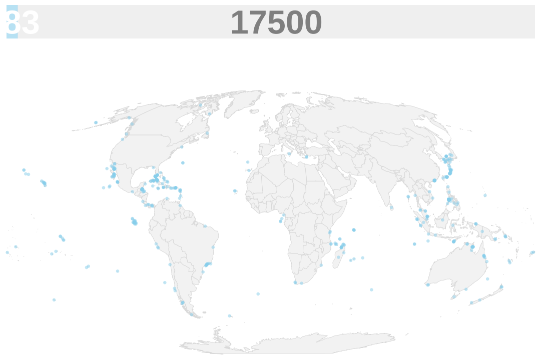


*Spatial coverage of data availability across islands globally. Islands with available data are represented in light blue. The bars on top of the map indicate how many of all islands larger than 1 km$^2$ globally have data (light blue) or not (grey).*

##### Key Information

| | |
|---|---|
| **Units** | Cover [%] |
| **Time Period** | 1994-2024 |
| **Disaggregation** | Spatially explicit |
| **Source** | Bird Life International, https://birdlife.org/ |
| **Accession Date** | Jan-25 |
| **DPSIR Framework** | Response |
| **Theory of Change (ToC)** | Output |

##### Methodology

We extracted global protected area data from the World Database on Protected Areas (WDPA) and the World Database on Other Effective Area-based Conservation Measures (WD-OECM) and processed data using QGIS 3.34.15. Both polygon and point records were reprojected to the Eckert IV equal-area projection (EPSG: 54012). For polygons, we removed areas with a reported size of zero. For points, we estimated their physical area by generating circular buffers, defining the radius such that the resulting circular area equals the site's reported area. The polygon and buffer layers were then merged to create a global base layer of protected areas and OECMs.

To calculate the protected area coverage of Alliance for Zero Extinction (AZE) sites on islands, we first merged the terrestrial island polygons from Weigelt et al. (2013) with their surrounding marine shelf (defined as the continental shelf within a 12-nautical-mile buffer and a bathymetry of 0 to 200 meters below sea level, see Mouillot et al. 2020). We then loaded the global KBA spatial dataset, and filtered out all non-AZE areas. We removed all overlapping areas as well to prevent double-counting and clipped the remaining areas to the island-and-shelf layer to get the AZE area on islands and their respective shelves. Next, we intersected the island AZEs layer with the global base layer, in which overlapping protected areas had been resolved to the earliest known designation year. Finally, we calculated the cumulative sum of yearly protected AZE area and divided it by the total AZE area on islands to produce the proportion of AZEs covered by protected areas.

##### Notes & Limitations

- The 12-nautical-mile buffer used to delineate the surrounding marine shelf represents a political/economic boundary rather than a biological one.
- National reporting delays or omissions may result in incomplete spatial coverage.
- The physical area of AZE sites, as well as protected areas and OECMs, provided exclusively as point coordinates is mathematically estimated via geometric buffering, where the buffer radius is defined such that the resulting circular area equals the site's reported area.
- The indicator measures formal legal designation, which does not guarantee active enforcement or conservation success ("paper parks").
- Utilizing establishment years introduces temporal error for sites with missing dates or those experiencing subsequent boundary and status revisions.
- AZE sites are derived from the KBA dataset, so each site polygon reflects the full KBA management boundary rather than the precise habitat footprint of the trigger species.

#### Complementary indicator: Mountain Green Cover Index

##### Description

This indicator tracks the proportion of mountain land area covered by green vegetation to assess the health and conservation status of mountain ecosystems.

##### Temporal Trends

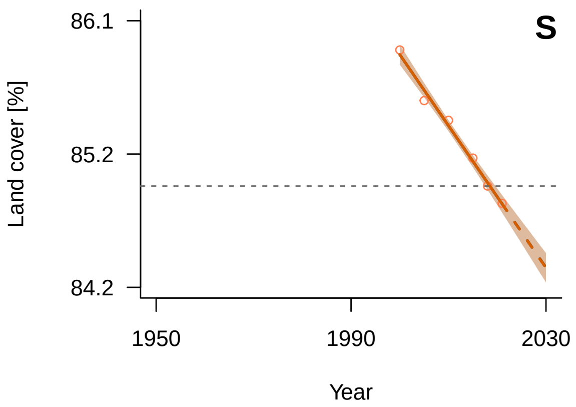


*General Additive Mixed Model (GAMM) fitted trend (solid line) with 95% confidence interval (shaded area). The dashed continuation of the trend line represents the extrapolated future trajectory until 2030. Blue indicates positive tracking towards the target; orange indicates negative tracking. The horizontal grey dashed line marks the 2020 benchmark value. The grey line (solid and dashed) and shading show the model averaging trend and confidence interval.*

##### Spatial Coverage

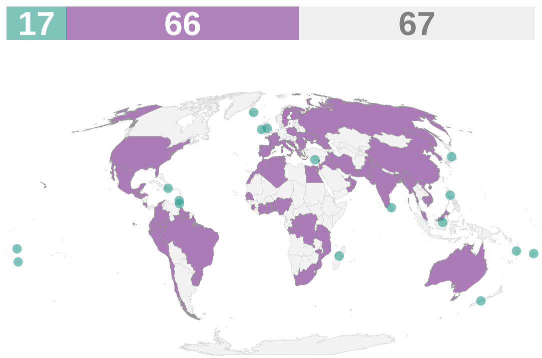


*Spatial coverage of data availability across 150 coastal and island nations. Purple: coastal countries; teal: island countries; grey: countries without data or landlocked. The bars on top of the map indicate the count of countries in each category (purple: coastal countries; teal: island countries; grey: no data).*

##### Key Information

| | |
|---|---|
| **Units** | Land cover [%] |
| **Time Period** | 2000-2021 |
| **Disaggregation** | Country |
| **Source** | UN SDGs, https://unstats.un.org/sdgs/dataportal/database |
| **Accession Date** | Jul-26 |
| **DPSIR Framework** | State |
| **Theory of Change (ToC)** | N/A |

##### Methodology

The dataset was already in a clean, ready-to-use format (country × year).

##### Notes & Limitations

N/A

### Target 4

#### Halt Species Extinction, Protect Genetic Diversity, and Manage Human-Wildlife Conflicts

Ensure urgent management actions to halt human induced extinction of known threatened species and for the recovery and conservation of species, in particular threatened species, to significantly reduce extinction risk, as well as to maintain and restore the genetic diversity within and between populations of native, wild and domesticated species to maintain their adaptive potential, including through in situ and ex situ conservation and sustainable management practices, and effectively manage human-wildlife interactions to minimize human-wildlife conflict for coexistence.

#### Headline indicator: Red List Index

##### Description

The Red List Index (RLI) shows trends in overall extinction risk for species.

##### Temporal Trends

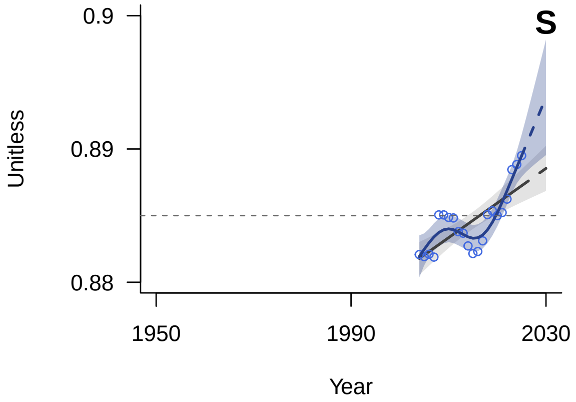


*General Additive Mixed Model (GAMM) fitted trend (solid line) with 95% confidence interval (shaded area). The dashed continuation of the trend line represents the extrapolated future trajectory until 2030. Blue indicates positive tracking towards the target; orange indicates negative tracking. The horizontal grey dashed line marks the 2020 benchmark value. The grey line (solid and dashed) and shading show the model averaging trend and confidence interval.*

##### Spatial Coverage

*No spatial coverage data:*

*The Red List Index is calculated as a global aggregate.*

##### Key Information

| | |
|---|---|
| **Units** | Unitless |
| **Time Period** | 1980-2023 |
| **Disaggregation** | Spatially explicit |
| **Source** | IUCN, https://www.iucnredlist.org |
| **Accession Date** | Feb-26 |
| **DPSIR Framework** | State |
| **Theory of Change (ToC)** | N/A |

##### Methodology

Spatial data were intersected with the global island layer (Weigelt et al., 2013) using QGIS 3.34.15 to generate a unique island-species list. This list served as the basis for querying historical species assessments from the IUCN Red List API via the 'rredlist' package in R 4.4.2. To calculate the Red List Index (RLI) following the methodology of Butchart et al. (2007), we established a baseline of species consistently assessed since 2004. Data Deficient (DD), Not Recognized (NR), and Not Evaluated (NE) species were strictly excluded. We assigned each remaining species a category weight corresponding to its extinction risk (ranging from 0 for Least Concern to 5 for Extinct). We applied a Last-Observation-Carried-Forward (LOCF) approach to fill assessment gaps. We focused on global assessments only.

#### Notes & Limitations

- The official IUCN global RLI includes amphibians, birds, mammals, cycads, and corals. We used only data for birds, mammals, and amphibians. Cycads lack spatial data for islands, and corals are marine and thus not relevant for terrestrial island indicators.
- We could not account for non genuine changes: the information is not exposed in the current API version (v3).
- This indicator may project an overly optimistic view of global extinction risk because the Red List overrepresents well-studied, highly protected taxa like birds. Conversely, severe declines in poorly assessed groups (e.g., invertebrates, plants, and fungi) may go undetected.
- The Red List's baseline effectively excludes pre-Linnaean extinctions, as well as post-Linnaean extinctions that occurred before systematic assessment began.
- Data Deficient (DD) taxa were excluded, yet these poorly documented taxa often face higher extinction risks than fully assessed species.
- The conservative 50-year rule of thumb could delay the recognition of recent extinctions.

#### Headline indicator: Mammal and Bird Extinctions

##### Description

This indicator tracks the cumulative number of documented mammal and bird extinctions occurring on islands.

##### Temporal Trends

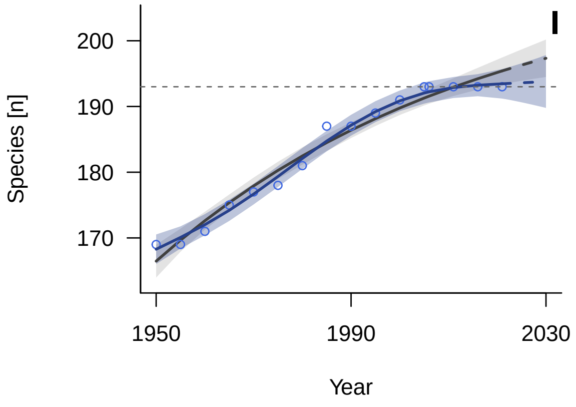


*General Additive Mixed Model (GAMM) fitted trend (solid line) with 95% confidence interval (shaded area). The dashed continuation of the trend line represents the extrapolated future trajectory until 2030. Blue indicates positive tracking towards the target; orange indicates negative tracking. The horizontal grey dashed line marks the 2020 benchmark value. The grey line (solid and dashed) and shading show the model averaging trend and confidence interval.*

##### Spatial Coverage

*No spatial coverage data:*
*Calculated as a global count of extinctions.*

##### Key Information

| | |
|---|---|
| **Units** | Species [n] |
| **Time Period** | 1950-2010 |
| **Disaggregation** | Species |
| **Source** | IUCN Red List, https://www.iucnredlist.org/search |
| **Accession Date** | Jan-25 |
| **DPSIR Framework** | Impact |
| **Theory of Change (ToC)** | N/A |

##### Methodology

We extracted global extinction data for birds and mammals from the IUCN Red List database, filtering specifically for species within the taxonomic classes Aves and Mammalia designated under the category '*EX - Extinct*'. We excluded species lacking a recorded extinction date. For entries containing multiple dates, we retained only the most recent year. We subsequently verified the geographic distribution of the remaining species, retaining exclusively those confirmed to occur on islands. Extinction dates were binned into 5-year intervals, and a cumulative sum was computed to track the running total of island extinctions over time. The Red List is regularly updated and no new island extinctions have been recorded since 2006, the cumulative total remained constant through 2025.

##### Notes & Limitations

- The IUCN Red List has uneven taxonomic coverage; assessments are heavily skewed toward vertebrates while many invertebrate, plant, and fungal groups remain poorly assessed. This may lead to over-optimistic assessments of overall extinction risk, as less-studied taxa could become extinct undetected.
- The Red List's baseline effectively excludes pre-Linnearn extinctions and post-Linnaean extinctions that occurred before systematic assessment began.
- Excluding species that lack a formally recorded extinction date underestimates the true cumulative total of island extinctions.
- The conservative '50-year rule of thumb' could delay the recognition of recent extinctions.

#### Complementary indicator: Number of island invasive alien species eradications

##### Description

This indicator tracks successful management interventions that completely eliminate target invasive alien species populations from island ecosystems.

##### Temporal Trends

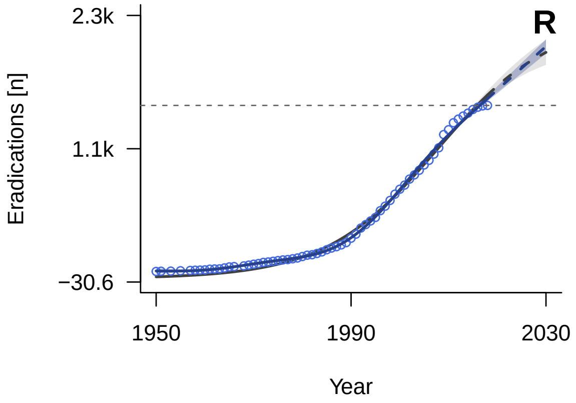


*General Additive Mixed Model (GAMM) fitted trend (solid line) with 95% confidence interval (shaded area). The dashed continuation of the trend line represents the extrapolated future trajectory until 2030. Blue indicates positive tracking towards the target; orange indicates negative tracking. The horizontal grey dashed line marks the 2020 benchmark value. The grey line (solid and dashed) and shading show the model averaging trend and confidence interval.*

##### Spatial Coverage

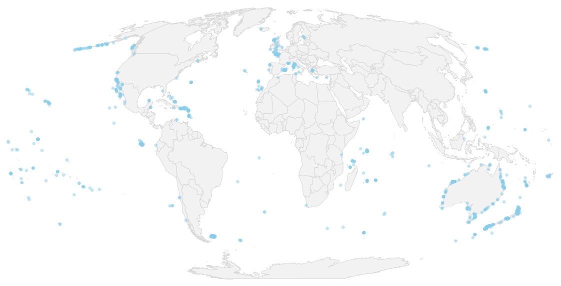

*Spatial coverage of data availability across islands globally, represented in light blue. This dataset includes islands smaller than 1 km$^2$.*

##### Key Information

| | |
|---|---|
| **Units** | Eradications [n] |
| **Time Period** | 1950-2018 |
| **Disaggregation** | Spatially explicit |
| **Source** | DIISE, https://diise.islandconservation.org/ |
| **Accession Date** | Jul-26 |
| **DPSIR Framework** | Response |
| **Theory of Change (ToC)** | Outcome |

##### Methodology

The database was organized by eradication event per island. Single successful eradication events were aggregated by year, excluding the events where the year of eradication was not documented. Then, the cumulative number of successful eradications over time we calculated.

##### Notes & Limitations

- This metric tracks exclusively the eradication of invasive vertebrates. It does not account for the eradication of invasive plants, invertebrates, or other taxa.
- While there may be a recent decline in the sheer number of successful eradications per year, the total treated area continues to increase.
- This indicator was discontinued in 2019 and has no data points after 2020; however, it was retained as the sole exception to this criterion because it is the only island-specific indicator.

### Target 5

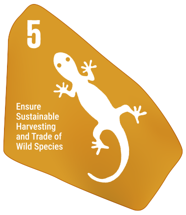


#### Ensure Sustainable, Safe and Legal Harvesting and Trade of Wild Species

Ensure that the use, harvesting and trade of wild species is sustainable, safe and legal, preventing overexploitation, minimizing impacts on non-target species and ecosystems, and reducing the risk of pathogen spill-over, applying the ecosystem approach, while respecting and protecting customary sustainable use by indigenous peoples and local communities.

#### Headline indicator: Proportion of Fish Stocks within Biologically Sustainable Levels

##### Description

Measures the sustainability of the world's marine capture fisheries by the abundance of the exploited fish stocks with respect to maximum sustainable yield levels.

##### Temporal Trends

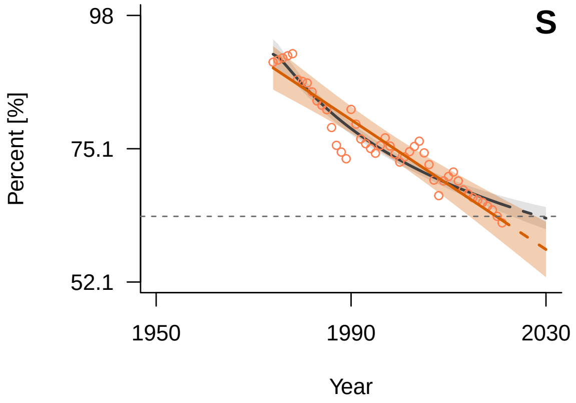


*General Additive Mixed Model (GAMM) fitted trend (solid line) with 95% confidence interval (shaded area). The dashed continuation of the trend line represents the extrapolated future trajectory until 2030. Blue indicates positive tracking towards the target; orange indicates negative tracking. The horizontal grey dashed line marks the 2020 benchmark value. The grey line (solid and dashed) and shading show the model averaging trend and confidence interval.*

##### Spatial Coverage

*No spatial coverage data:*
*Aggregated at the ocean basin level.*

##### Key Information

| | |
|---|---|
| **Units** | Percent [%] |
| **Time Period** | 1974-2021 |
| **Disaggregation** | Regional basins |
| **Source** | UN SDGs, https://unstats.un.org/sdgs/dataportal/database |
| **Accession Date** | Jan-25 |
| **DPSIR Framework** | State |
| **Theory of Change (ToC)** | N/A |

##### Methodology

We extracted data on Sustainable Development Goal (SDG) indicator 14.4.1, which measures the proportion of fish stocks within biologically sustainable levels (Series: Proportion of fish stocks within biologically sustainable levels (not overexploited) (%) ER_H2O_FWTL). Due to significant data gaps at the country level (approximately 92% of national records were missing), we selected the global aggregate series.

#### Notes & Limitations

- Because the data are reported as a single global aggregate, they are not representative of fish stock sustainability for islands.
- This indicator may not be representative of fish stocks that are not actively monitored.

#### Component indicator: Red List Index for Birds, Mammals & Amphibians Showing Trends Driven by Utilisation

##### Description

The Red List Index (RLI) shows trends in overall extinction risk for species. This indicator focuses on trends driven by utilisation for birds, mammals, and amphibians.

##### Temporal Trends

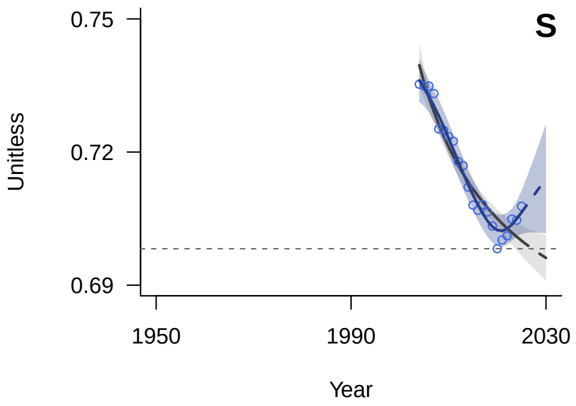


*General Additive Mixed Model (GAMM) fitted trend (solid line) with 95% confidence interval (shaded area). The dashed continuation of the trend line represents the extrapolated future trajectory until 2030. Blue indicates positive tracking towards the target; orange indicates negative tracking. The horizontal grey dashed line marks the 2020 benchmark value. The grey line (solid and dashed) and shading show the model averaging trend and confidence interval.*

##### Spatial Coverage

*No spatial coverage data:*

*The Red List Index is calculated as a global aggregate.*

##### Key Information

| | |
|---|---|
| **Units** | Unitless |
| **Time Period** | 2004-2025 |
| **Disaggregation** | Spatially explicit |
| **Source** | IUCN Red List, https://www.iucnredlist.org |
| **Accession Date** | Feb-26 |
| **DPSIR Framework** | State |
| **Theory of Change (ToC)** | N/A |

##### Methodology

We extracted spatial data from threat category 5.1 - Hunting & trapping terrestrial animals. Spatial data were intersected with the global island layer (Weigelt et al., 2013) using QGIS 3.34.15 to generate a unique island-species list. This list served as the basis for querying historical species assessments from the IUCN Red List API via the 'rredlist' package in R 4.4.2. To calculate the Red List Index (RLI) following the methodology of Butchart et al. (2007), we established a baseline of species consistently assessed since 2004. Data Deficient (DD), Not Recognized (NR), and Not Evaluated (NE) species were strictly excluded. We assigned each remaining species a category weight corresponding to its extinction risk (ranging from 0 for Least Concern to

5 for Extinct). We applied a Last-Observation-Carried-Forward (LOCF) approach to fill assessment gaps. We focused on global assessments only.

**Notes & Limitations**

- We could not account for non genuine changes: the information is not exposed in the current API version (v3).
- This indicator may project an overly optimistic view of global extinction risk because the Red List overrepresents well-studied, highly protected taxa like birds. Conversely, severe declines in poorly assessed groups (e.g., invertebrates, plants, and fungi) may go undetected.
- The Red List's baseline effectively excludes post-Linnaean extinctions that occurred before systematic assessment began.
- Data Deficient (DD) taxa were excluded, yet these poorly documented taxa often face higher extinction risks than fully assessed species.
- The conservative 50-year rule of thumb could delay the recognition of recent extinctions.

#### Complementary indicator: Number of Category 1 Parties to CITES

##### Description

Number of category 1 parties. CITES (the Convention on International Trade in Endangered Species of Wild Fauna and Flora) is an international agreement between governments that aims to ensure that international trade in specimens of wild animals and plants does not threaten the survival of the species.

##### Temporal Trends

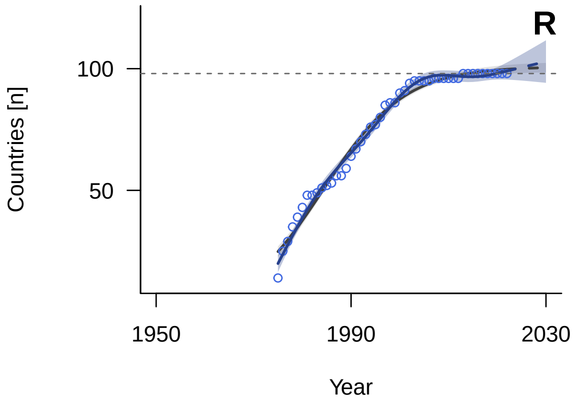


*General Additive Mixed Model (GAMM) fitted trend (solid line) with 95% confidence interval (shaded area). The dashed continuation of the trend line represents the extrapolated future trajectory until 2030. Blue indicates positive tracking towards the target; orange indicates negative tracking. The horizontal grey dashed line marks the 2020 benchmark value. The grey line (solid and dashed) and shading show the model averaging trend and confidence interval.*

##### Spatial Coverage

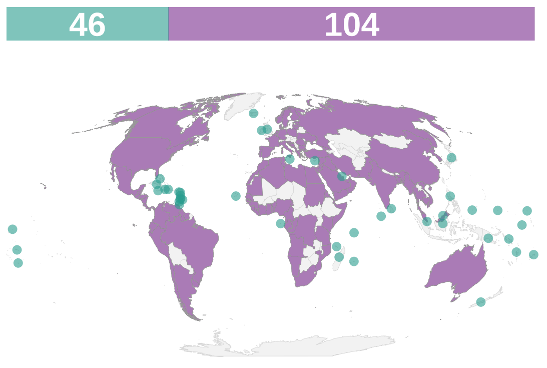


*Spatial coverage of data availability across 150 coastal and island nations. Purple: coastal countries; teal: island countries; grey: countries without data or landlocked. The bars on top of the map indicate the count of countries in each category (purple: coastal countries; teal: island countries; grey: no data).*

##### Key Information

| | |
|---|---|
| **Units** | Countries [n] |
| **Time Period** | 1975-2024 |
| **Disaggregation** | Country |
| **Source** | CITES, https://cites.org/eng/legislation/parties |
| **Accession Date** | Jan-25 |
| **DPSIR Framework** | Response |
| **Theory of Change (ToC)** | Process |

##### Methodology

We extracted data on CITES contracting parties, specifically tracking countries designated as Category 1 (indicating their domestic legislation is believed to generally meet all four requirements for effective CITES implementation). For each country and year, we tracked the entry-into-force date and created a cumulative binary timeline, carrying a value of 1 forward from the year of joining. Annual values were then summed across all retained countries to calculate the total number of Category 1 parties per year.

**Notes & Limitations**

- Measures the adoption of legislation rather than the actual enforcement or sustainability of trade on the ground.
- This indicator is restricted to legal, international trade of CITES-listed species; it does not account for illegal wildlife trade, domestic trade, trade of non-CITES species, or localized consumption.

#### Complementary indicator: CITES Trade Database - Number of Trades in Wild Species

##### Description

The CITES Trade Database is the most comprehensive and authoritative database on international trade in wildlife. CITES (the Convention on International Trade in Endangered Species of Wild Fauna and Flora) is an international agreement between governments. It aims to ensure that international trade in specimens of wild animals and plants does not threaten the survival of the species.

##### Temporal Trends

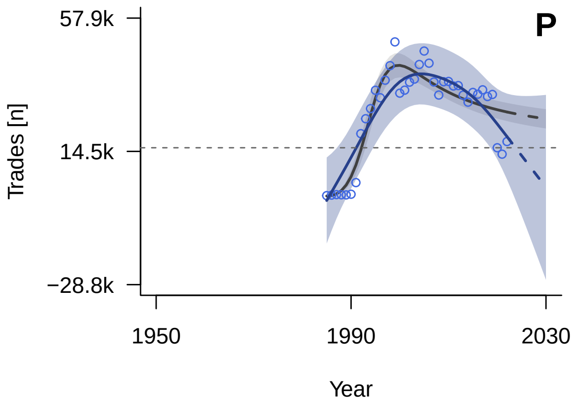


*General Additive Mixed Model (GAMM) fitted trend (solid line) with 95% confidence interval (shaded area). The dashed continuation of the trend line represents the extrapolated future trajectory until 2030. Blue indicates positive tracking towards the target; orange indicates negative tracking. The horizontal grey dashed line marks the 2020 benchmark value. The grey line (solid and dashed) and shading show the model averaging trend and confidence interval.*

##### Spatial Coverage

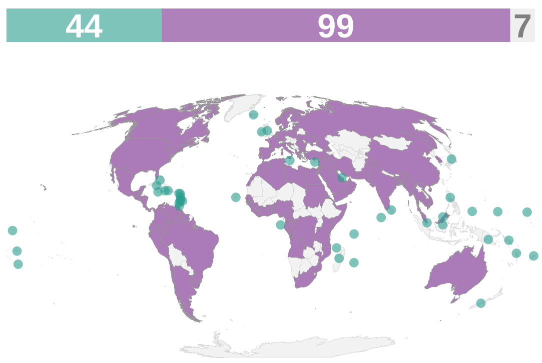


*Spatial coverage of data availability across 150 coastal and island nations. Purple: coastal countries; teal: island countries; grey: countries without data or landlocked. The bars on top of the map indicate the count of countries in each category (purple: coastal countries; teal: island countries; grey: no data).*

##### Key Information

| | |
|---|---|
| **Units** | Trades [n] |
| **Time Period** | 1985-2023 |
| **Disaggregation** | Country |
| **Source** | CITES, https://trade.cites.org/ |
| **Accession Date** | Jan-25 |
| **DPSIR Framework** | Pressure |
| **Theory of Change (ToC)** | N/A |

##### Methodology

We extracted the CITES Trade Database (v2024.1). We included all species across all taxa and filtered for wild-sourced specimens only (Source = "W") traded for hunting (H), personal (P), circus/travelling exhibition (Q), or commercial (C) purposes. When the origin country differed from the exporting country, the origin was used instead. Because quantity units are heterogeneous across records (e.g., number of individuals, kg, $m^3$)

and cannot be meaningfully summed, we used the total number of trade events per exporter country per year instead. The year 2023 was excluded as it returned an anomalously low value (the smallest across the database), indicative of a reporting lag at the time of data extraction.

##### Notes & Limitations

- The indicator counts the number of trade transactions rather than the quantity of specimens or biomass traded.
- CITES trade data are based on permits issued and annual reports submitted by Parties, which may not fully reflect actual trade volumes due to reporting delays, non-compliance, or illegal trade operating outside the permit system.
- The most recent year (2023) was excluded due to a reporting lag; global totals for recent years may therefore be underestimated.

### Target 6

#### Reduce the Introduction of Invasive Alien Species by 50% and Minimize Their Impact

Eliminate, minimize, reduce and or mitigate the impacts of invasive alien species on biodiversity and ecosystem services by identifying and managing pathways of the introduction of alien species, preventing the introduction and establishment of priority invasive alien species, reducing the rates of introduction and establishment of other known or potential invasive alien species by at least 50 per cent, by 2030, eradicating or controlling invasive alien species especially in priority sites, such as islands.

#### Headline indicator: Rate of Invasive Alien Species Establishment

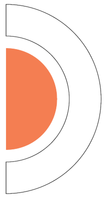

##### Description

The number of new invasive alien species that successfully establish populations within a new country.

##### Temporal Trends

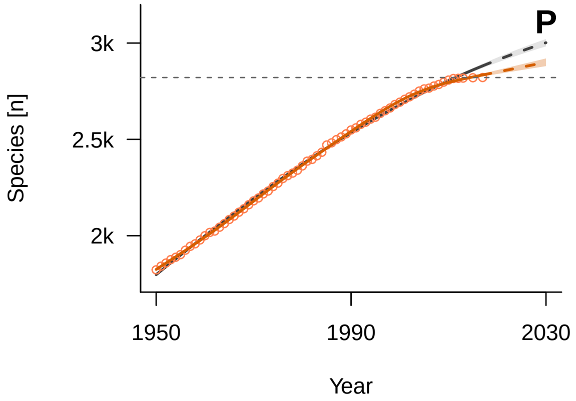


*General Additive Mixed Model (GAMM) fitted trend (solid line) with 95% confidence interval (shaded area). The dashed continuation of the trend line represents the extrapolated future trajectory until 2030. Blue indicates positive tracking towards the target; orange indicates negative tracking. The horizontal grey dashed line marks the 2020 benchmark value. The grey line (solid and dashed) and shading show the model averaging trend and confidence interval.*

##### Spatial Coverage

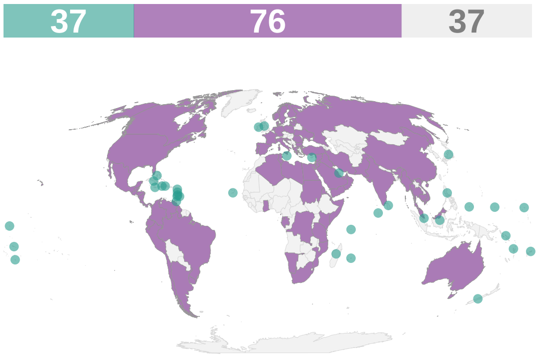


*Spatial coverage of data availability across 150 coastal and island nations. Purple: coastal countries; teal: island countries; grey: countries without data or landlocked. The bars on top of the map indicate the count of countries in each category (purple: coastal countries; teal: island countries; grey: no data).*

##### Key Information

| | |
|---|---|
| **Units** | Species [n] |
| **Time Period** | -142021 |
| **Disaggregation** | Country |
| **Source** | SInAS, https://zenodo.org/records/10038256 |
| **Accession Date** | Jan-25 |
| **DPSIR Framework** | Pressure |
| **Theory of Change (ToC)** | N/A |

##### Methodology

We obtained data from the SInAS database. We excluded records lacking essential data (such as missing geographic locations or introduction dates) and included only UN countries. Territories and jurisdictions (e.g., Hawaiian Islands, Galápagos, French Polynesia) were recoded to their sovereign parent countries. Only the earliest recorded introduction date was retained.

##### Notes & Limitations

- There is a reporting bias towards better-studied regions and taxa.

- The date of first record often lags behind the actual date of introduction or establishment.
- There are temporal reporting biases as delays in the detection, publication, and digitization of species occurrences.
- This indicator likely underestimates the true number of established invasive species due to invasion debt (i.e., the time lags between initial introduction, establishment, and eventual detection).

#### Complementary indicator: Red List Index for Birds Showing Trends Driven by Invasive Alien Species

##### Description

The Red List Index (RLI) shows trends in overall extinction risk for species. This indicator focuses on trends driven by invasive species for birds alone.

##### Temporal Trends

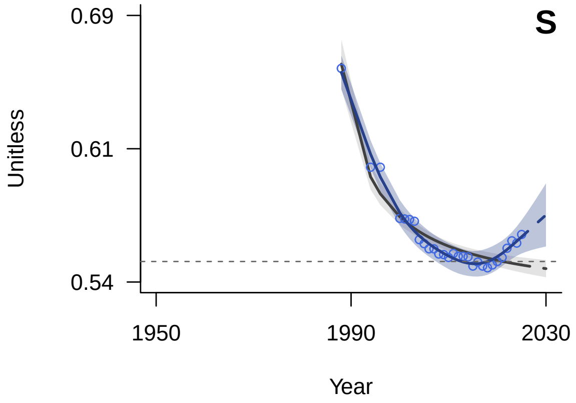


*General Additive Mixed Model (GAMM) fitted trend (solid line) with 95% confidence interval (shaded area). The dashed continuation of the trend line represents the extrapolated future trajectory until 2030. Blue indicates positive tracking towards the target; orange indicates negative tracking. The horizontal grey dashed line marks the 2020 benchmark value. The grey line (solid and dashed) and shading show the model averaging trend and confidence interval.*

##### Spatial Coverage

*No spatial coverage data:*

*The Red List Index is calculated as a global aggregate.*

##### Key Information

| | |
|---|---|
| **Units** | Unitless |
| **Time Period** | 1988-2025 |
| **Disaggregation** | Spatially explicit |
| **Source** | IUCN Red List, https://www.iucnredlist.org |
| **Accession Date** | Feb-26 |
| **DPSIR Framework** | State |
| **Theory of Change (ToC)** | N/A |

##### Methodology

We extracted spatial data for birds from threat category Invasive species. Spatial data were intersected with the global island layer (Weigelt et al., 2013) using QGIS 3.34.15 to generate a unique island-species list. This list served as the basis for querying historical species assessments from the IUCN Red List API via the ‘rredlist’ package in R 4.4.2. To calculate the Red List Index (RLI) following the methodology of Butchart et al. (2007), we established a baseline of species consistently assessed since 1988. Data Deficient (DD), Not Recognized (NR), and Not Evaluated (NE) species were strictly excluded. We assigned each remaining species a category weight corresponding to its extinction risk (ranging from 0 for Least Concern to 5 for

Extinct). We manually excluded one species that feature non-genuine changes, which were primarily driven by taxonomic splits (i.e., *Puffinus bannermani*). We applied a Last-Observation-Carried-Forward (LOCF) approach to fill assessment gaps. We focused on global assessments only.

##### Notes & Limitations

- We could not systematically account for non genuine changes: the information cannot be extracted via API.
- This indicator may project an overly optimistic view of global extinction risk because birds are a well-studied, highly protected taxa. Conversely, severe declines in poorly assessed groups (e.g., invertebrates, plants, and fungi) may go undetected.
- The Red List's baseline effectively excludes post-Linnaean extinctions that occurred before systematic assessment began.
- Data Deficient (DD) taxa were excluded, yet these poorly documented taxa often face higher extinction risks than fully assessed species.
- The conservative 50-year rule of thumb could delay the recognition of recent extinctions.

### Target 7

#### Reduce Pollution to Levels That Are Not Harmful to Biodiversity

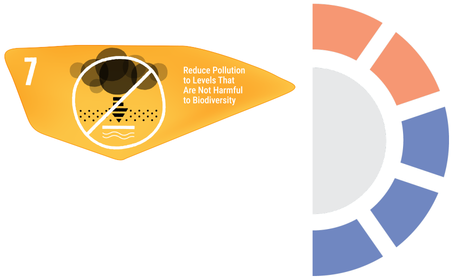


Reduce pollution risks and the negative impact of pollution from all sources, by 2030, to levels that are not harmful to biodiversity and ecosystem functions and services, considering cumulative effects, including: reducing excess nutrients lost to the environment by at least half including through more efficient nutrient cycling and use; reducing the overall risk from pesticides and highly hazardous chemicals by at least half including through integrated pest management, based on science, taking into account food security and livelihoods; and also preventing, reducing, and working towards eliminating plastic pollution.

#### Component indicator: Pesticide Use

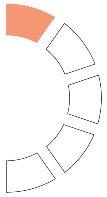

##### Description

The total amount of agricultural pesticides, measured in tonnes of active ingredients, distributed or sold for use.

##### Temporal Trends

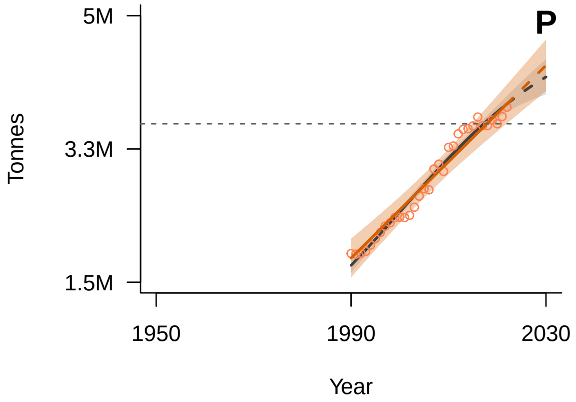


*General Additive Mixed Model (GAMM) fitted trend (solid line) with 95% confidence interval (shaded area). The dashed continuation of the trend line represents the extrapolated future trajectory until 2030. Blue indicates positive tracking towards the target; orange indicates negative tracking. The horizontal grey dashed line marks the 2020 benchmark value. The grey line (solid and dashed) and shading show the model averaging trend and confidence interval.*

##### Spatial Coverage

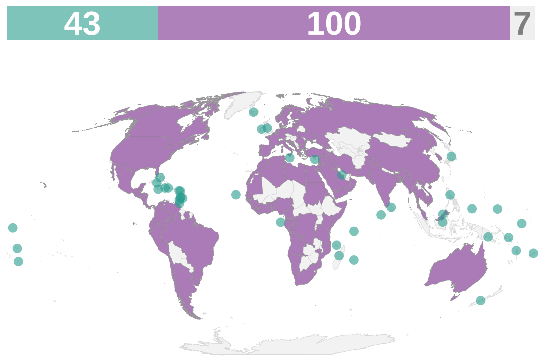


*Spatial coverage of data availability across 150 coastal and island nations. Purple: coastal countries; teal: island countries; grey: countries without data or landlocked. The bars on top of the map indicate the count of countries in each category (purple: coastal countries; teal: island countries; grey: no data).*

##### Key Information

| | |
|---|---|
| **Units** | Tonnes |
| **Time Period** | 1990-2022 |
| **Disaggregation** | Country |
| **Source** | FAO, https://www.fao.org/faostat/en/#data/RP |
| **Accession Date** | Jan-25 |
| **DPSIR Framework** | Pressure |
| **Theory of Change (ToC)** | N/A |

##### Methodology

We extracted agricultural pesticide use data from the Food and Agriculture Organization (FAOSTAT) database. The dataset includes estimated, imputed, and official values. For each country and year, we used the total pesticide use in tonnes (t) of active ingredients. Annual values were then summed across all retained countries to calculate the global annual totals.

#### Notes & Limitations

- Because the data are reported at the national scale, they are not representative of pesticide use on islands.
- Limited to agricultural use, excluding industrial, technical, or public health applications.
- Reporting by countries is sporadic, with a global rate of 32% and a low of 10% for Oceania. Consequently, a large fraction of the data is imputed, and the FAO relies heavily on proxies such as pesticide sales.

#### Component indicator: Fertiliser Use

##### Description

This indicator tracks the quantity of mineral and chemical fertilizers used in agriculture.

##### Temporal Trends

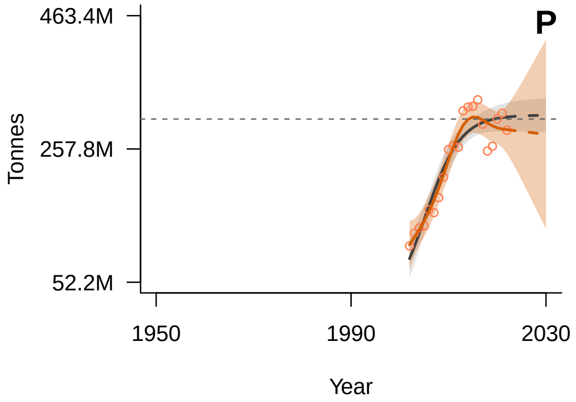


*General Additive Mixed Model (GAMM) fitted trend (solid line) with 95% confidence interval (shaded area). The dashed continuation of the trend line represents the extrapolated future trajectory until 2030. Blue indicates positive tracking towards the target; orange indicates negative tracking. The horizontal grey dashed line marks the 2020 benchmark value. The grey line (solid and dashed) and shading show the model averaging trend and confidence interval.*

##### Spatial Coverage

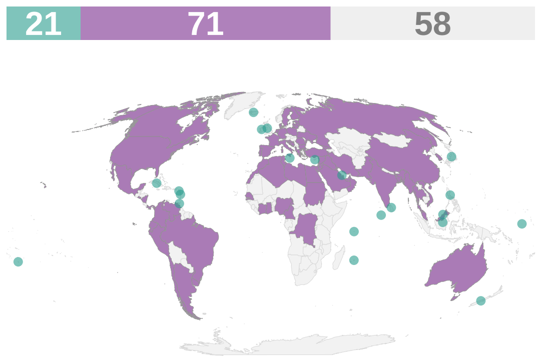


*Spatial coverage of data availability across 150 coastal and island nations. Purple: coastal countries; teal: island countries; grey: countries without data or landlocked. The bars on top of the map indicate the count of countries in each category (purple: coastal countries; teal: island countries; grey: no data).*

##### Key Information

| | |
|---|---|
| **Units** | Tonnes |
| **Time Period** | 2002-2022 |
| **Disaggregation** | Country |
| **Source** | FAO, https://www.fao.org/faostat/en/#data/RFB/visualize |
| **Accession Date** | Jan-25 |
| **DPSIR Framework** | Pressure |
| **Theory of Change (ToC)** | N/A |

##### Methodology

We extracted agricultural fertilizer use data from the Food and Agriculture Organization (FAOSTAT) database, specifically focusing on fertilizers by nutrient. The dataset includes estimated, imputed, and official values for nitrogen (N), phosphate ($P_2O_5$), and potash ($K_2O$). For each country and year, we summed values across all fertilizer nutrient types to produce total fertilizer use in tonnes (t) per country per year. Annual values were then summed across all retained countries to calculate the global annual totals.

**Notes & Limitations**

- Because the data are reported at the national scale, they are not representative of fertilizer use on islands.
- Reporting by countries is sporadic, with a global rate of 33%. Consequently, a large fraction of the data is imputed or estimated.
- Does not include organic fertilizers.
- Limited to agricultural use, excluding industrial and technical applications.

#### Component indicator: CFC Emissions in ODP Tonnes

##### Description

This UNEP indicator tracks the consumption and production of Chlorofluorocarbons (CFCs) measured in ODP Tonnes.

##### Temporal Trends

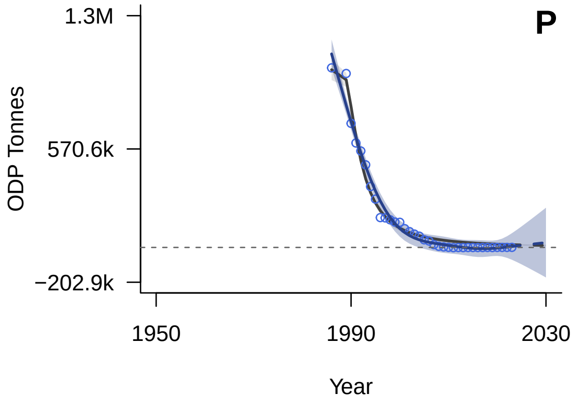


*General Additive Mixed Model (GAMM) fitted trend (solid line) with 95% confidence interval (shaded area). The dashed continuation of the trend line represents the extrapolated future trajectory until 2030. Blue indicates positive tracking towards the target; orange indicates negative tracking. The horizontal grey dashed line marks the 2020 benchmark value. The grey line (solid and dashed) and shading show the model averaging trend and confidence interval.*

##### Spatial Coverage

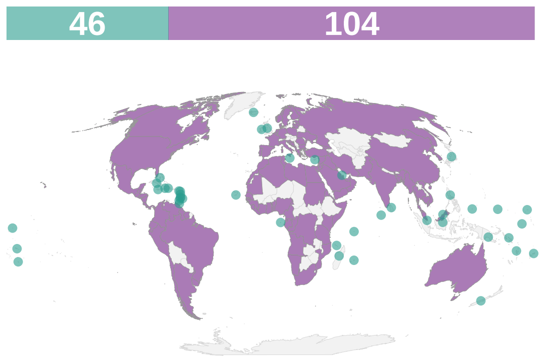


*Spatial coverage of data availability across 150 coastal and island nations. Purple: coastal countries; teal: island countries; grey: countries without data or landlocked. The bars on top of the map indicate the count of countries in each category (purple: coastal countries; teal: island countries; grey: no data).*

##### Key Information

| | |
|---|---|
| **Units** | ODP Tonnes |
| **Time Period** | 1986-2023 |
| **Disaggregation** | Country |
| **Source** | UN, https://ozone.unep.org/countries/data-table |
| **Accession Date** | Jan-25 |
| **DPSIR Framework** | Pressure |
| **Theory of Change (ToC)** | N/A |

##### Methodology

The dataset was already in a clean, ready-to-use format (country × year). Annual consumption values were aggregated by country and year to produce global annual totals.

##### Notes & Limitations

- EU reports aggregate values for the Union. We have included the EU, treated as a coastal country, therefore including a few landlocked european countries.

- Some values are negative for a given year, which implies that quantities destroyed or exported for the year exceeded the sum of production and imports, and that such destroyed or exported quantities came from stockpiles.

#### Component indicator: Red List Index for Birds Showing Trends Driven by Pollution

##### Description

The Red List Index (RLI) shows trends in overall extinction risk for species. This indicator focuses on trends driven by pollution for birds alone.

##### Temporal Trends

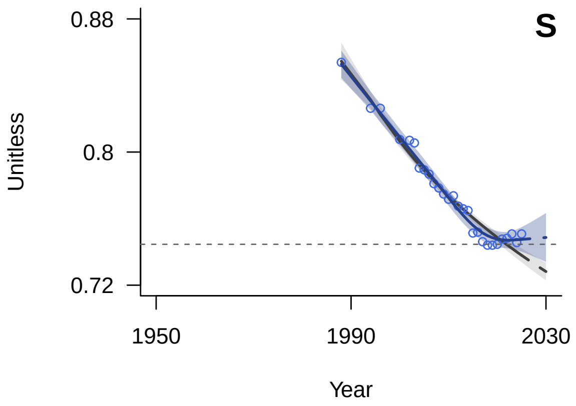


*General Additive Mixed Model (GAMM) fitted trend (solid line) with 95% confidence interval (shaded area). The dashed continuation of the trend line represents the extrapolated future trajectory until 2030. Blue indicates positive tracking towards the target; orange indicates negative tracking. The horizontal grey dashed line marks the 2020 benchmark value. The grey line (solid and dashed) and shading show the model averaging trend and confidence interval.*

##### Spatial Coverage

*No spatial coverage data:*

*The Red List Index is calculated as a global aggregate.*

##### Key Information

| | |
|---|---|
| **Units** | Unitless |
| **Time Period** | 1988-2025 |
| **Disaggregation** | Spatially explicit |
| **Source** | IUCN Red List, https://www.iucnredlist.org |
| **Accession Date** | Feb-26 |
| **DPSIR Framework** | State |
| **Theory of Change (ToC)** | N/A |

##### Methodology

We extracted spatial data for birds from threat category Pollution. Spatial data were intersected with the global island layer (Weigelt et al., 2013) using QGIS 3.34.15 to generate a unique island-species list. This list served as the basis for querying historical species assessments from the IUCN Red List API via the ‘rredlist’ package in R 4.4.2. To calculate the Red List Index (RLI) following the methodology of Butchart et al. (2007), we established a baseline of species consistently assessed since 1988. Data Deficient (DD), Not Recognized (NR), and Not Evaluated (NE) species were strictly excluded. We assigned each remaining species a category weight corresponding to its extinction risk (ranging from 0 for Least Concern to 5 for Extinct). We manually excluded two species that feature non-genuine changes, which were primarily driven by taxonomic splits (i.e., *Puffinus bannermani*, *Sarkidiornis sylvicola*). We applied a Last-Observation-Carried-Forward (LOCF) approach to fill assessment gaps. We focused on global assessments only.

##### Notes & Limitations

- We could not systematically account for non genuine changes: the information cannot be extracted via the current API version (v3).
- This indicator may project an overly optimistic view of global extinction risk because birds are a well-studied, highly protected taxa. Conversely, severe declines in poorly assessed groups (e.g., invertebrates, plants, and fungi) may go undetected.
- The Red List's baseline effectively excludes post-Linnaean extinctions that occurred before systematic assessment began.
- Data Deficient (DD) taxa were excluded, yet these poorly documented taxa often face higher extinction risks than fully assessed species.
- The conservative 50-year rule of thumb could delay the recognition of recent extinctions.

#### Component indicator: Cropland nutrient budget

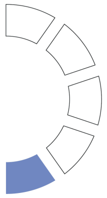

##### Description

This indicator tracks the net balance between total nutrient inputs applied to agricultural soils and total nutrient outputs harvested or lost to the environment.

##### Temporal Trends

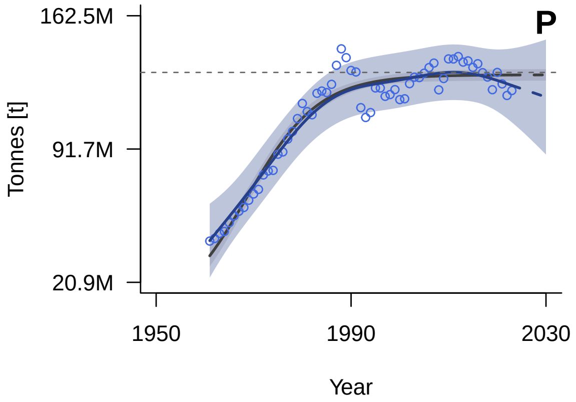


*General Additive Mixed Model (GAMM) fitted trend (solid line) with 95% confidence interval (shaded area). The dashed continuation of the trend line represents the extrapolated future trajectory until 2030. Blue indicates positive tracking towards the target; orange indicates negative tracking. The horizontal grey dashed line marks the 2020 benchmark value. The grey line (solid and dashed) and shading show the model averaging trend and confidence interval.*

##### Spatial Coverage

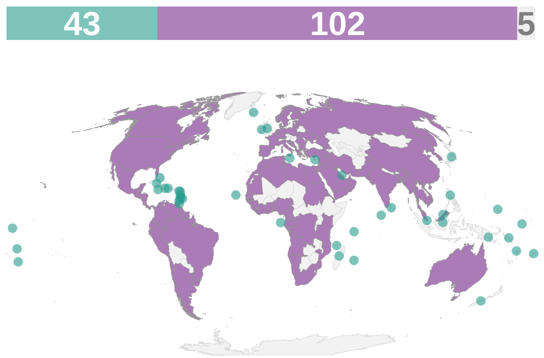


*Spatial coverage of data availability across 150 coastal and island nations. Purple: coastal countries; teal: island countries; grey: countries without data or landlocked. The bars on top of the map indicate the count of countries in each category (purple: coastal countries; teal: island countries; grey: no data).*

##### Key Information

| | |
|---|---|
| **Units** | Tonnes [t] |
| **Time Period** | 1961-2023 |
| **Disaggregation** | Country |
| **Source** | FAO, https://www.fao.org/faostat/en/#data/ESB |
| **Accession Date** | Jul-26 |
| **DPSIR Framework** | Pressure |
| **Theory of Change (ToC)** | N/A |

##### Methodology

The dataset was structured into a country-by-year matrix and summed to calculate the total nutrient balance (nitrogen, phosphorus, and potassium).

##### Notes & Limitations

- Comparability may be limited over time due to breaks in the underlying source data time series.
- Missing values were often imputed by the FAO.

### Target 8

#### Minimize the Impacts of Climate Change on Biodiversity and Build Resilience

Minimize the impact of climate change and ocean acidification on biodiversity and increase its resilience through mitigation, adaptation, and disaster risk reduction actions, including through nature-based solution and/or ecosystem-based approaches, while minimizing negative and fostering positive impacts of climate action on biodiversity.

*No indicators available for this target.*

### Target 9

#### Manage Wild Species Sustainably To Benefit People

Ensure that the management and use of wild species are sustainable, thereby providing social, economic and environmental benefits for people, especially those in vulnerable situations and those most dependent on biodiversity, including through sustainable biodiversity-based activities, products and services that enhance biodiversity, and protecting and encouraging customary sustainable use by indigenous peoples and local communities.

#### Complementary indicator: Spawning stock biomass (related to commercially exploited species)

##### Description

This indicator tracks the total combined weight of sexually mature individuals within commercially harvested fish populations.

##### Temporal Trends

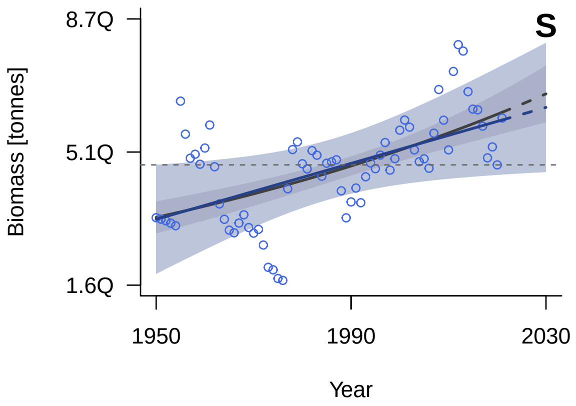


*General Additive Mixed Model (GAMM) fitted trend (solid line) with 95% confidence interval (shaded area). The dashed continuation of the trend line represents the extrapolated future trajectory until 2030. Blue indicates positive tracking towards the target; orange indicates negative tracking. The horizontal grey dashed line marks the 2020 benchmark value. The grey line (solid and dashed) and shading show the model averaging trend and confidence interval.*

##### Spatial Coverage

*No spatial coverage data:*
*Calculated as a global aggregate.*

##### Key Information

| | |
|---|---|
| **Units** | Biomass [tonnes] |
| **Time Period** | 1950-2021 |
| **Disaggregation** | Basins |
| **Source** | RAM Legacy Dataset, https://www.ramlegacy.org/ |
| **Accession Date** | Jul-26 |
| **DPSIR Framework** | State |
| **Theory of Change (ToC)** | N/A |

##### Methodology

The data was provided at a global scale rather than at the country level. The dataset was pre-processed to calculate the global total across all stocks between 1950 and 2021.

##### Notes & Limitations

- The data suffers from a 2-3 year reporting lag. Aggregated global values for the most recent years (i.e., 2022-2024) are incomplete and have been excluded.

#### Target 10

##### Enhance Biodiversity and Sustainability in Agriculture, Aquaculture, Fisheries, and Forestry

Ensure that areas under agriculture, aquaculture, fisheries and forestry are managed sustainably, in particular through the sustainable use of biodiversity, including through a substantial increase of the application of biodiversity friendly practices, such as sustainable intensification, agroecological and other innovative approaches contributing to the resilience and long-term efficiency and productivity of these production systems and to food security, conserving and restoring biodiversity and maintaining nature’s contributions to people, including ecosystem functions and services .

#### Headline indicator: Progress towards Sustainable Forest Management

##### Description

The total forest area managed in compliance with published standards of a forest management certification scheme and independently verified by a third party.

##### Temporal Trends

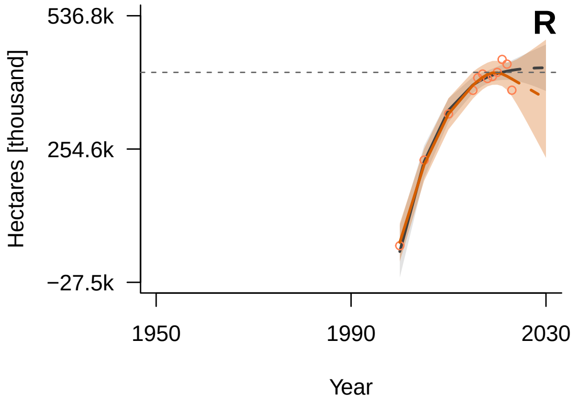


*General Additive Mixed Model (GAMM) fitted trend (solid line) with 95% confidence interval (shaded area). The dashed continuation of the trend line represents the extrapolated future trajectory until 2030. Blue indicates positive tracking towards the target; orange indicates negative tracking. The horizontal grey dashed line marks the 2020 benchmark value. The grey line (solid and dashed) and shading show the model averaging trend and confidence interval.*

##### Spatial Coverage

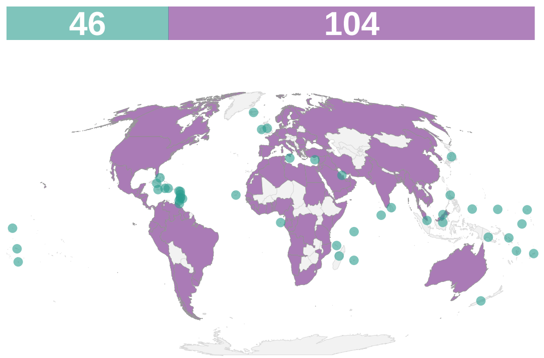


*Spatial coverage of data availability across 150 coastal and island nations. Purple: coastal countries; teal: island countries; grey: countries without data or landlocked. The bars on top of the map indicate the count of countries in each category (purple: coastal countries; teal: island countries; grey: no data).*

##### Key Information

| | |
|---|---|
| **Units** | Hectares [thousand] |
| **Time Period** | 2000-2023 |
| **Disaggregation** | Country |
| **Source** | UN SDGs, https://unstats.un.org/sdgs/dataportal/database |
| **Accession Date** | Jan-25 |
| **DPSIR Framework** | Response |
| **Theory of Change (ToC)** | Output |

##### Methodology

We extracted data on Sustainable Development Goal (SDG) indicator 15.2.1, specifically sub-indicator (e), which measures the forest area under an independently verified forest management certification scheme. This dataset includes certifications from both the Forest Stewardship Council (FSC) and the Programme for the Endorsement of Forest Certification (PEFC) and accounts for double-certified areas. We isolated the geographic area, reported value (in thousand hectares), and time period for each entry.

##### Notes & Limitations

- The removal of the Russian Federation from major international certification schemes in 2023 introduces a sharp artificial decline in the global certified area that does not necessarily reflect an immediate change in on-the-ground ecological conditions.
- The indicator only tracks formally certified forest management, potentially underestimating actual sustainable practices in regions or local communities where obtaining certification is economically or administratively prohibitive.
- Certified forest area serves as an administrative metric of management adherence and does not directly measure or guarantee the actual quality of biodiversity conservation outcomes within those forests.
- Certified forests are not only multiple use forest; there are also certified forest plantations, which could be biodiversity poor and bear limited potential for other forest services.

#### Complementary indicator: Forestry production and trade (wood fuel)

##### Description

This indicator tracks the total physical volume of wood harvested and traded specifically for energy production and direct fuel use.

##### Temporal Trends

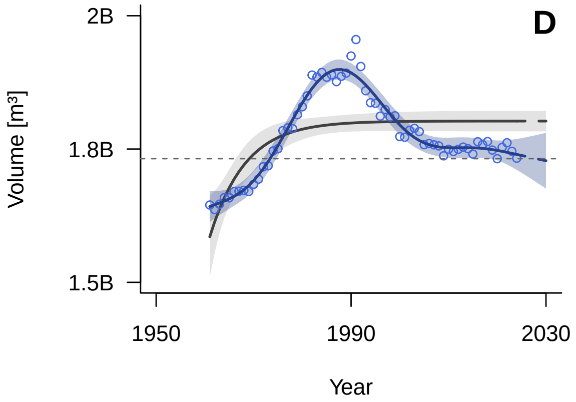


*General Additive Mixed Model (GAMM) fitted trend (solid line) with 95% confidence interval (shaded area). The dashed continuation of the trend line represents the extrapolated future trajectory until 2030. Blue indicates positive tracking towards the target; orange indicates negative tracking. The horizontal grey dashed line marks the 2020 benchmark value. The grey line (solid and dashed) and shading show the model averaging trend and confidence interval.*

##### Spatial Coverage

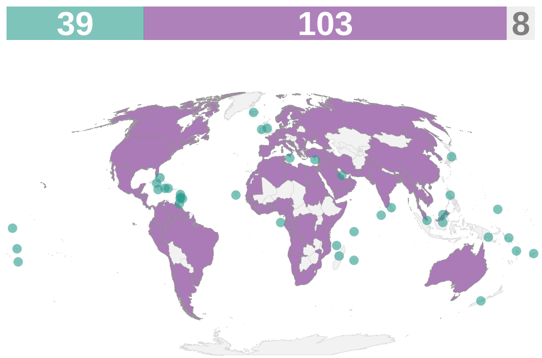


*Spatial coverage of data availability across 150 coastal and island nations. Purple: coastal countries; teal: island countries; grey: countries without data or landlocked. The bars on top of the map indicate the count of countries in each category (purple: coastal countries; teal: island countries; grey: no data).*

##### Key Information

| | |
|---|---|
| **Units** | Volume [m³] |
| **Time Period** | 1961-2024 |
| **Disaggregation** | Country |
| **Source** | FAO, https://www.fao.org/faostat/en/#data/FO |
| **Accession Date** | Jul-26 |
| **DPSIR Framework** | Driver |
| **Theory of Change (ToC)** | N/A |

##### Methodology

The dataset was already in a clean, ready-to-use format (country × year). Data were filtered to extract total wood fuel production by summing coniferous and non-coniferous categories.

##### Notes & Limitations

- Because we are only using total production rather than including imports and exports, this indicator does not reflect true net domestic supply.

- The quality of submitted data varies considerably between countries. Missing values are supplemented with alternative sources or imputed.
- For production data overall, the rate of imputation and estimation is highly variable (between 5% and 80% of global volume depending on the specific forest product).
- Geographical comparability for roundwood (which includes wood fuel) can be limited due to differences in national methods and coverage.

### Target 11

#### Restore, Maintain and Enhance Nature’s Contributions to People

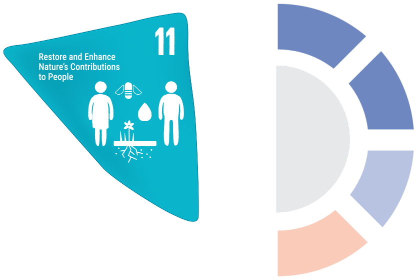


Restore, maintain and enhance nature’s contributions to people, including ecosystem functions and services, such as regulation of air, water, and climate, soil health, pollination and reduction of disease risk, as well as protection from natural hazards and disasters, through nature-based solutions and/or ecosystem-based approaches for the benefit of all people and nature.

#### Component indicator: Annual mean levels of fine particulate matter (e.g. PM2.5 and PM10) in cities

##### Description

This indicator tracks average urban air pollution levels by measuring airborne concentrations of fine inhalable particulate matter under 2.5 and 10 micrometers in diameter.

##### Temporal Trends

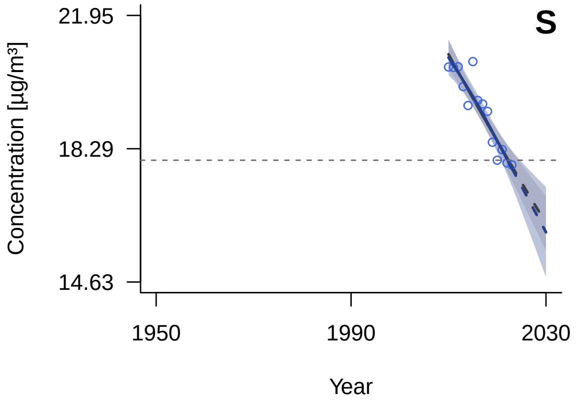


*General Additive Mixed Model (GAMM) fitted trend (solid line) with 95% confidence interval (shaded area). The dashed continuation of the trend line represents the extrapolated future trajectory until 2030. Blue indicates positive tracking towards the target; orange indicates negative tracking. The horizontal grey dashed line marks the 2020 benchmark value. The grey line (solid and dashed) and shading show the model averaging trend and confidence interval.*

##### Spatial Coverage

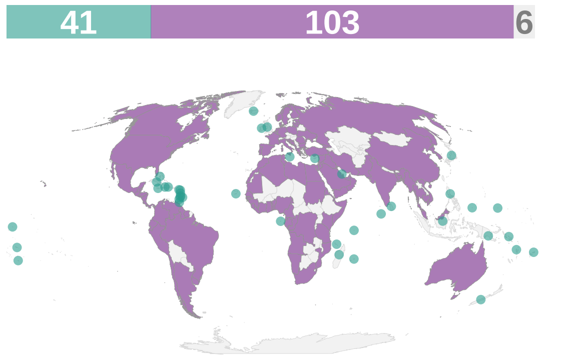


*Spatial coverage of data availability across 150 coastal and island nations. Purple: coastal countries; teal: island countries; grey: countries without data or landlocked. The bars on top of the map indicate the count of countries in each category (purple: coastal countries; teal: island countries; grey: no data).*

##### Key Information

| | |
|---|---|
| **Units** | Concentration [µg/m³] |
| **Time Period** | 2010-2023 |
| **Disaggregation** | Country |
| **Source** | UN SDGs, https://unstats.un.org/sdgs/dataportal/database |
| **Accession Date** | Jul-26 |
| **DPSIR Framework** | State |
| **Theory of Change (ToC)** | N/A |

##### Methodology

The dataset was already in a clean, ready-to-use format (country × year). Data were filtered to extract the overall national fine particulate matter levels.

##### Notes & Limitations

- Data quality varies across countries.

- Data gaps particularly in Small Island Developing States (SIDS) in the Western Pacific and the Latin America and Caribbean regions.

### Component indicator: Level of water stress: freshwater withdrawal as a proportion of available freshwater resources

#### Description

This indicator tracks pressure on renewable freshwater systems by measuring total freshwater extracted relative to total naturally renewable supply.

#### Temporal Trends

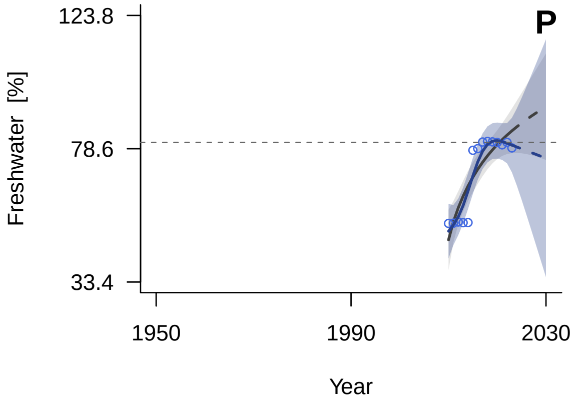


*General Additive Mixed Model (GAMM) fitted trend (solid line) with 95% confidence interval (shaded area). The dashed continuation of the trend line represents the extrapolated future trajectory until 2030. Blue indicates positive tracking towards the target; orange indicates negative tracking. The horizontal grey dashed line marks the 2020 benchmark value. The grey line (solid and dashed) and shading show the model averaging trend and confidence interval.*

#### Spatial Coverage

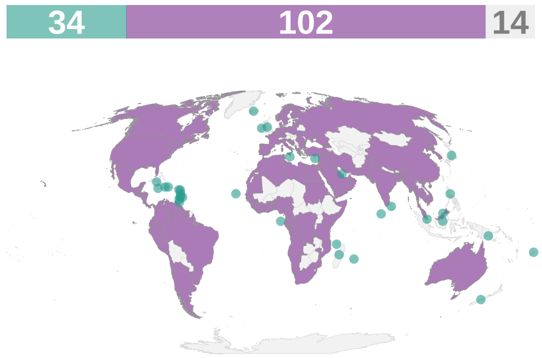


*Spatial coverage of data availability across 150 coastal and island nations. Purple: coastal countries; teal: island countries; grey: countries without data or landlocked. The bars on top of the map indicate the count of countries in each category (purple: coastal countries; teal: island countries; grey: no data).*

#### Key Information

| | |
|---|---|
| **Units** | Freshwater [%] |
| **Time Period** | 2010-2023 |
| **Disaggregation** | Country |
| **Source** | UN SDGs, https://unstats.un.org/sdgs/dataportal/database |
| **Accession Date** | Jul-26 |
| **DPSIR Framework** | Pressure |
| **Theory of Change (ToC)** | N/A |

#### Methodology

The dataset was already in a clean, ready-to-use format (country × year). Data were filtered to extract the overall national level of water stress.

#### Notes & Limitations

N/A

#### Complementary indicator: Proportion of population using safely managed drinking water services

##### Description

This indicator tracks the percentage of the population using an improved drinking water source that is located on premises, available when needed, and free from fecal and priority chemical contamination.

##### Temporal Trends

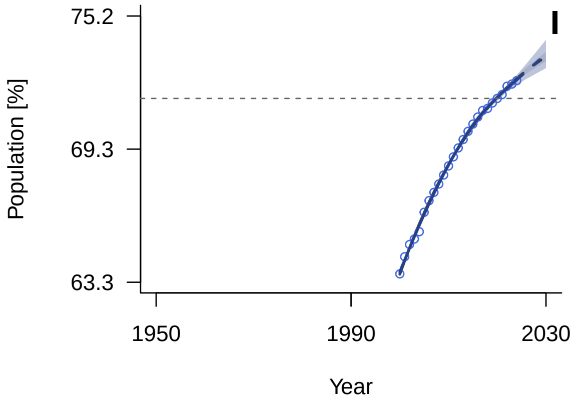


*General Additive Mixed Model (GAMM) fitted trend (solid line) with 95% confidence interval (shaded area). The dashed continuation of the trend line represents the extrapolated future trajectory until 2030. Blue indicates positive tracking towards the target; orange indicates negative tracking. The horizontal grey dashed line marks the 2020 benchmark value. The grey line (solid and dashed) and shading show the model averaging trend and confidence interval.*

##### Spatial Coverage

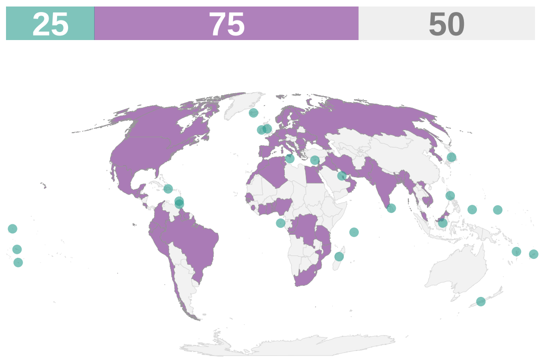


*Spatial coverage of data availability across 150 coastal and island nations. Purple: coastal countries; teal: island countries; grey: countries without data or landlocked. The bars on top of the map indicate the count of countries in each category (purple: coastal countries; teal: island countries; grey: no data).*

##### Key Information

| | |
|---|---|
| **Units** | Population [%] |
| **Time Period** | 2000-2024 |
| **Disaggregation** | Country |
| **Source** | UN SDGs, https://unstats.un.org/sdgs/dataportal/database |
| **Accession Date** | Jul-26 |
| **DPSIR Framework** | Impact |
| **Theory of Change (ToC)** | N/A |

##### Methodology

The dataset was already in a clean, ready-to-use format (country × year). Data were aggregated by country and year.

#### Notes & Limitations

- Definitions regarding the availability and safety of drinking water have yet to be globally standardized across household surveys and administrative sources.

### Complementary indicator: Number of deaths, missing persons and directly affected persons attributed to disasters per 100,000 population

#### Description

This indicator tracks the mortality, disappearance, and direct population disruption caused by natural and human-induced disasters relative to population size.

##### Temporal Trends

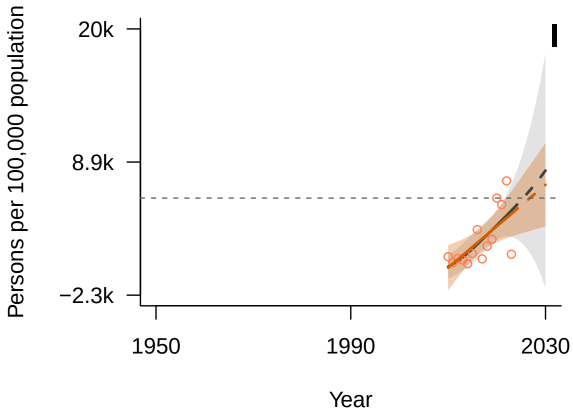


*General Additive Mixed Model (GAMM) fitted trend (solid line) with 95% confidence interval (shaded area). The dashed continuation of the trend line represents the extrapolated future trajectory until 2030. Blue indicates positive tracking towards the target; orange indicates negative tracking. The horizontal grey dashed line marks the 2020 benchmark value. The grey line (solid and dashed) and shading show the model averaging trend and confidence interval.*

##### Spatial Coverage

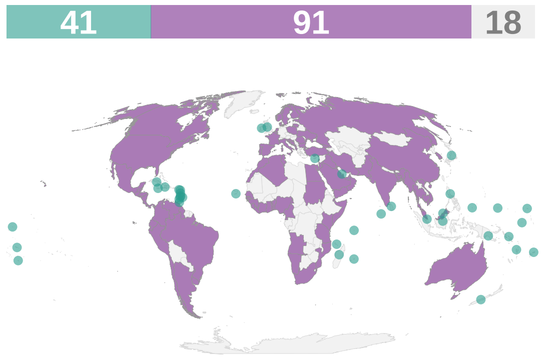


*Spatial coverage of data availability across 150 coastal and island nations. Purple: coastal countries; teal: island countries; grey: countries without data or landlocked. The bars on top of the map indicate the count of countries in each category (purple: coastal countries; teal: island countries; grey: no data).*

#### Key Information

| | |
|---|---|
| **Units** | Persons per 100,000 population |
| **Time Period** | 2010-2023 |
| **Disaggregation** | Country |
| **Source** | UN SDGs, https://unstats.un.org/sdgs/dataportal/database |
| **Accession Date** | Jul-26 |
| **DPSIR Framework** | Impact |
| **Theory of Change (ToC)** | N/A |

#### Methodology

The dataset was already in a clean, ready-to-use format (country × year). Two time series were aggregated: Number of directly affected persons attributed to disasters per 100,000 population and Number of deaths and missing persons attributed to disasters per 100,000 population. Data were then aggregated by country and year.

#### Notes & Limitations

- International data sources often record only events that surpass a specific impact threshold, which can lead to the underestimation of actual values.
- The reliance on secondary data sources with inconsistent methodologies across countries.

### Target 12

#### Enhance Green Spaces and Urban Planning for Human Well-Being and Biodiversity

Significantly increase the area and quality and connectivity of, access to, and benefits from green and blue spaces in urban and densely populated areas sustainably, by mainstreaming the conservation and sustainable use of biodiversity, and ensure biodiversity-inclusive urban planning, enhancing native biodiversity, ecological connectivity and integrity, and improving human health and well-being and connection to nature and contributing to inclusive and sustainable urbanization and the provision of ecosystem functions and services.

*No indicators available for this target.*

#### Target 13

##### Increase the Sharing of Benefits From Genetic Resources, Digital Sequence Information and Traditional Knowledge

Take effective legal, policy, administrative and capacity-building measures at all levels, as appropriate, to ensure the fair and equitable sharing of benefits that arise from the utilization of genetic resources and from digital sequence information on genetic resources, as well as traditional knowledge associated with genetic resources, and facilitating appropriate access to genetic resources, and by 2030 facilitating a significant increase of the benefits shared, in accordance with applicable international access and benefit-sharing instruments.

#### Component indicator: Total number of internationally recognized certificates published on the Access and Benefit-sharing Clearing-House

##### Description

This indicator tracks legally binding documentation published to prove that genetic resources were accessed with prior informed consent and under mutually agreed terms.

##### Temporal Trends

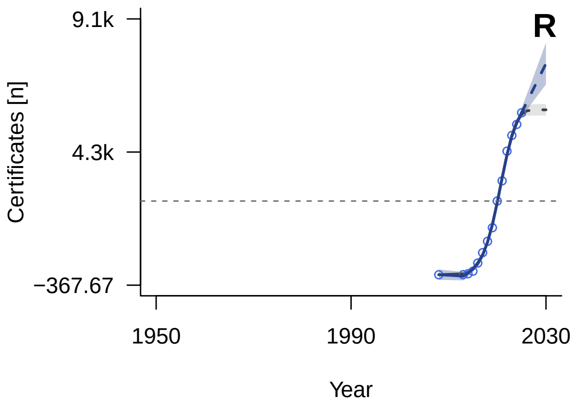


*General Additive Mixed Model (GAMM) fitted trend (solid line) with 95% confidence interval (shaded area). The dashed continuation of the trend line represents the extrapolated future trajectory until 2030. Blue indicates positive tracking towards the target; orange indicates negative tracking. The horizontal grey dashed line marks the 2020 benchmark value. The grey line (solid and dashed) and shading show the model averaging trend and confidence interval.*

##### Spatial Coverage

*No spatial coverage data:*
*Calculated as certificate counts.*

##### Key Information

| | |
|---|---|
| **Units** | Certificates [n] |
| **Time Period** | 2008-2025 |
| **Disaggregation** | Certificates |
| **Source** | CBD ABSCH, https://absch.cbd.int/en/ |
| **Accession Date** | Jul-26 |
| **DPSIR Framework** | Response |
| **Theory of Change (ToC)** | Output |

##### Methodology

The dataset was processed to aggregate individual internationally recognized certificates into a cumulative global count over time.

##### Notes & Limitations

- Only counts certificates issued by countries that are signatories to the Nagoya Protocol.

### Complementary indicator: Number of Countries that Have Ratified the Nagoya Protocol

#### Description

The Nagoya Protocol is a multilateral international agreement which aims at sharing the benefits arising from the utilization of genetic resources in a fair and equitable way.

##### Temporal Trends

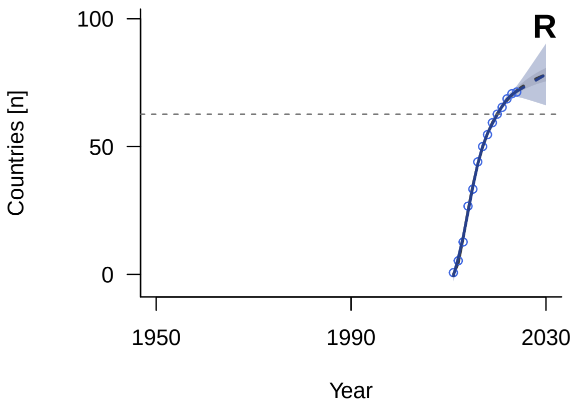


*General Additive Mixed Model (GAMM) fitted trend (solid line) with 95% confidence interval (shaded area). The dashed continuation of the trend line represents the extrapolated future trajectory until 2030. Blue indicates positive tracking towards the target; orange indicates negative tracking. The horizontal grey dashed line marks the 2020 benchmark value. The grey line (solid and dashed) and shading show the model averaging trend and confidence interval.*

##### Spatial Coverage

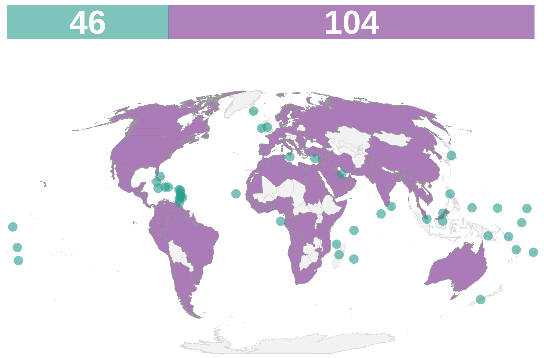


*Spatial coverage of data availability across 150 coastal and island nations. Purple: coastal countries; teal: island countries; grey: countries without data or landlocked. The bars on top of the map indicate the count of countries in each category (purple: coastal countries; teal: island countries; grey: no data).*

#### Key Information

| | |
|---|---|
| **Units** | Countries [n] |
| **Time Period** | 2011-2024 |
| **Disaggregation** | Country |
| **Source** | CBD, https://www.cbd.int/abs/nagoya-protocol/signatories |
| **Accession Date** | Jan-25 |
| **DPSIR Framework** | Response |
| **Theory of Change (ToC)** | Process |

#### Methodology

The Nagoya Protocol ratification list was extracted and organized by year. Values were carried forward from the year of ratification onwards. Global annual totals were calculated by summing the values for each year, then expressed as a proportion.

**Notes & Limitations**

- Ratification is a legal commitment but does not guarantee that the domestic legislative, administrative, or policy measures required by the Protocol are fully functional or enforced at the national level.

#### Complementary indicator: Number of checkpoint communiqués published on the Access and Benefit-sharing Clearing-House

##### Description

This indicator tracks the number of official records published to confirm that utilization of genetic resources was monitored at designated national checkpoints.

##### Temporal Trends

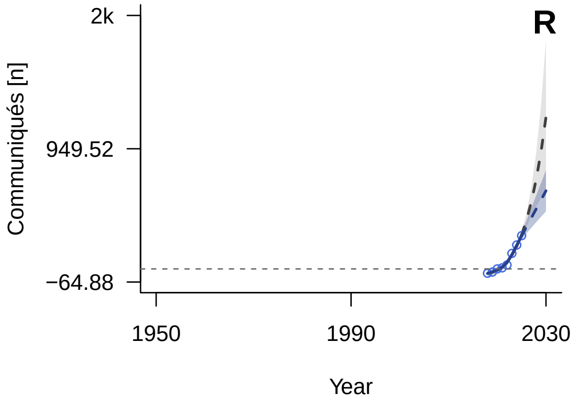


*General Additive Mixed Model (GAMM) fitted trend (solid line) with 95% confidence interval (shaded area). The dashed continuation of the trend line represents the extrapolated future trajectory until 2030. Blue indicates positive tracking towards the target; orange indicates negative tracking. The horizontal grey dashed line marks the 2020 benchmark value. The grey line (solid and dashed) and shading show the model averaging trend and confidence interval.*

##### Spatial Coverage

*No spatial coverage data:*
*Calculated as communique counts.*

##### Key Information

| | |
|---|---|
| **Units** | Communiqués [n] |
| **Time Period** | 2018-2025 |
| **Disaggregation** | Communiqués |
| **Source** | CBD ABSCH, https://absch.cbd.int/en/ |
| **Accession Date** | Jul-26 |
| **DPSIR Framework** | Response |
| **Theory of Change (ToC)** | Output |

##### Methodology

The dataset was processed to aggregate individual checkpoint communiqués into a cumulative global count over time.

##### Notes & Limitations

- When disaggregating data at the national level, it is not possible to definitively distinguish between a true zero (an absence of relevant events) and a missing value (resulting from non-compliance in

reporting).
- Only counts communiqués issued by countries that are signatories to the Nagoya Protocol.

### Target 14

#### Integrate Biodiversity in Decision-Making at Every Level

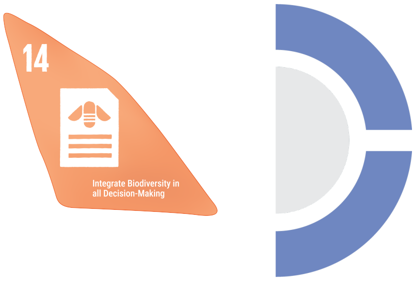


Ensure the full integration of biodiversity and its multiple values into policies, regulations, planning and development processes, poverty eradication strategies, strategic environmental assessments, environmental impact assessments and, as appropriate, national accounting, within and across all levels of government and across all sectors, in particular those with significant impacts on biodiversity, progressively aligning all relevant public and private activities, fiscal and financial flows with the goals and targets of this framework.

#### Component indicator: Number of Countries Implementing Natural Resource Accounts within the System of Environmental-Economic Accounting (SEEA)

##### Description

Recurring UN survey that tracks how many countries have integrated environmental data (such as water and forest use) into their national economic accounts. It provides the data used to monitor SDG indicator 15.9.1(b) regarding the global integration of biodiversity values into national planning.

##### Temporal Trends

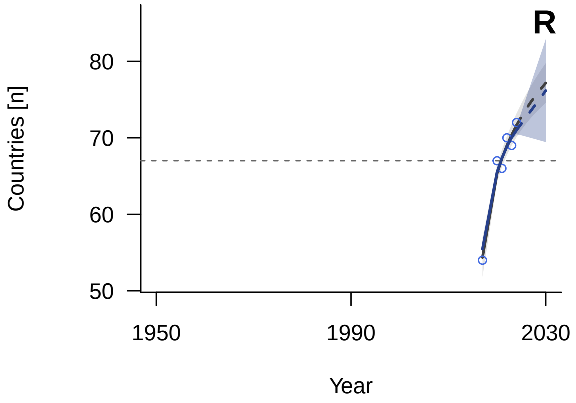


*General Additive Mixed Model (GAMM) fitted trend (solid line) with 95% confidence interval (shaded area). The dashed continuation of the trend line represents the extrapolated future trajectory until 2030. Blue indicates positive tracking towards the target; orange indicates negative tracking. The horizontal grey dashed line marks the 2020 benchmark value. The grey line (solid and dashed) and shading show the model averaging trend and confidence interval.*

##### Spatial Coverage

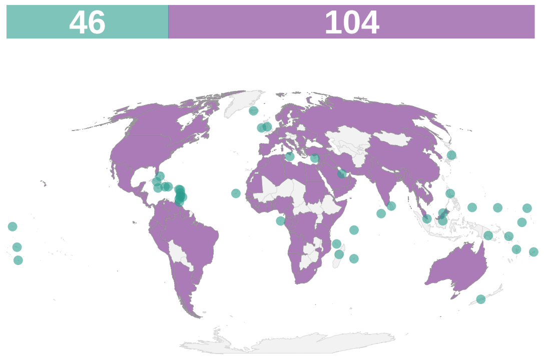


*Spatial coverage of data availability across 150 coastal and island nations. Purple: coastal countries; teal: island countries; grey: countries without data or landlocked. The bars on top of the map indicate the count of countries in each category (purple: coastal countries; teal: island countries; grey: no data).*

##### Key Information

| | |
|---|---|
| **Units** | Countries [n] |
| **Time Period** | 2017-2024 |
| **Disaggregation** | Country |
| **Source** | UN, https://seea.un.org/content/global-assessment-environmental-economic-accounting |
| **Accession Date** | Apr-25 |
| **DPSIR Framework** | Response |
| **Theory of Change (ToC)** | Process |

##### Methodology

The dataset was already in a clean, ready-to-use format (country × year). Annual values were summed across all countries to produce the total number of countries implementing SEEA per year.

##### Notes & Limitations

- A binary status (e.g., 1 or 0) does not reflect the regularity or quality of the environmental-economic accounts compiled by a country.
- The dataset contains missing values for certain countries in specific years. Changes in global aggregates may partially reflect varying response rates rather than actual changes in implementation.

#### Component indicator: Number of NBSAPs Submitted Since 2010

##### Description

The National Biodiversity Strategy and Action Plan (NBSAP) are plans developed by countries outlining strategies that integrate conservation and sustainable use of biological diversity into all relevant sectors of the national economy. It is the primary instrument for the implementation of the Convention on Biological Diversity (CBD) at the national level and are mandatory for participating countries.

##### Temporal Trends

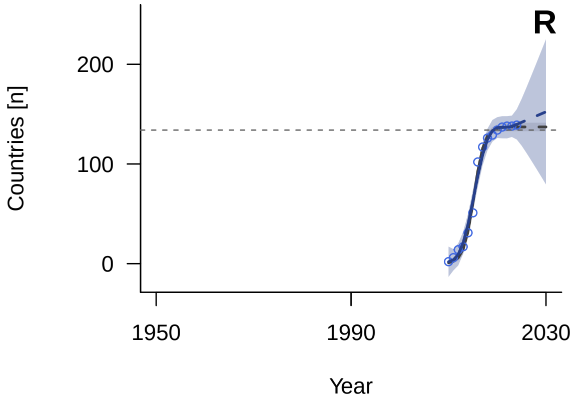


*General Additive Mixed Model (GAMM) fitted trend (solid line) with 95% confidence interval (shaded area). The dashed continuation of the trend line represents the extrapolated future trajectory until 2030. Blue indicates positive tracking towards the target; orange indicates negative tracking. The horizontal grey dashed line marks the 2020 benchmark value. The grey line (solid and dashed) and shading show the model averaging trend and confidence interval.*

##### Spatial Coverage

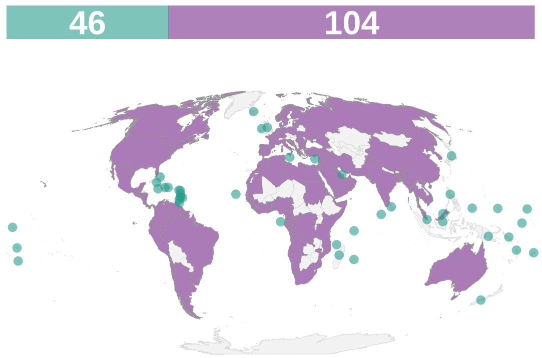


*Spatial coverage of data availability across 150 coastal and island nations. Purple: coastal countries; teal: island countries; grey: countries without data or landlocked. The bars on top of the map indicate the count of countries in each category (purple: coastal countries; teal: island countries; grey: no data).*

##### Key Information

| | |
|---|---|
| **Units** | Countries [n] |
| **Time Period** | 2010-2024 |
| **Disaggregation** | Country |
| **Source** | CBD, https://www.cbd.int/nbsap/introduction.shtml |
| **Accession Date** | Jan-25 |
| **DPSIR Framework** | Response |
| **Theory of Change (ToC)** | Output |

##### Methodology

The NBSAP submission list was extracted and organized by year. Values were carried forward from the year of submission onwards. When multiple years were present, we used the earliest entry. Global annual totals were calculated by summing the values for each year, expressed as a proportion.

##### Notes & Limitations

- This indicator tracks only the submission of the document but does not assess the quality of the strategy or the extent of its actual implementation or funding at the national level.
- The indicator provides an overoptimistic view of current policy relevance because it focuses on older strategies primarily aligned with the 2010 Aichi Targets. While a separate dataset for NBSAPs aligned with the post-2022 Kunming-Montreal Global Biodiversity Framework exists, it was excluded as its short time series did not meet inclusion criteria.

### Target 15

#### Businesses Assess, Disclose and Reduce Biodiversity-Related Risks and Negative Impacts

Take legal, administrative or policy measures to encourage and enable business, and in particular to ensure that large and transnational companies and financial institutions: (a) Regularly monitor, assess, and transparently disclose their risks, dependencies and impacts on biodiversity, including with requirements for all large as well as transnational companies and financial institutions along their operations, supply and value chains and portfolios; (b) Provide information needed to consumers to promote sustainable consumption patterns; (c) Report on compliance with access and benefit-sharing regulations and measures, as applicable; in order to progressively reduce negative impacts on biodiversity, increase positive impacts, reduce biodiversity-related risks to business and financial institutions, and promote actions to ensure sustainable patterns of production.

*No indicators available for this target.*

### Target 16

#### Enable Sustainable Consumption Choices To Reduce Waste and Overconsumption

Ensure that people are encouraged and enabled to make sustainable consumption choices including by establishing supportive policy, legislative or regulatory frameworks, improving education and access to relevant and accurate information and alternatives, and by 2030, reduce the global footprint of consumption in an equitable manner, including through halving global food waste, significantly reducing overconsumption and substantially reducing waste generation, in order for all people to live well in harmony with Mother Earth.

#### Component indicator: Ecological Footprint

##### Description

A measure of how much area of biologically productive land and water an individual, population, or activity requires to produce all the resources it consumes and to absorb the waste it generates, using prevailing technology and resource management practices.

##### Temporal Trends

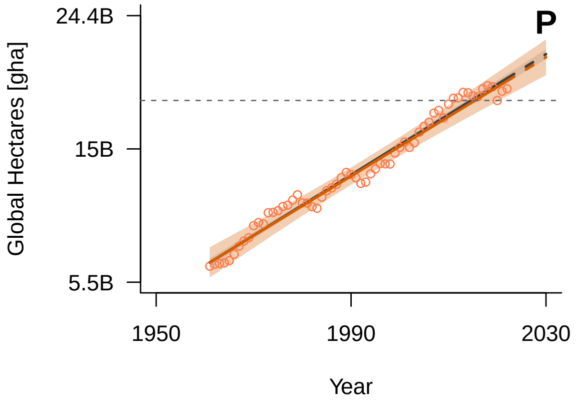


*General Additive Mixed Model (GAMM) fitted trend (solid line) with 95% confidence interval (shaded area). The dashed continuation of the trend line represents the extrapolated future trajectory until 2030. Blue indicates positive tracking towards the target; orange indicates negative tracking. The horizontal grey dashed line marks the 2020 benchmark value. The grey line (solid and dashed) and shading show the model averaging trend and confidence interval.*

##### Spatial Coverage

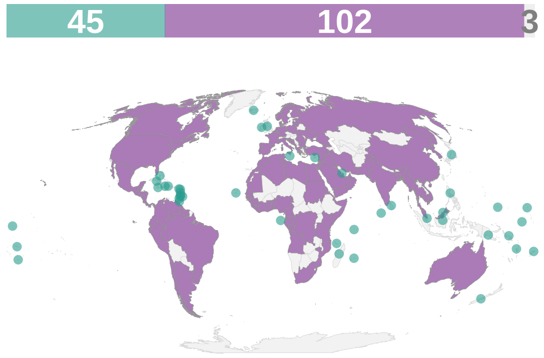


*Spatial coverage of data availability across 150 coastal and island nations. Purple: coastal countries; teal: island countries; grey: countries without data or landlocked. The bars on top of the map indicate the count of countries in each category (purple: coastal countries; teal: island countries; grey: no data).*

##### Key Information

| | |
|---|---|
| **Units** | Global Hectares [gha] |
| **Time Period** | 1961-2022 |
| **Disaggregation** | Country |
| **Source** | Global Footprint Network, https://data.footprintnetwork.org/ |
| **Accession Date** | Jan-25 |
| **DPSIR Framework** | Pressure |
| **Theory of Change (ToC)** | N/A |

##### Methodology

We extracted ecological footprint data from the Global Footprint Network database. We used the total ecological footprint of consumption (EFConsTotGHA), measured in global hectares (gha), which quantifies the area of biologically productive land and water required to produce the resources consumed and absorb the waste generated by each country. Annual values were summed across all retained countries to calculate the global annual totals.

**Notes & Limitations**

N/A

#### Complementary indicator: CO2 emission per unit of value added

##### Description

This indicator tracks the carbon intensity of economic production by measuring carbon dioxide emissions generated relative to total gross value added.

##### Temporal Trends

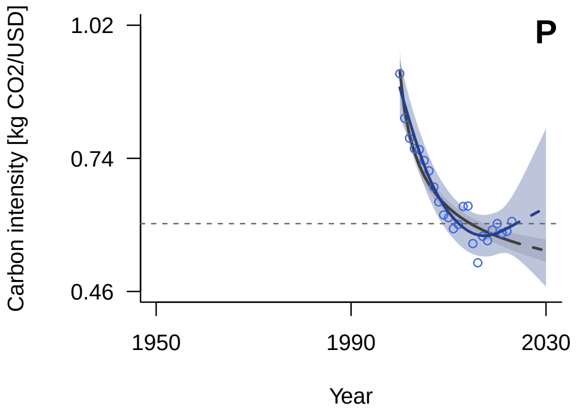


*General Additive Mixed Model (GAMM) fitted trend (solid line) with 95% confidence interval (shaded area). The dashed continuation of the trend line represents the extrapolated future trajectory until 2030. Blue indicates positive tracking towards the target; orange indicates negative tracking. The horizontal grey dashed line marks the 2020 benchmark value. The grey line (solid and dashed) and shading show the model averaging trend and confidence interval.*

##### Spatial Coverage

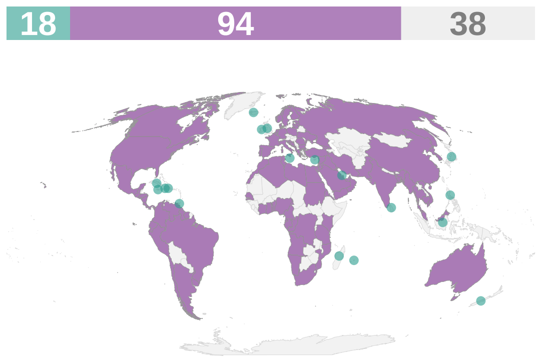


*Spatial coverage of data availability across 150 coastal and island nations. Purple: coastal countries; teal: island countries; grey: countries without data or landlocked. The bars on top of the map indicate the count of countries in each category (purple: coastal countries; teal: island countries; grey: no data).*

##### Key Information

| | |
|---|---|
| **Units** | Carbon intensity [kg $CO_2$/USD] |
| **Time Period** | 2000-2023 |
| **Disaggregation** | Country |
| **Source** | UN SDGs, https://unstats.un.org/sdgs/dataportal/database |
| **Accession Date** | Jul-26 |
| **DPSIR Framework** | Pressure |
| **Theory of Change (ToC)** | N/A |

##### Methodology

The dataset was already in a clean, ready-to-use format (country × year). Data were aggregated by country and year using the arithmetic mean.

##### Notes & Limitations

- Many countries lack standardized and consistent systems for tracking and estimating CO2 emissions.
- National methodologies can sometimes differ from internationally agreed standards.
- Energy consumption data and value-added data are drawn from different sources.

- Discrepancies can arise if countries have not submitted energy consumption data that is adequately disaggregated by sector or energy source, or due to the conversion of value data into USD.
- The metric uses deflated USD (Constant USD).

#### Complementary indicator: Change in water-use efficiency over time

##### Description

This indicator tracks the change in economic value generated per volume of water abstracted over time across major economic sectors.

##### Temporal Trends

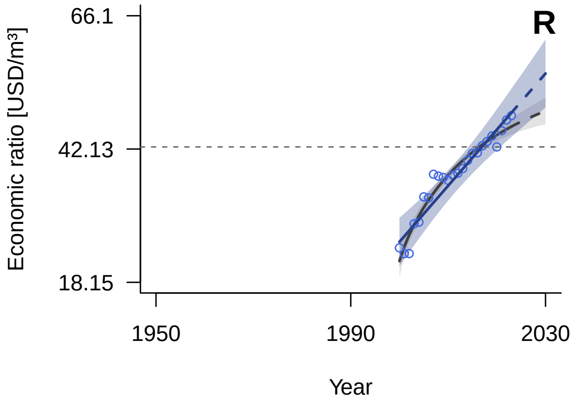


*General Additive Mixed Model (GAMM) fitted trend (solid line) with 95% confidence interval (shaded area). The dashed continuation of the trend line represents the extrapolated future trajectory until 2030. Blue indicates positive tracking towards the target; orange indicates negative tracking. The horizontal grey dashed line marks the 2020 benchmark value. The grey line (solid and dashed) and shading show the model averaging trend and confidence interval.*

##### Spatial Coverage

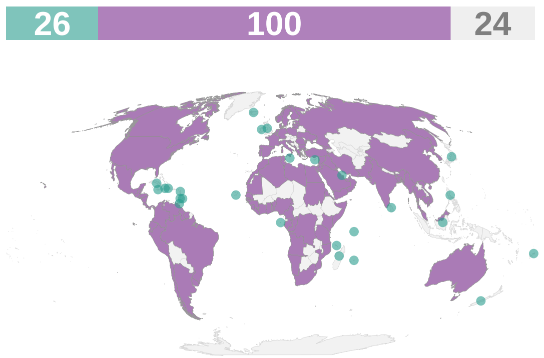


*Spatial coverage of data availability across 150 coastal and island nations. Purple: coastal countries; teal: island countries; grey: countries without data or landlocked. The bars on top of the map indicate the count of countries in each category (purple: coastal countries; teal: island countries; grey: no data).*

##### Key Information

| | |
|---|---|
| **Units** | Economic ratio [USD/m³] |
| **Time Period** | 2000-2023 |
| **Disaggregation** | Country |
| **Source** | UN SDGs, https://unstats.un.org/sdgs/dataportal/database |
| **Accession Date** | Jul-26 |
| **DPSIR Framework** | Response |
| **Theory of Change (ToC)** | Outcome |

##### Methodology

The dataset was already in a clean, ready-to-use format (country × year). Data were aggregated by country and year.

##### Notes & Limitations

- The metric uses deflated USD (Constant USD).

### Target 17

#### Strengthen Biosafety and Distribute the Benefits of Biotechnology

Establish, strengthen capacity for, and implement in all countries biosafety measures as set out in Article 8(g) of the Convention on Biological Diversity and measures for the handling of biotechnology and distribution of its benefits as set out in Article 19 of the Convention.

### Complementary indicator: Number of Countries That Have Ratified the Cartagena Protocol on Biosafety to the Convention on Biological Diversity

#### Description

Multilateral international agreement which aims to ensure the safe handling, transport and use of living modified organisms (LMOs) resulting from modern biotechnology that may have adverse effects on biological diversity, taking also into account risks to human health. It was adopted on 29 January 2000 and entered into force on 11 September 2003.

#### Temporal Trends

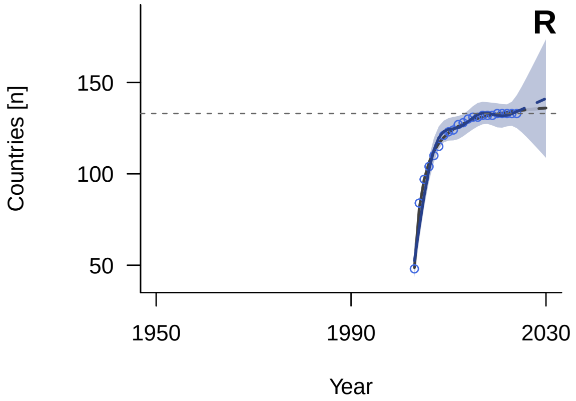


*General Additive Mixed Model (GAMM) fitted trend (solid line) with 95% confidence interval (shaded area). The dashed continuation of the trend line represents the extrapolated future trajectory until 2030. Blue indicates positive tracking towards the target; orange indicates negative tracking. The horizontal grey dashed line marks the 2020 benchmark value. The grey line (solid and dashed) and shading show the model averaging trend and confidence interval.*

#### Spatial Coverage

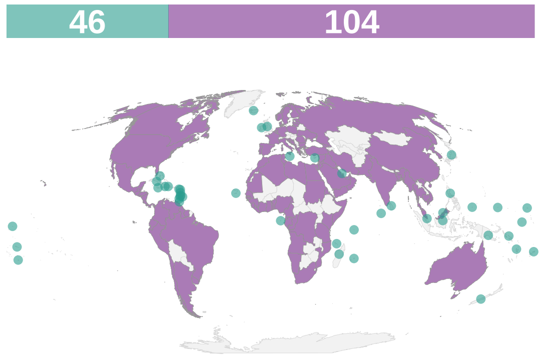


*Spatial coverage of data availability across 150 coastal and island nations. Purple: coastal countries; teal: island countries; grey: countries without data or landlocked. The bars on top of the map indicate the count of countries in each category (purple: coastal countries; teal: island countries; grey: no data).*

#### Key Information

| | |
|---|---|
| **Units** | Countries [n] |
| **Time Period** | 2003-2025 |
| **Disaggregation** | Country |
| **Source** | CBD, https://bch.cbd.int/protocol/parties |
| **Accession Date** | Jan-25 |
| **DPSIR Framework** | Response |
| **Theory of Change (ToC)** | Process |

#### Methodology

The Cartagena Protocol ratification list was extracted and organized by year. Values were carried forward from the year of ratification onwards. Global annual totals were calculated by summing the values for each year.

##### Notes & Limitations

- Ratification indicates a legal commitment but does not ensure that the necessary national biosafety frameworks, monitoring systems, or enforcement mechanisms are active or effective.

#### Target 18

##### Reduce Harmful Incentives by at Least $500 Billion per Year, and Scale Up Positive Incentives for Biodiversity

Identify by 2025, and eliminate, phase out or reform incentives, including subsidies, harmful for biodiversity, in a proportionate, just, fair, effective and equitable way, while substantially and progressively reducing them by at least 500 billion United States dollars per year by 2030, starting with the most harmful incentives, and scale up positive incentives for the conservation and sustainable use of biodiversity.

#### Headline indicator: Positive Incentives in Place to Promote Biodiversity Conservation and Sustainable Use

##### Description

The number of active policy instruments, such as environmental subsidies and grants tracked within the OECD PINE database.

##### Temporal Trends

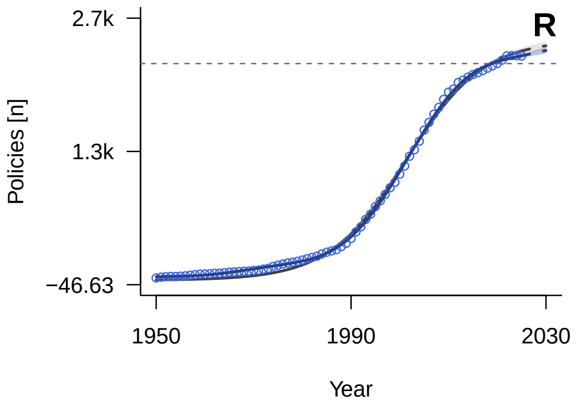


*General Additive Mixed Model (GAMM) fitted trend (solid line) with 95% confidence interval (shaded area). The dashed continuation of the trend line represents the extrapolated future trajectory until 2030. Blue indicates positive tracking towards the target; orange indicates negative tracking. The horizontal grey dashed line marks the 2020 benchmark value. The grey line (solid and dashed) and shading show the model averaging trend and confidence interval.*

##### Spatial Coverage

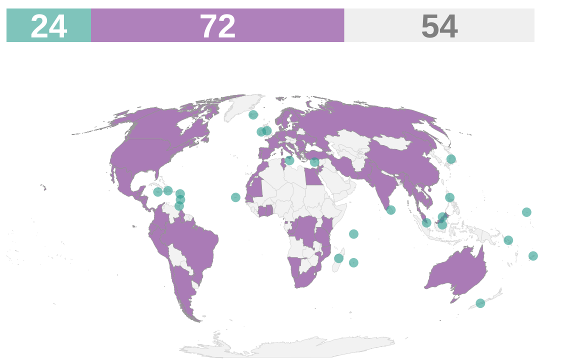


*Spatial coverage of data availability across 150 coastal and island nations. Purple: coastal countries; teal: island countries; grey: countries without data or landlocked. The bars on top of the map indicate the count of countries in each category (purple: coastal countries; teal: island countries; grey: no data).*

##### Key Information

| | |
|---|---|
| **Units** | Policies [n] |
| **Time Period** | 1950-2025 |
| **Disaggregation** | Country |
| **Source** | OECD, https://oecd-main.shinyapps.io/pinedatabase/ |
| **Accession Date** | Jan-25 |
| **DPSIR Framework** | Response |
| **Theory of Change (ToC)** | Input |

##### Methodology

We extracted policy instrument records from the OECD Policy Instruments for the Environment (PINE) database. We excluded inactive policies lacking either a start or end date, as well as active policies without a recorded start date (n = 1,527 records, 33.3%). Each retained policy was then carried forward across its active lifespan, or through to 2025 for policies still in force. The total number of active instruments was then summed by country for each year.

#### Notes & Limitations

- The indicator counts the number of active policy instruments without distinguishing their scope or effectiveness.
- Policies still marked as active are assumed to remain in force through to the present year (2025), which may overestimate the count in case of reporting lags.
- The PINE database coverage is biased towards OECD member countries.

#### Headline indicator: Value of Subsidies and Other Incentives Harmful to Biodiversity

##### Description

This indicator quantifies the total financial value of government subsidies harmful to biodiversity, currently utilizing fossil fuel, agricultur, and fisheries subsidy data.

##### Temporal Trends

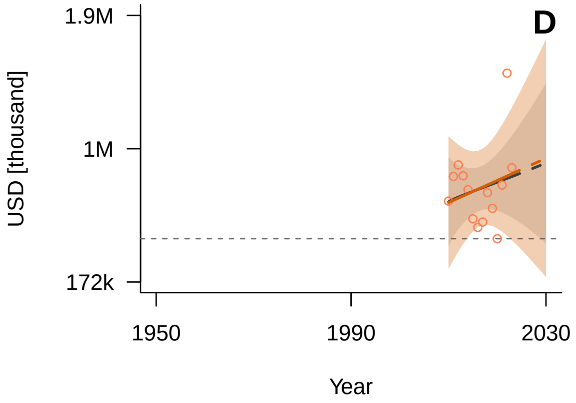


*General Additive Mixed Model (GAMM) fitted trend (solid line) with 95% confidence interval (shaded area). The dashed continuation of the trend line represents the extrapolated future trajectory until 2030. Blue indicates positive tracking towards the target; orange indicates negative tracking. The horizontal grey dashed line marks the 2020 benchmark value. The grey line (solid and dashed) and shading show the model averaging trend and confidence interval.*

##### Spatial Coverage

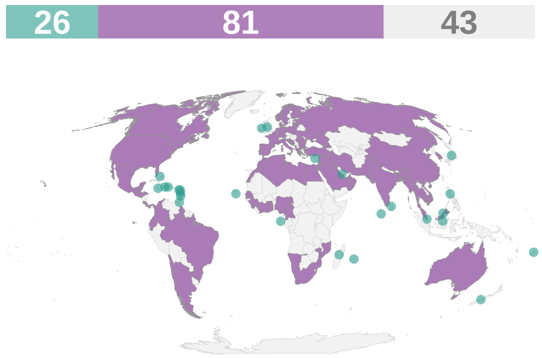


*Spatial coverage of data availability across 150 coastal and island nations. Purple: coastal countries; teal: island countries; grey: countries without data or landlocked. The bars on top of the map indicate the count of countries in each category (purple: coastal countries; teal: island countries; grey: no data).*

##### Key Information

| | |
|---|---|
| **Units** | USD [thousand] |
| **Time Period** | 2010-2023 |
| **Disaggregation** | Country |
| **Source** | UN SDGs, https://unstats.un.org/sdgs/dataportal/database |
| **Accession Date** | Jan-25 |
| **DPSIR Framework** | Driver |
| **Theory of Change (ToC)** | N/A |

##### Methodology

This indicator aims to capture the total value of subsidies harmful to biodiversity across three sectors: fossil fuels, fisheries, and agriculture. However, only fossil fuel subsidy (FFS) data from the UN SDGs database could be included, as fisheries and agricultural subsidies lacked comparable country-level data that could be consistently aggregated. We excluded regional and continental aggregates and retained country-level FFS values for the period 2010–2022, converting from nominal USD billions to thousands of USD. All monetary values were deflated to constant 2024 USD using US GDP deflators from the World Bank.

##### Notes & Limitations

- The indicator captures only fossil fuel subsidies, as agricultural and fisheries subsidies could not be included due to a lack of comparable country-level data. This potentially underestimates the total global value of biodiversity-harmful incentives.
- Monetary values are deflated using a single US-based GDP deflator instead of country-specific price indices. This may misrepresent the actual historical value of subsidies in non-US economies by ignoring local inflation and exchange rate fluctuations.

#### Target 19

##### Mobilize $200 Billion per Year for Biodiversity From all Sources, Including $30 Billion Through International Finance

Substantially and progressively increase the level of financial resources from all sources, in an effective, timely and easily accessible manner, including domestic, international, public and private resources, in accordance with Article 20 of the Convention, to implement national biodiversity strategies and action plans, by 2030 mobilizing at least 200 billion United States dollars per year, including by: (a) Increasing total biodiversity related international financial resources from developed countries, including official development assistance, and from countries that voluntarily assume obligations of developed country Parties, to developing countries, in particular the least developed countries and small island developing States, as well as countries with economies in transition, to at least US$ 20 billion per year by 2025, and to at least US$ 30 billion per year by 2030; (b) Significantly increasing domestic resource mobilization, facilitated by the preparation and implementation of national biodiversity finance plans or similar instruments according to national needs, priorities and circumstances; (c) Leveraging private finance, promoting blended finance, implementing strategies for raising new and additional resources, and encouraging the private sector to invest in biodiversity, including through impact funds and other instruments; (d) Stimulating innovative schemes such as payment for ecosystem services, green bonds, biodiversity offsets and credits, benefit-sharing mechanisms, with environmental and social safeguards; (e) Optimizing co-benefits and synergies of finance targeting the biodiversity and climate crises; (f) Enhancing the role of collective actions, including by indigenous peoples and local communities, Mother Earth centric actions[1] and non-market-based approaches including community based natural resource management and civil society cooperation and solidarity aimed at the conservation of biodiversity; (g) Enhancing the effectiveness, efficiency and transparency of resource provision and use.

#### Headline indicator: International Public Funding, Including Official Development Assistance (ODA) for Conservation and Sustainable Use of Biodiversity and Ecosystems

##### Description

This indicator measures total international public finance disbursements to developing countries that specifically target the conservation and sustainable use of biodiversity and ecosystems.

##### Temporal Trends

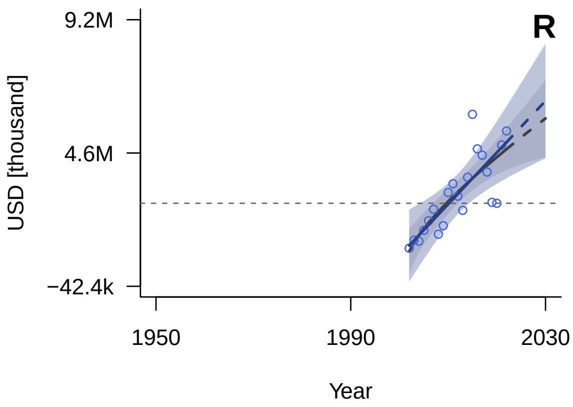


*General Additive Mixed Model (GAMM) fitted trend (solid line) with 95% confidence interval (shaded area). The dashed continuation of the trend line represents the extrapolated future trajectory until 2030. Blue indicates positive tracking towards the target; orange indicates negative tracking. The horizontal grey dashed line marks the 2020 benchmark value. The grey line (solid and dashed) and shading show the model averaging trend and confidence interval.*

##### Spatial Coverage

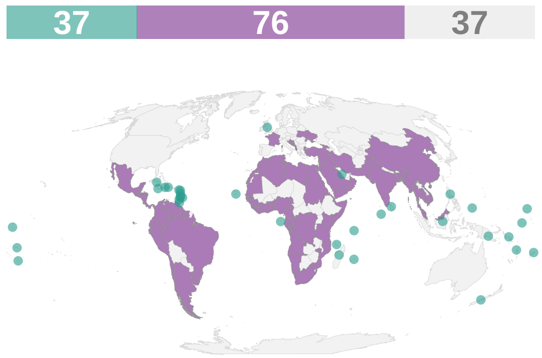


*Spatial coverage of data availability across 150 coastal and island nations. Purple: coastal countries; teal: island countries; grey: countries without data or landlocked. The bars on top of the map indicate the count of countries in each category (purple: coastal countries; teal: island countries; grey: no data).*

##### Key Information

| | |
|---|---|
| **Units** | USD [thousand] |
| **Time Period** | 2002-2023 |
| **Disaggregation** | Country |
| **Source** | OECD, https://data-explorer.oecd.org |
| **Accession Date** | Feb-25 |
| **DPSIR Framework** | Response |
| **Theory of Change (ToC)** | Input |

##### Methodology

We extracted biodiversity-related Official Development Assistance (ODA) data from the OECD Rio Markers database. We retained only biodiversity-flagged funds. Regional and continental aggregates and entries flagged as "unspecified" were excluded. We also extracted TOSSD (Total Official Support for Sustainable Development) data, but this source was ultimately excluded due to insufficient temporal coverage (only 4 data points). The year 2023 was excluded due to a reporting lag at the time of data extraction.

**Notes & Limitations**

- The Rio Markers database covers ODA flows from OECD Development Assistance Committee (DAC) members only, and therefore does not capture South-South cooperation or private sector biodiversity finance.
- The most recent year (2023) was excluded due to a reporting lag; recent totals may therefore be either under- or overestimated.
- TOSSD data were excluded due to insufficient temporal coverage (only 4 data points).

#### Component indicator: Funding Provided by the Global Environment Facility

##### Description

This indicator measures the total financial resources allocated and disbursed by the GEF for projects specifically targeting the protection, restoration, and sustainable management of global biodiversity and ecosystems.

##### Temporal Trends

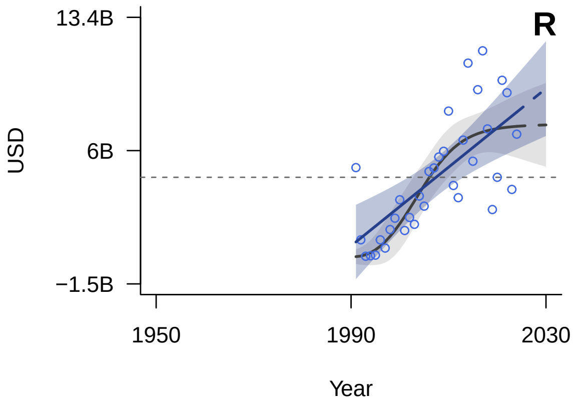


*General Additive Mixed Model (GAMM) fitted trend (solid line) with 95% confidence interval (shaded area). The dashed continuation of the trend line represents the extrapolated future trajectory until 2030. Blue indicates positive tracking towards the target; orange indicates negative tracking. The horizontal grey dashed line marks the 2020 benchmark value. The grey line (solid and dashed) and shading show the model averaging trend and confidence interval.*

##### Spatial Coverage

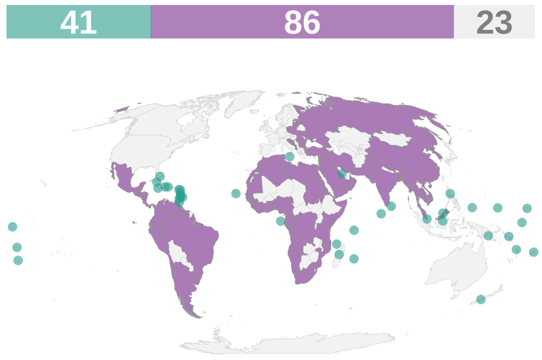


*Spatial coverage of data availability across 150 coastal and island nations. Purple: coastal countries; teal: island countries; grey: countries without data or landlocked. The bars on top of the map indicate the count of countries in each category (purple: coastal countries; teal: island countries; grey: no data).*

##### Key Information

| | |
|---|---|
| **Units** | USD |
| **Time Period** | 1991-2024 |
| **Disaggregation** | Country |
| **Source** | GEF, https://www.thegef.org/projects-operations/database |
| **Accession Date** | Jan-25 |
| **DPSIR Framework** | Response |
| **Theory of Change (ToC)** | Input |

##### Methodology

We extracted biodiversity funding data from the Global Environment Facility (GEF) database. For each project, we summed the GEF Grant and co-financing amounts to represent the total financial resources mobilized. We included only approved projects (completed = 3,407, project approved = 2,175, concept approved = 491) and excluded cancelled (222), received by GEF secretariat (18), and concept proposed (2) statuses. For multi-country projects, the total funding was split equally among all listed recipient countries (removing regional and global tags). All records linked to the placeholder year 1970 were removed. Values were deflated to constant 2024 USD using World Bank GDP deflators, and summed across all retained countries.

**Notes & Limitations**

- It is unclear if the source data are reported in nominal or constant USD; we assumed they are nominal and deflated them to constant 2024 USD.
- For multi-country projects, funding was split equally among recipients, which may not reflect actual national allocations.

### Target 20

#### Strengthen Capacity-Building, Technology Transfer, and Scientific and Technical Cooperation for Biodiversity

Strengthen capacity-building and development, access to and transfer of technology, and promote development of and access to innovation and technical and scientific cooperation, including through South-South, North-South and triangular cooperation, to meet the needs for effective implementation, particularly in developing countries, fostering joint technology development and joint scientific research programmes for the conservation and sustainable use of biodiversity and strengthening scientific research and monitoring capacities, commensurate with the ambition of the goals and targets of the framework.

#### Complementary indicator: Global imports of information and communications technology goods as presented by bilateral trade flows by information and communications technology goods category

##### Description

This indicator tracks international trade volumes and bilateral flows of information and communications technology products.

##### Temporal Trends

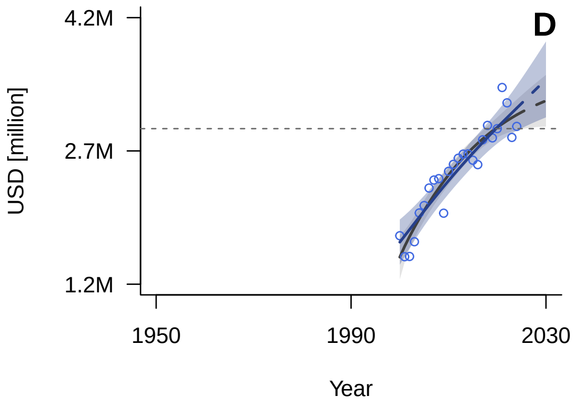


*General Additive Mixed Model (GAMM) fitted trend (solid line) with 95% confidence interval (shaded area). The dashed continuation of the trend line represents the extrapolated future trajectory until 2030. Blue indicates positive tracking towards the target; orange indicates negative tracking. The horizontal grey dashed line marks the 2020 benchmark value. The grey line (solid and dashed) and shading show the model averaging trend and confidence interval.*

##### Spatial Coverage

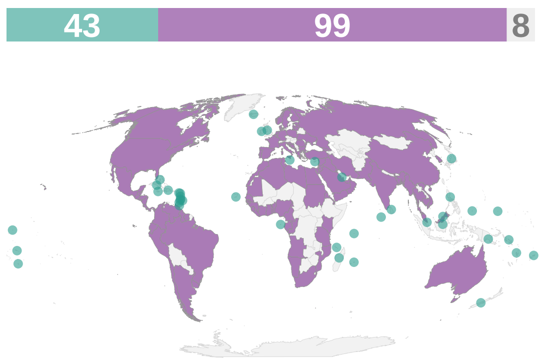


*Spatial coverage of data availability across 150 coastal and island nations. Purple: coastal countries; teal: island countries; grey: countries without data or landlocked. The bars on top of the map indicate the count of countries in each category (purple: coastal countries; teal: island countries; grey: no data).*

##### Key Information

| | |
|---|---|
| **Units** | USD [million] |
| **Time Period** | 2000-2024 |
| **Disaggregation** | Country |
| **Source** | UN Trade Data Hub, https://unctadstat.unctad.org/EN/Index.html |
| **Accession Date** | Jul-26 |
| **DPSIR Framework** | Driver |
| **Theory of Change (ToC)** | N/A |

##### Methodology

The dataset was aggregated by country and year to track the total value of information and communications technology (ICT) goods imported globally (US$ at current prices in millions).

##### Notes & Limitations

- Product details are coded according to varying editions of the Harmonized System (HS) (ranging from 1992 to 2022) depending on the country and year. Because this classification framework changes over time and differs across countries, it can introduce inconsistencies in the data.
- For countries with missing data in specific years, values are estimated to calculate regional aggregates.

#### Target 21

##### Ensure That Knowledge Is Available and Accessible To Guide Biodiversity Action

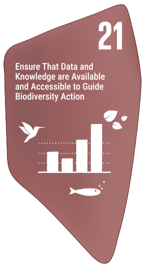


Ensure that the best available data, information and knowledge, are accessible to decision makers, practitioners and the public to guide effective and equitable governance, integrated and participatory management of biodiversity, and to strengthen communication, awareness-raising, education, monitoring, research and knowledge management and, also in this context, traditional knowledge, innovations, practices and technologies of indigenous peoples and local communities should only be accessed with their free, prior and informed consent[1], in accordance with national legislation.

#### Complementary indicator: Knowledge Transfer (Number of Biodiversity Papers in Web of Science per Year)

##### Description

This indicator measures the annual volume of global scientific research focused on island biodiversity.

##### Temporal Trends

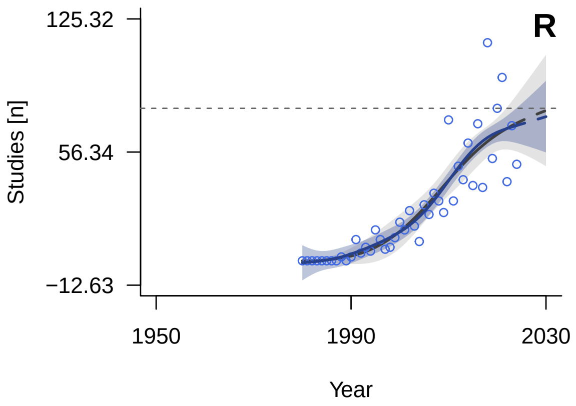


*General Additive Mixed Model (GAMM) fitted trend (solid line) with 95% confidence interval (shaded area). The dashed continuation of the trend line represents the extrapolated future trajectory until 2030. Blue indicates positive tracking towards the target; orange indicates negative tracking. The horizontal grey dashed line marks the 2020 benchmark value. The grey line (solid and dashed) and shading show the model averaging trend and confidence interval.*

##### Spatial Coverage

*No spatial coverage data:*
*Calculated as a global count of published papers.*

##### Key Information

| | |
|---|---|
| **Units** | Studies [n] |
| **Time Period** | 1987-2024 |
| **Disaggregation** | Paper |
| **Source** | WOS, https://www.webofscience.com |
| **Accession Date** | Jan-25 |
| **DPSIR Framework** | Response |
| **Theory of Change (ToC)** | Output |

##### Methodology

We accessed Web Of Science (WOS) and selected "*all datasets*", "*Title*" instead of topic, and typed the keywords (biodiversity AND island*). Then, we selected "*Add data range*" -> "*Publication Dates*" -> "*Custom range*", and added the data range. In this case, from 1980 to today, for consistency with Tittensor et al. 2014. We selected "*Analyse results*", then "*Research area*" and selected "*Publication year*" instead. We scrolled down, selected "*All data rows (up to 100,000)*" and "*Download data table*". We then aggregated data by year. The year 2025 was excluded due to inclomplete data at time of data extraction.

##### Notes & Limitations

- Web of Science is not exhaustive and inherently biases toward English-language, high-impact journals.
- Restricting the query exclusively to the "Title" field (rather than "Topic") yields a highly conservative estimate, omitting relevant papers that only mention biodiversity and islands in their abstracts or keywords.
- The search query (`biodiversity AND island*`) fails to capture island research referring to specific geographic locations without explicitly using the word "island" in the title.

### Target 22

#### Ensure Participation in Decision-Making and Access to Justice and Information Related to Biodiversity for all

Ensure the full, equitable, inclusive, effective and gender-responsive representation and participation in decision-making, and access to justice and information related to biodiversity by indigenous peoples and local communities, respecting their cultures and their rights over lands, territories, resources, and traditional knowledge, as well as by women and girls, children and youth, and persons with disabilities and ensure the full protection of environmental human rights defenders.

*No indicators available for this target.*

#### Target 23

##### Ensure Gender Equality and a Gender-Responsive Approach for Biodiversity Action

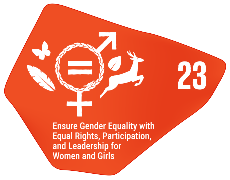


Ensure gender equality in the implementation of the framework through a gender-responsive approach where all women and girls have equal opportunity and capacity to contribute to the three objectives of the Convention, including by recognizing their equal rights and access to land and natural resources and their full, equitable, meaningful and informed participation and leadership at all levels of action, engagement, policy and decision-making related to biodiversity.

#### Component indicator: Proportion of seats held by women in (a) national parliaments and (b) local governments

##### Description

This World Bank indicator measures the percentage of parliamentary seats held by women in a single or lower chamber of national parliaments. It serves as a key metric for Sustainable Development Goal 5.5.1, assessing gender parity and women's access to political decision-making processes globally.

##### Temporal Trends

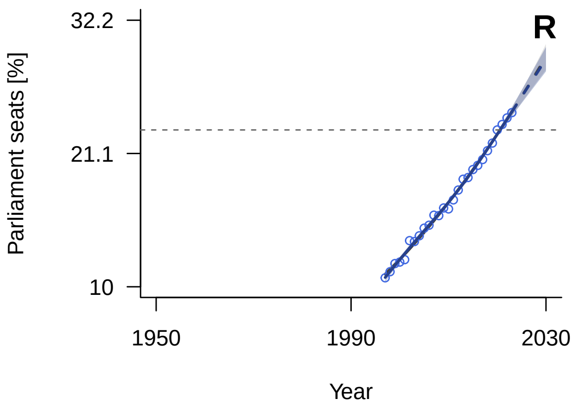


*General Additive Mixed Model (GAMM) fitted trend (solid line) with 95% confidence interval (shaded area). The dashed continuation of the trend line represents the extrapolated future trajectory until 2030. Blue indicates positive tracking towards the target; orange indicates negative tracking. The horizontal grey dashed line marks the 2020 benchmark value. The grey line (solid and dashed) and shading show the model averaging trend and confidence interval.*

##### Spatial Coverage

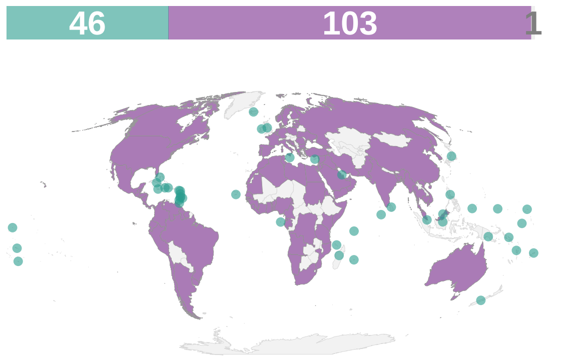


*Spatial coverage of data availability across 150 coastal and island nations. Purple: coastal countries; teal: island countries; grey: countries without data or landlocked. The bars on top of the map indicate the count of countries in each category (purple: coastal countries; teal: island countries; grey: no data).*

##### Key Information

| | |
|---|---|
| **Units** | Parliament seats [%] |
| **Time Period** | 1997-2023 |
| **Disaggregation** | Country |
| **Source** | World Bank, https://data.worldbank.org/indicator/SG.GEN.PARL.ZS |
| **Accession Date** | Jan-25 |
| **DPSIR Framework** | Response |
| **Theory of Change (ToC)** | Process |

##### Methodology

Data for the proportion of seats held by women in national parliaments were extracted from the World Bank World Development Indicators. We calculated annual mean values by country.

##### Notes & Limitations

- It only accounts for national-level parliaments (single or lower house) and does not reflect female representation in local governments, sub-national legislatures, or executive branches.
- The data reflects the proportion of seats held, which does not necessarily translate into the actual influence, leadership positions, or decision-making power held by women within those bodies.
